# Blood Biochemical Responses to Acute Exercise: Findings from the Molecular Transducers of Physical Activity Consortium (MoTrPAC)

**DOI:** 10.64898/2026.03.02.704798

**Authors:** Jeremy M. Robbins, Daniel H. Katz, Gina M. Many, Prashant Rao, Gregory R. Smith, Gaurav Tiwari, Christopher Jin, Guillaume Spielmann, Samuel Montalvo, Gayatri Iyer, David Amar, Damon Leach, Brian J. Coyne, Malene E. Lindholm, Mark A. Sarzynski, Bret H. Goodpaster, Martin J. Walsh, Clary B. Clish, Charles F. Burant, Robert E. Gerszten, MoTrPAC Study Group

## Abstract

Exercise benefits numerous organ systems and tissues, including those not directly involved in locomotor activity. However, the molecular responses to exercise are only partially understood, and identifying circulating biochemical changes may provide further insights into how exercise confers systemic health benefits. Here, we perform large-scale plasma proteomic, metabolomic, and whole blood transcriptional profiling in sedentary human participants undergoing acute endurance exercise, resistance exercise, or a non-exercise control in up to 7 timepoints over a 24-hour period. We identify shared biochemical responses between exercise modes, but also numerous differences in immune responses, lipid metabolism, and pathways reflective of tissue repair and angiogenesis. Integrative analyses of blood and muscle identify differential molecular responses that reflect baseline variation in cardiorespiratory fitness. Taken together, our findings highlight novel temporal and exercise mode-specific, blood-based molecular responses to acute exercise and provide a new resource for the scientific community.

## Introduction

Exercise’s health benefits are uniquely extensive, leading to improvements in cardiorespiratory fitness (CRF), metabolism, muscle strength, mood and cognition, chronic inflammation, healthy aging, and a lower risk of chronic diseases including cancer, obesity, diabetes and cardiovascular disease.^1–7^ It is thus unsurprising that exercise is among the most robust physiologic stimuli, affecting a network of cellular processes and organ systems spanning the cardiovascular, pulmonary, musculoskeletal, neurologic, and endocrine systems among others.^8,9^ While exercise is widely regarded as beneficial, its effects are heterogeneous and vary according to both its exposure (i.e., mode, duration, and intensity) as well as host factors such as age, training background, nutritional status, and the presence or absence of underlying pathologic conditions. Furthermore, there is increasing recognition that genetic and genomic heterogeneity influence inter-individual differences in the response to exercise.^10,11^ Despite this, our understanding of the biochemical pathways that underlie the physiologic effects of exercise and exercise-response traits is incomplete, providing strong rationale to investigate its molecular transducers, a principal goal of the Molecular Transducers of Physical Activity Consortium (MoTrPAC).^12^

Blood provides a readily accessible window into the systemic response to exercise and the molecular signals that coordinate communication across organs. Exercise induces widespread molecular changes throughout the body, including tissues not directly involved in locomotor activity.^9^ These concepts are underscored by the emerging paradigm of “exerkines”, or exercise-stimulated soluble biochemicals that can act in autocrine, paracrine, or endocrine fashion to confer exercise’s health benefits.^13,14^ Candidate exerkines now include small molecule metabolites, lipids, peptides and nucleic acids in the plasma.^15–24^ However, for most of the molecules, their temporal changes during acute exercise, their responses to distinct exercise modes, and their tissue source(s) remain poorly defined. Moreover, the salutary effects of exercise may arise not only from the induction of beneficial signals, but also through the suppression or enhanced clearance of deleterious circulating factors, an area that has received comparatively little attention.^25^

Beyond providing insight into inter-organ signaling, molecular profiling in the blood may identify measures of exercise responsiveness and physiological health. Prior works by our group and others have shown that blood-based signatures can identify differential health responses to an exercise intervention,^26^ reflect CRF,^27,28^ and predict long-term human health outcomes.^29^ Yet previous studies have examined either a single molecular layer, a limited set of analytes, one timepoint and/or exercise mode, or have lacked a non-exercise control arm.^28,30–34^ These limitations motivate a more comprehensive, controlled, multi-omic analysis of the circulating response to exercise.

Here, we describe large-scale plasma proteomic, metabolomic, and whole-blood transcriptomic profiling in well-phenotyped sedentary adults from the MoTrPAC Study in order to systematically characterize the circulating multi-omic responses to acute exercise. We compare responses to acute endurance exercise (EE) and resistance exercise (RE) at up to seven timepoints over the course of 24 hours using a non-exercise control (CON) group to rigorously isolate the effects of exercise from those driven by time of day, fasting, and biospecimen collection. We further integrate blood-based and tissue biochemical features across cardiorespiratory and muscle strength traits to identify shared molecular pathways that underlie exercise phenotypes.

## Results

### Research design and clinical characteristics

A total of 175 sedentary adult participants underwent baseline clinical phenotyping and blood biochemical profiling during and after an acute bout of endurance (EE) or resistance exercise (RE), or no exercise (CON). Participants were enrolled before MoTrPAC was suspended due to the SARS-CoV-2 pandemic and the enrollment procedure is described in detail elsewhere.^35^ The study design, including group assignment, exercise familiarization protocols, clinical phenotyping, biospecimen collection, and acute exercise protocols has been published.^36^ Briefly, healthy but sedentary participants were randomized to EE, RE, or CON arms. EE consisted of a 40-minute bout of cycle ergometry at a power output estimated to elicit ∼65% VO_2_peak. RE consisted of eight upper and lower body exercises, each for three sets of 8-12 repetitions to volitional fatigue. CON participants did not perform exercise but followed the same biospecimen collection protocol as the EE group. Before the acute bouts, EE and RE completed 2 and 3 exercise familiarization sessions, respectively, to establish the appropriate exercise workloads for each participant. The blood biospecimen sampling schedule is shown in **Figure 1A**. Skeletal muscle and adipose tissue were also collected and presented separately (MoTrPAC Clinical and Multi-Omic Landscape papers in submission to *Nature Medicine*; MoTrPAC Skeletal Muscle and Adipose Tissue Companion papers in submission to *Nature Metabolism*).^35,37–39^

**Figure 1.**
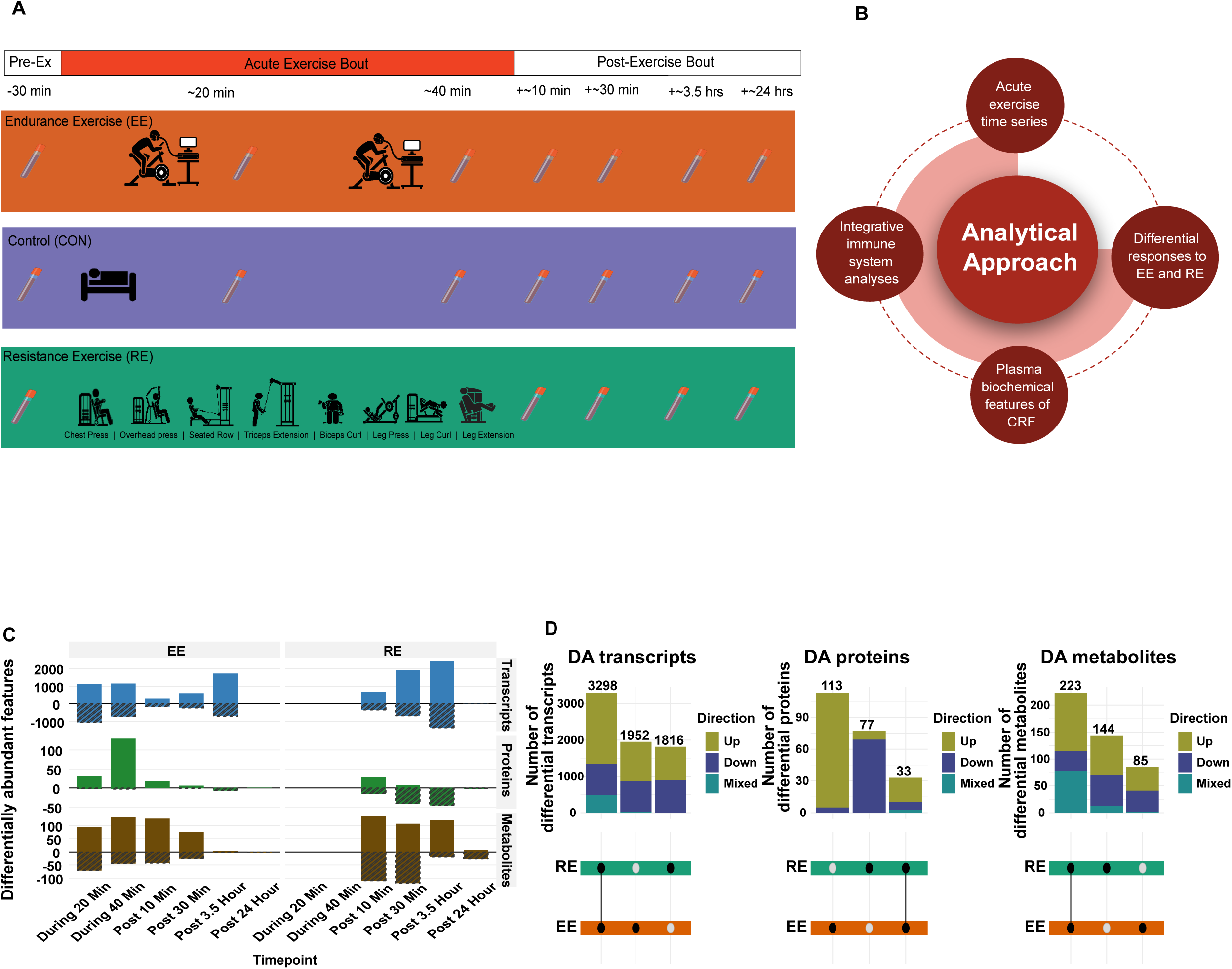
Overview of clinical study design. (A) Overview of the study design showing sedentary participants randomized to endurance exercise (EE), resistance exercise (RE), or control (CON). Blood biospecimens collections are indicated by blood tube icons. Samples are collected before, during (EE & CON only), or after the exercise bouts. (B) Overview of the analysis plan. (C) Histogram showing the number of analyte changes by exercise mode and -ome according to the main differential abundance (DA) model (p-adj <0.05). Analytes which increase and decrease are shown above and below the zero-line, respectively. (D) Upset plots showing the overlap of DA features (relative to control) between exercise modes within blood transcripts, plasma proteins and metabolites. Bars represent the number of gene features within a group. Black dots show the measured group, and a connected line between black dots represents shared gene features between EE and RE at any timepoint. The UpSet plot is ordered by the greatest number of features.

Clinical characteristics of the cohort are summarized in **Table 1**. Mean age and BMI of the cohort was 41 ± 15 years and 26.9 ± 4.0 kg/m^2^, respectively, and 72% of the participants were female. Most participants achieved a maximal effort during cardiopulmonary exercise testing (CPET) with peak respiratory exchange ratio of 1.16 ± 0.08. Mean VO_2_peak was 24 ± 7 ml*kg^-1^*min^-1^. Sex-stratified characteristics are shown in **Supplementary Table 1 (Table S1**). Sample sizes varied across the omic platforms due to technical constraints and participation burden (**Table S2**).

**Table 1.** Clinical Characteristics of the MoTrPAC Pre-COVID Blood Cohort

| Characteristic | Overall<br>N = 175<sup>1</sup> | CON<br>N = 37<sup>1</sup> | EE<br>N = 65<sup>1</sup> | RE<br>N = 73<sup>1</sup> |
| --- | --- | --- | --- | --- |
| Sex |  |  |  |  |
| Female | 126 (72%) | 31 (84%) | 46 (71%) | 49 (67%) |
| Male | 49 (28%) | 6 (16%) | 19 (29%) | 24 (33%) |
| Age (yrs) | 41 (15) | 42 (15) | 42 (14) | 40 (15) |
| Height (cm) | 168 (9) | 167 (8) | 168 (9) | 168 (10) |
| Weight (kg) | 76 (14) | 73 (13) | 76 (13) | 77 (14) |
| BMI (kg/m<sup>2</sup>) | 26.9 (4.0) | 26.2 (3.9) | 27.0 (4.0) | 27.2 (4.0) |
| Waist Circumference (cm) | 92 (12) | 89 (13) | 93 (11) | 93 (12) |
| Systolic BP (mmHg) | 116 (12) | 114 (11) | 117 (11) | 116 (12) |
| Diastolic BP (mmHg) | 72 (8) | 72 (8) | 73 (9) | 72 (8) |
| Hemoglobin A1c (%) | 5.30 (0.30) | 5.29 (0.21) | 5.29 (0.32) | 5.31 (0.32) |
| eGFR (mL/min/1.73 m<sup>2</sup>) | 101 (20) | 99 (19) | 103 (18) | 101 (22) |
| **Cardiopulmonary Exercise Test (CPET)** |  |  |  |  |
| VO2peak (L/min) | 1.86 (0.65) | 1.73 (0.56) | 1.90 (0.68) | 1.89 (0.65) |
| VO2peak (ml/min/kg) | 24 (7) | 24 (6) | 25 (7) | 25 (7) |
| Workload (Watts) | 152 (50) | 142 (40) | 155 (53) | 153 (52) |
| O2 Pulse (ml/beat) | 10.6 (3.3) | 10.0 (2.8) | 10.9 (3.5) | 10.8 (3.4) |
| RER | 1.16 (0.08) | 1.16 (0.08) | 1.17 (0.07) | 1.15 (0.08) |
| **Strength Tests** |  |  |  |  |
| Isometric Knee Extension Torque (Nm) | 148 (57) | 137 (53) | 145 (54) | 156 (60) |
| Hand Grip Strength (kg) | 30 (11) | 29 (8) | 30 (11) | 31 (12) |

<sup>1</sup> n (%); Mean (SD)

After quality control and annotation, analyses included 23,086 molecular features: 1,417 plasma proteins measured by antibody-based proteomics, 1,143 plasma metabolites measured across four targeted and six untargeted LC–MS platforms, and 20,526 PAXgene whole-blood transcripts. Differential abundance (DA) analysis identified molecular features altered by EE or RE compared to CON. Across all exercise timepoints, 7741 unique features were DA, including 7066 blood transcripts (34% of platform), 223 proteins (20%), and 452 metabolites (40% of the annotated features across the multiple platforms). RE produced more DA features than EE across the four shared, post-exercise timepoints (2640 versus 351 unique features, respectively), with 2951 shared DA features between modes (**Figures 1C-D**). Overlap between EE and RE across -omic layers and timepoints is shown in **Figures 1D** and **S1A**.

Notably, numerous plasma protein, metabolite, and blood transcripts changed over the 24 h period within the CON group (**Figure S1B**), reflecting a combination of factors, including fasting, biospecimen collection and other time-dependent effects. Examples include increased plasma IL-6 at 3.5 h^40^; decreases in triglycerides, and an increase in monoethylglycinexylidide, a lidocaine derivative associated with the biopsy procedure. The inclusion of the CON arm reclassified many apparent exercise responses. Among metabolites previously reported to be exercise-responsive^31^, 52 were no longer significant after comparing with CON changes (**Figures S1C-D**), indicating that changes are not due to exercise (**Table S3**).

### Plasma metabolomic changes in response to acute endurance and resistance exercise reflect distinct physiologic demands

RefMet enrichment analyses revealed distinct plasma metabolomic responses to EE and RE. (**Figure 2A**). RE elicited early enrichment of purine ribonucleosides, including inosine and xanthosine, as well as a marked increase in hypoxanthine (all p-adj<0.05), consistent with increased nucleotide turnover in response to RE.^34^ RE also produced a late decrease in phosphatidylinositols (PIs), significant only at post 24 h, and a sustained decrease in C24 bile acids from post 10 min through 24 h, suggesting a prolonged signaling response distinct from the acute substrate fluxes seen during exercise. Both classes are increasingly implicated in glucose and lipid homeostasis^41,42^, raising the possibility that they act as exercise-responsive signaling molecules (**Figures 2A-B**). In addition, RE also led to decreases in branched-chain amino acids and other key metabolites involved in anabolic signaling and nitrogen metabolism that persisted through 3.5 h of recovery.

**Figure 2.**
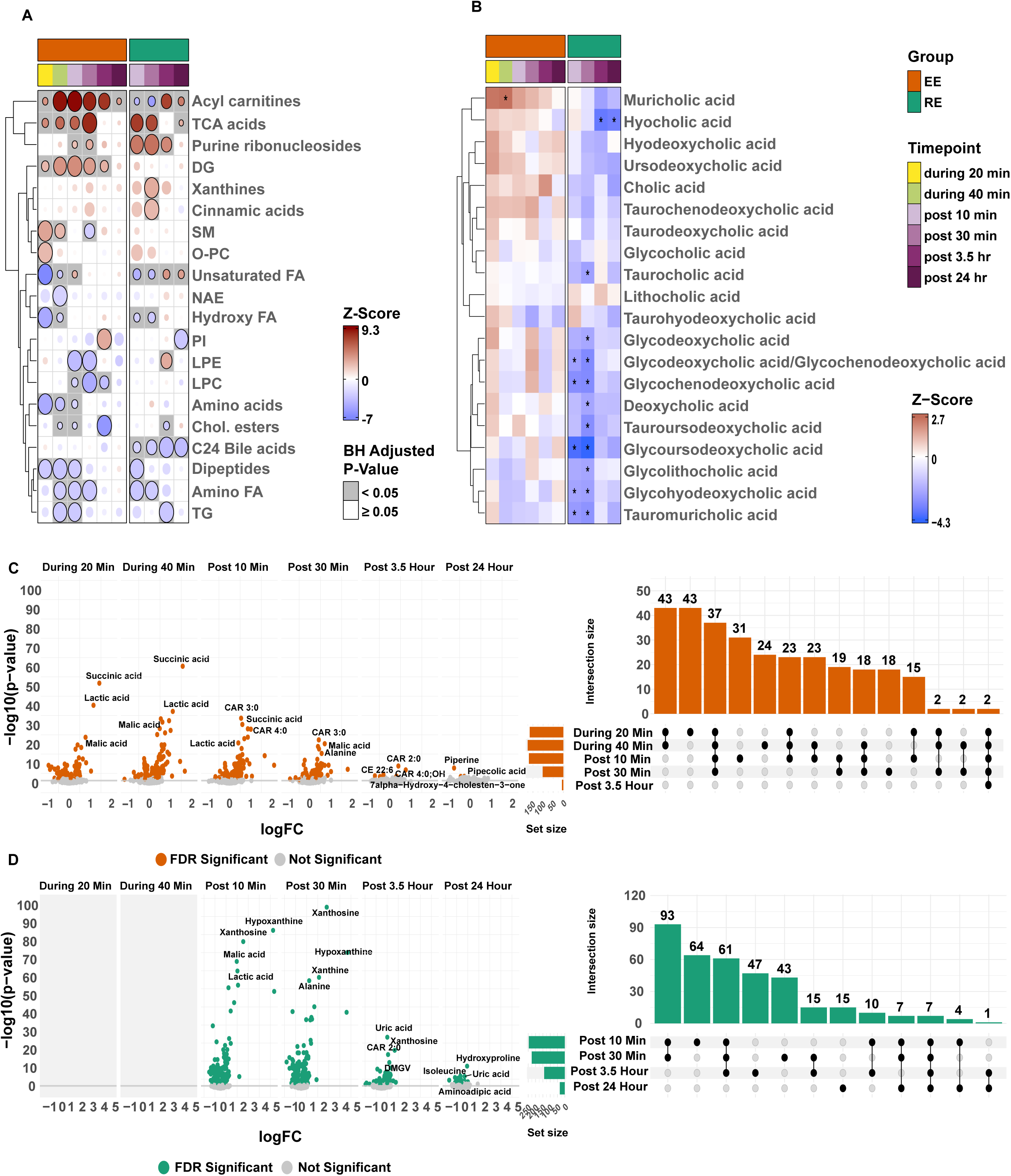
Plasma metabolomic changes in response to EE and RE. (A-B) Metabolite class enrichment analysis using CAMERA-PR applied to metabolites features at each mode-timepoint pair. Panels show enrichment results for all annotated metabolite classes that display an exercise effect (A) and C24 bile acids (B). The plotted bubbles indicate the z-score (color/intensity) and significance (grey/white background) of the enrichment term; an * denotes p-adj <0.05. The key for post-exercise timepoint sampling and modalities is presented in panel A. (C-D) Volcano plots for plasma metabolite changes (p-adj <0.05 in the main DA analysis) at each timepoint for (C) EE and (D) RE with corresponding UpSet plots depicting overlap between DA analytes at each timepoint for each exercise mode. Features in volcano plots are plotted according to their log fold-change (X-axis) and -log10(p-value) (Y-axis). Selected features are labeled. For UpSet plots, black dots indicate number of significant features per plotted group. A connected line between black dots indicates that the plotted bar represents features shared between those groups. The UpSet plot is ordered by the greatest number of features per timepoint(s).

RE also produced sustained steroid hormonal responses. At 24 h, both dehydroepiandrosterone sulfate (DHEA-S) and 5alpha-Androstane-3alpha-ol-17-one sulfate were reduced, while pregnenolone sulfate demonstrated a distinct biphasic response, with an initial increase at post 10 and 30 min, followed by a decrease below baseline levels at post 3.5 and 24 h. These findings extend prior observations that acute RE induces an immediate rise in circulating steroids followed by a progressive decline below baseline levels within 48-72 h post exercise, possibly reflecting increased tissue uptake and/or utilization.^43,44^ Most metabolite changes resolved by 24 hours but hydroxyproline, a marker of collagen turnover, remained strongly upregulated after RE, consistent with a sustained tissue-remodeling response.

In contrast to RE, EE led to transient decreases in triglycerides (TG), lysophosphatidylethanolamines (LPE) and lysophosphatidylcholines (LPC) consistent with their role as a fuel source during submaximal aerobic exercise (**Figures 2A and C-D**).^45,46^ Similarly, EE transiently increased tricarboxylic acid (TCA) intermediates including succinate, fumarate, and malate, as well as diglycerides (DG), and acylcarnitines. These changes are consistent with the increased mitochondrial fatty-acid and glucose oxidation that characterizes fuel utilization during submaximal EE.^46,47^ Ethanolamine, a product of membrane phospholipid turnover that was strongly, positively associated with cardiorespiratory fitness (CRF) in analyses described below, was among the most upregulated metabolites during EE.

### RE and EE lead to divergent lipid responses in exercise recovery

RE elicited a greater glycolytic demand than EE as reflected by a larger lactate excursion post 10 min, and enrichment analysis further supported increased glycolytic reliance by revealing divergent temporal responses in acylcarnitines and free fatty acids between exercise modalities (**Figure 2A**). Collectively, these findings likely reflect the relative hypoxia and deprivation of acylcarnitine substrates induced by RE. Several medium- and long-chain acylcarnitines increased during and immediately after EE but decreased after RE (**Figures S2A and 3A-B**). While acylcarnitine responses have been reported after acute exercise^34,48,49^, we integrated plasma and skeletal muscle metabolomics and transcriptomics data from the same participants to investigate their potential tissue of origin and regulation. The divergent plasma pattern was broadly mirrored in the skeletal muscle^38^ (**Figure 3A**), consistent with increased mitochondrial fatty acid oxidation in the muscle during EE with export of partially oxidized fatty acids as acylcarnitines.^50^ By contrast, RE, which induces restriction of blood flow and increased reliance on muscle glycogen and anaerobic glycolysis to supply ATP, results in increased lactate production.^51^ These integrated analyses support the notion that skeletal muscle is the likely source of acylcarnitines in the blood.

**Figure 3.**
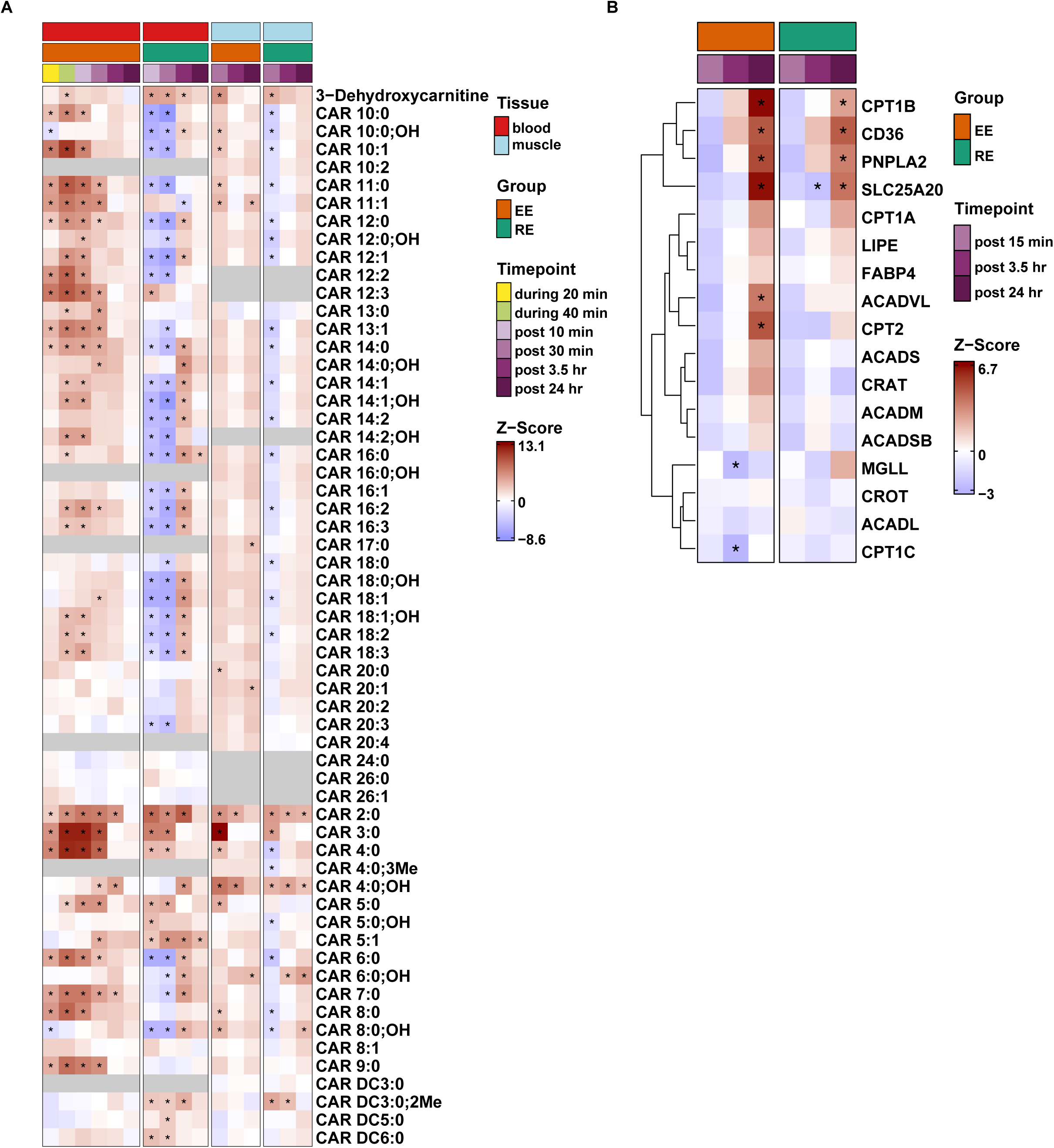
Temporal Patterns of Divergent Metabolite and Gene Expression Responses Across Plasma and Muscle. (A-B) Heatmaps of exercise mode-divergent changes in acylcarnitines (Car) and fatty acids in the (A) plasma and skeletal muscle, and (B) transcriptomic responses of key fatty acid oxidation genes in skeletal muscle following EE (orange) and RE (green) in the main DA model; an * denotes an p-adj <0.05 and a black field in (A) indicates a given Car species was not detected in a given tissue. Plasma metabolites shown correspond to those highlighted in Figure S2. The key for post-exercise timepoint sampling and modalities is presented in panel A.

We next examined mRNA changes in key regulators of lipid and acylcarnitine metabolism in skeletal muscle and found that EE induced an increase in genes involved in lipid oxidation including carnitine palmitoyltransferases (CPT1B, CPT2) and the very long chain acyl-CoA dehydrogenase (ACADVL) (post 24 h), that were either unique or of greater magnitude than RE (**Figure 3B**). Although previous studies of endurance training have shown increased expression of genes involved in fatty acid metabolism and transport in skeletal muscle, the transcriptional effects of a single acute bout have been less consistent.^52,53^ Notably, this gene expression pattern was not evident in the Meta-analysis of Skeletal Muscle Response to Exercise (MetaMEx) database of publicly available acute exercise transcriptomic responses.^54^ Collectively, our findings highlight that muscle is the likely tissue source of circulating medium- and long-chain acylcarnitines during acute exercise and provide new evidence that even a single bout of EE, and to a lesser extent RE, is sufficient to increase transcription of genes involved in lipid oxidation. In-depth analyses of mode-specific regulation of transcription factors and genes involved in lipid oxidation are presented in the MoTrPAC human skeletal muscle paper.^38^

### Plasma metabolomic responses to acute exercise differ according to baseline cardiorespiratory fitness

CRF is one of the strongest independent predictors of longevity^55–57^ and represents the integrated capacity to deliver and utilize oxygen during sustained physical activity. As such, CRF is a fundamental “readout” of metabolism. To examine relationships between circulating metabolites and rigorously ascertained physiologic traits, including CRF, we applied canonical correlation analysis (CCA), an unsupervised, multivariate method that identifies shared axes of variation between high-dimensional datasets.^58^ Using resting plasma metabolomics, the first three primary canonical variates captured distinct domains of physiologic function and metabolic domains (**Figures 4A and S3; Table S4**). The first canonical variate (CV1) was dominated by metrics of aerobic fitness, including VO_2_peak (absolute and relative), O_2_ pulse, and peak workload, and was positively weighted by the dipeptides (Pro-Phe, Leu-Pro), tryptophan, creatinine, and ethanolamine (**Figure 4B**). Because CRF and muscular strength are correlated yet distinct traits, we next examined how CV1 metabolites directly relate to measured VO_2_peak, handgrip strength, and isometric knee-extension as well as their responses to EE and RE (**Figure 4C**). A subset of metabolites demonstrated trait-specific relationships. For example, ethanolamine was more strongly associated with aerobic fitness than with strength, indicating that metabolite signatures of these traits overlap but also contain unique components.

**Figure 4.**
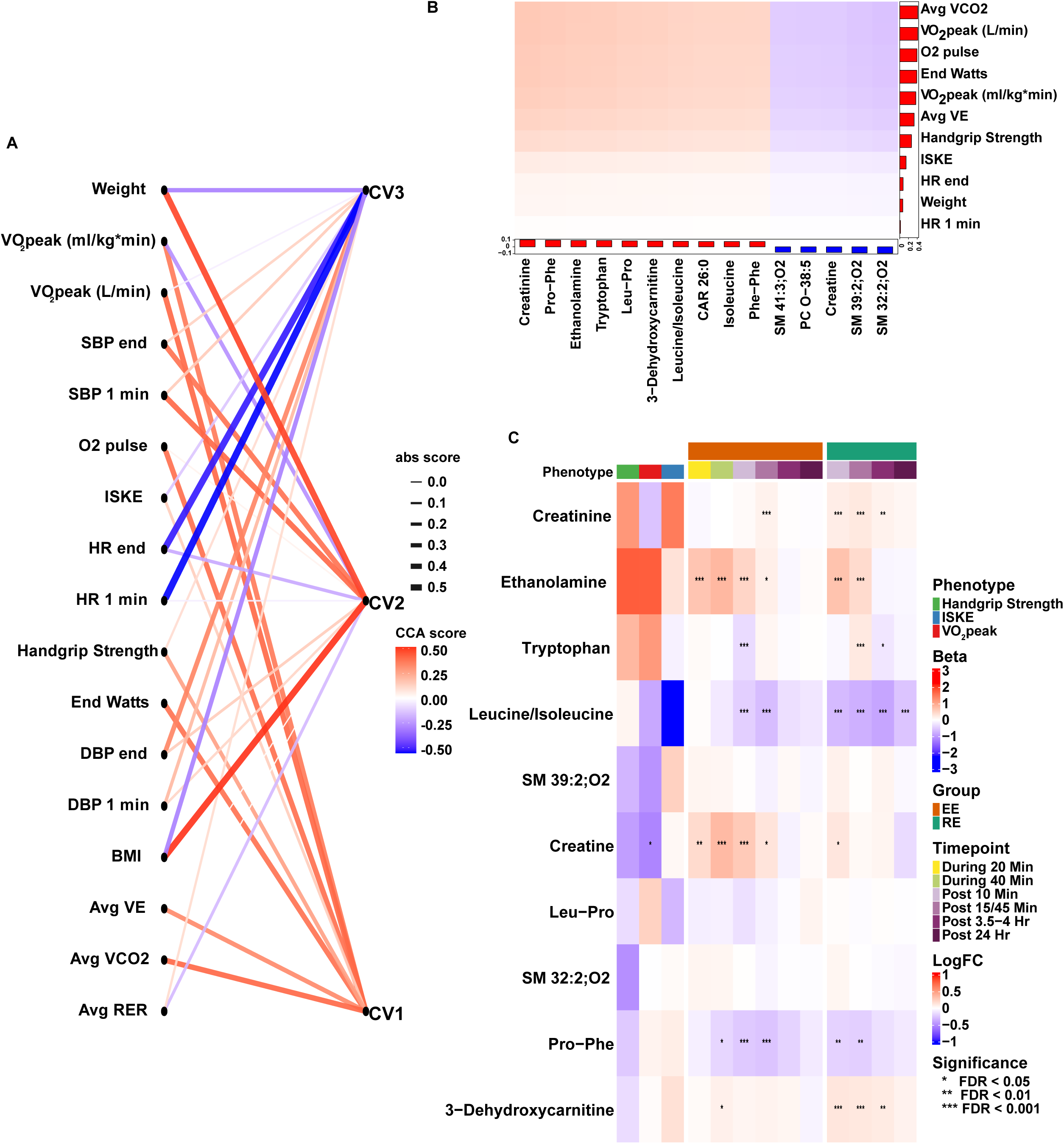
Canonical Correlation Analysis Linking Plasma Metabolites to Clinical and Exercise Traits. (A) Network plot showing associations between the top three metabolite-derived canonical variates (CV1–CV3) and clinical or exercise-related traits. Canonical variates are weighted combinations of variables (metabolites) that are maximally correlated with corresponding combinations of traits (clinical and exercise-related phenotypes). Nodes on the left represent traits and nodes on the right represent CVs; edges connect each trait-CV pair, with edge color indicating the CCA score (red = positive, blue = negative) and edge width proportional to its absolute value. CV1 is associated with key endurance and strength traits such as VO₂peak, O₂pulse and peak torque; CV2 is associated with exercise and recovery blood pressure; CV3 is associated with body composition and heart rate recovery. (B) Loadings for metabolites and phenotypes contributing to CV1. Bar height reflects the magnitude of contribution to the respective variate. Red bars denote positive loadings; blue bars denote negative loadings. (C) Heatmap summarizing associations between representative metabolites contributing to CV1 and physiological phenotypes (left), together with their acute responses to EE and RE across all sampling timepoints (right). The left panel shows standardized regression coefficients (β) relating baseline metabolite abundance to VO_₂_peak, handgrip strength, and isometric knee-extension torque. The right panel shows log₂ fold changes in metabolite abundance relative to baseline following EE and RE at each sampling timepoint. CV1 = metabolite canonical variate 1; CV2 = metabolite canonical variate 2; CV3 = metabolite canonical variate 3; end Watts = end Watts at termination; handgrip strength = average handgrip strength; peak torque = average peak torque from isokinetic test; avg VCO2 = average VCO2 during CPET; avg VE = average ventilation during CPET; DBP end = end diastolic blood pressure; SBP end = end systolic blood pressure; DBP 1 min = diastolic blood pressure at 1 minute recovery; SBP 1 min = systolic blood pressure at 1 minute recovery; HR end = end heart rate; HR 1 min = heart rate at 1 minute recovery; avg RER = average respiratory exchange ratio during CPET; O2 pulse= oxygen pulse (VO_2_/heart rate)

To test whether the CRF-associated CV1 signature was generalizable beyond MoTrPAC, we examined our findings in the HERITAGE Family Study (N=654).^59^ Of the ten metabolites contributing most strongly to the CV1 axis, six were available in HERITAGE and used to reconstruct the score using the original canonical weights. The resulting score was strongly, positively associated with VO max after adjustment for age, sex, and BMI (β=4.59, p=4.3×10 ^6^). Notably, several of these core fitness-associated metabolites were acutely modified by exercise (**Figure 4C**).

We next tested whether plasma metabolomic responses to acute EE and RE varied according to baseline VO_2_peak (ml O_2_*kg body mass^-1^min^-1^). At the first shared timepoint (post 10 min), 52 metabolites showed fitness-dependent responses after EE, with higher VO_2_peak correlated with increases in plasma levels of TCA cycle and glycolytic intermediates, metabolites involved in central carbon metabolism, as well as odd-chain saturated and polyunsaturated fatty acids (**Figures 5A and S4; Table S5**). Given that CRF influenced plasma metabolomic responses to submaximal endurance exercise in pathways known to govern endurance capacity (e.g., fatty acid oxidation, TCA cycle flux), we performed joint pathway analysis using KEGG annotation in MetaboAnalyst^60^ to test whether metabolite changes were in turn related to skeletal muscle transcriptional expression. **Figure 5B and Table S6** show the associations between skeletal muscle gene transcripts associated with VO_2_peak and plasma metabolite changes at post 10 min. Short-chain acylcarnitines changes were strongly correlated with baseline expression of canonical mitochondrial oxidative phosphorylation genes - including subunits of complexes I, II, III, IV and V - indicative of complete oxidation of acylcarnitines, while changes in amino acids, including glycine, valine, and serine were positively correlated with skeletal muscle expression of mitochondrial complex I and ATP synthase (complex V) subunits (e.g., *NDUFB9, NDUFB1, ATP5MC3*). We also found less established examples of CRF-related skeletal muscle enzymes and plasma metabolite changes. For example, we found that skeletal muscle Protein phosphatase 3 Catalytic Subunit Beta (*PPP3CB*) – and, to a lesser extent, Protein phosphatase 3 Regulatory Subunit B, alpha (*PPP3R1*), which encode critical catalytic and regulatory subunits of calcineurin, respectively, were inversely associated with both VO_2_peak and changes in fatty acids. Although calcineurin signaling has previously been described in the context of fiber-type switching^61^ and energy homeostasis^62^, these findings highlight a newly described relationship to lipid metabolism and aerobic capacity.

**Figure 5.**
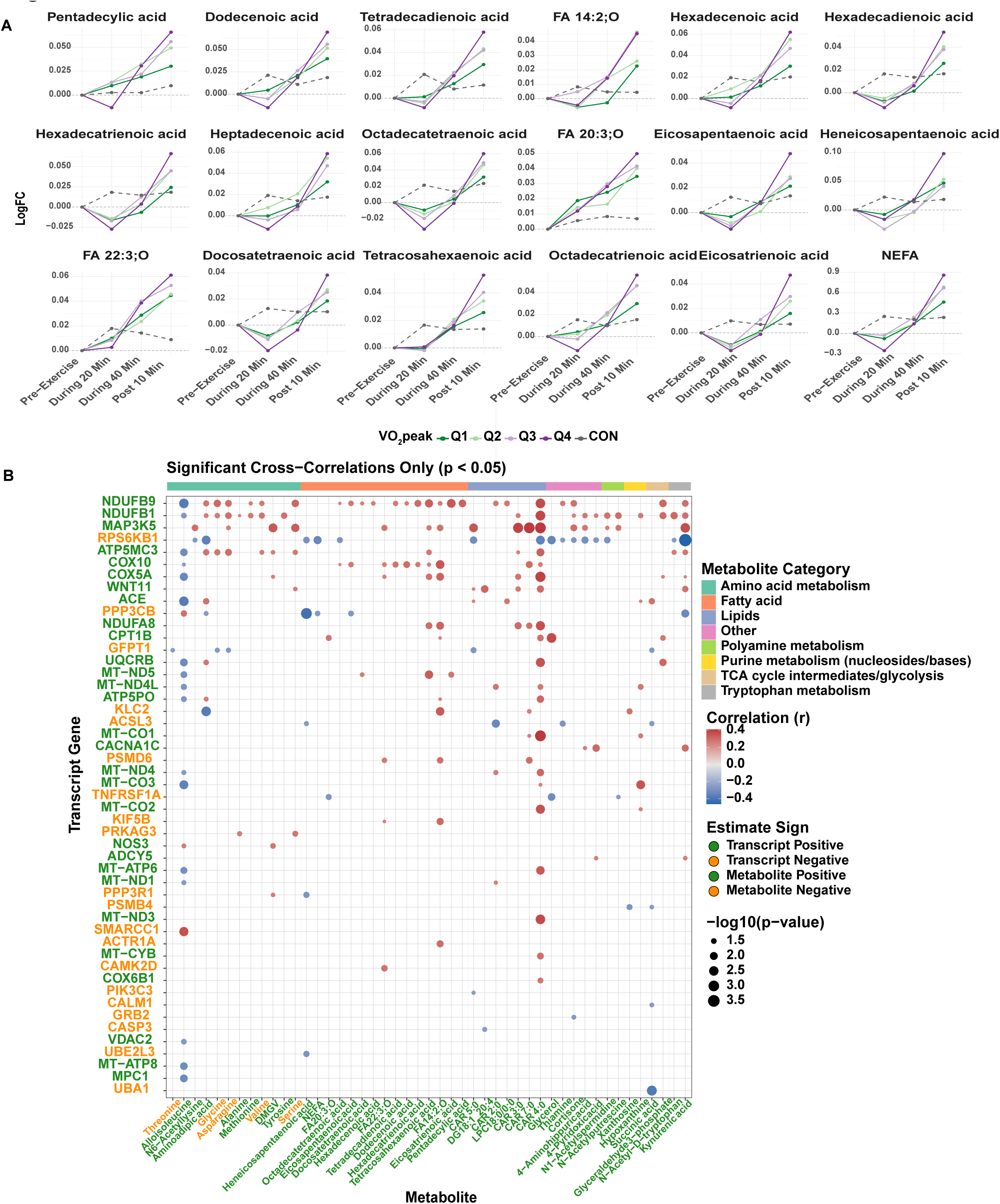
Plasma metabolite changes and skeletal muscle transcriptional expression according to VO_2_peak. (A) Plasma free fatty acids exhibiting a significant interaction between baseline VO_2_peak and EE change at the post-10 min timepoint. Representative trajectories are shown after stratifying participants into quartiles of baseline VO_2_peak (Q1–Q4). The dashed grey line represents CON. Values are presented as log_₂_ fold change relative to baseline. (B) Bubble plot showing correlations between resting skeletal muscle transcripts associated with baseline VO_2_peak and changes in plasma metabolites whose EE responses were modified by baseline VO_2_peak. Skeletal muscle transcripts associated with VO_2_peak (ml/kg/min) (p-adj <0.05) in linear regression models adjusted for age, sex, and site id. Bubble color denotes the direction and strength of the correlation, with red indicating positive correlations and blue indicating negative correlations. Bubble size represents statistical significance. Transcript labels are colored according to the direction of association with baseline VO_2_peak (green, positive; orange, negative). The colored annotation bar above the heatmap indicates metabolite functional categories.

All baseline (pre-exercise) molecular feature relationships with VO_2_peak, one-repetition maximum for leg press, leg extension, chest press, peak exercise lactate, non-esterified fatty acids, and Homeostatic Model Assessment for Insulin Resistance (HOMA-IR) are available for exploration in the MoTrPAC Multi-omic Landscape paper (in submission to *Nature Medicine*)^37^, and blood biochemical changes to acute EE and RE according to all of the clinical traits are available for interrogation in an interactive tool provided at www.motrpac-data.org.

### Resistance and endurance exercise elicit distinct plasma proteomic kinetic responses

Plasma proteomic changes during EE were highly concordant at 20 and 40 min, with 31 of 33 DA proteins changed with the same direction of effect at both timepoints (**Figure 6A**). By 40 minutes of EE, 133 plasma proteins were DA with 98% increasing. The few proteins that decreased included kallikrein-related peptidase 6 (KLK6), neuronal pentraxin receptor (NPTXR), and protein tyrosine phosphatase receptor type N2 (PTPRN2), which regulate insulin storage and secretion in response to glucose.^63^ Newly identified protein changes include increases in neprilysin (MME) - an endopeptidase that degrades vasoactive peptides and is a major pharmacologic target in the treatment of heart failure^64^, angiopoietin-related protein 7 (ANGPTL7), low-density lipoprotein receptor-related protein 1 (LRP1) – a central regulator of lipid metabolism and vascular integrity^65,66^ whose soluble form clears amyloid beta (Aβ) peptide and is a potential therapeutic target in Alzheimer’s disease^67,68^, stem cell growth factor-β (SCGF-β; or CLEC11A) - a hematopoietic^69^ and osteogenic growth factor^70^ recently shown to promote insulin secretion and beta-cell function in humans^71^, and CCN family member 1 (CCN1), highlighted as a candidate exerkine in the MoTrPAC Multi-omic Landscape paper (in submission to *Nature Medicine*).^37^ In addition, we confirm many previously identified plasma protein changes (e.g., tissue-type plasminogen activator [PLAT]; stanniocalcin-1 [STC1]; granulysin [GNLY]; fractalkine [CX3CL1]; and hepatocyte growth factor [HGF] among others^32,33,72–74^).

**Figure 6.**
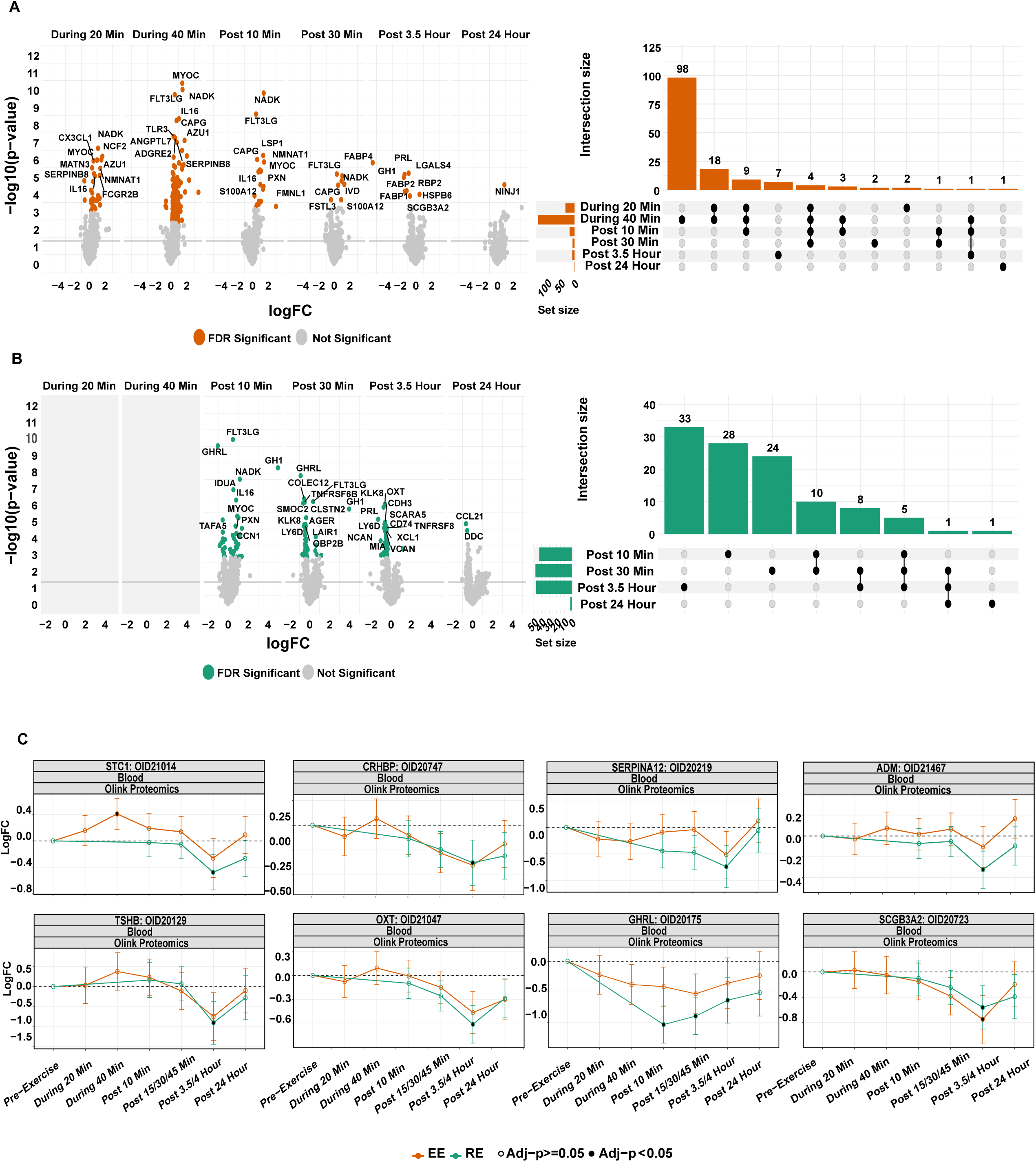
Plasma proteomic changes in response to acute endurance and resistance exercise. (A-B) Volcano plots for plasma protein changes (p-adj <0.05 in the main DA analysis) at each timepoint for (A) EE and (B) RE with corresponding UpSet plots depicting overlap between DA analytes at each timepoint for each exercise mode. Features in volcano plots are plotted according to their log fold-change (X-axis) and -log10(p-value) (Y-axis). Selected features are labeled. For UpSet plots, black dots indicate the number of significant features per the plotted group. A connected line between black dots indicates that the plotted bar represents features shared between those groups. The UpSet plot is ordered by the greatest number of features per timepoint(s). (C) Log2 fold-change in protein abundance during each acute bout, relative to control. Shown are stanniocalcin-1 (STC1), corticotropin releasing hormone binding protein (CRHBP), serpin family A member 12 (SERPINA12; Vaspin), adrenomedullin (ADM), thyroid stimulating hormone subunit beta (TSHB), oxytocin (OXT), ghrelin (GHRL), and secretoglobin family 3A member 2 (SCGB3A2). Points with black dots indicate significance (p-adj< 0.05) by linear mixed effects model. Error bars indicate 95% confidence intervals.

The early EE response also suggested immune cell mobilization and activation. Several cytokines increased, including C-X-C motif chemokines 16 and 17 (CXCL16, CXCL17), and C-C motif chemokine 16 (CCL16), as well as the monocyte chemoattractant interleukin, Interleukin- 16 (IL16). Proteins involved in leukocyte adhesion, trafficking, and activation also increased, among them integrin β7 (ITGB7), lymphocyte antigen 9 (LY9), CD300 molecule E (CD300E), and C-type lectin domain family 4 member G (CLEC4G). EE also increased two cytokine receptors, interleukin-18 receptor 1 (IL18R1) and interleukin-1 receptor type 1 (IL1R1). IL1R1 is the signaling receptor for IL-1 and the target of the anti-inflammatory exerkine, IL-1 receptor antagonist (IL1RN).^13^ Because the circulating form of IL1R1 can bind IL-1 without signaling, its increase may reflect the restraining of IL-1 activity rather than a pro-inflammatory signal. Nearly all the DA plasma proteins during EE returned to their baseline levels by 30 min following exercise.

Overall, and in a pattern that largely mirrored the plasma metabolome, RE and EE led to similar directional effects in plasma proteins, but RE led to a greater number and magnitude of effect on plasma protein changes compared to EE (**Figures 6A-B**, **S5A-C**). After RE, 85% (41/48), 98% (46/47) and 100% (2/2) of the DA proteins decreased after 30 min, 3.5 h, and 24 h, respectively. While plasma volume expansion after an acute bout of strenuous exercise may occur after initial hemoconcentration during exercise, this process has been described after 24 hours of recovery.^75,76^ Thus, the mechanisms underlying RE-induced plasma protein decreases remain unclear, however hypoxia during RE may contribute through altered protein localization and/or transcriptional repression via HIF1 and related pathways.^77–79^

In contrast to the transient protein increases seen during the EE bout, almost all (52/55) of the DA proteins in late exercise recovery (post 3.5-24 h) decreased in both modes except heat shock protein beta-6 (HSPB6 or HSP20) and nerve-injury induced protein 1 (soluble NINJ1), two proteins previously shown to have cardioprotective effects^80–82^ which increased after EE, and isovaleryl-CoA dehydrogenase (IVD) (**Figures 6A-B**). Late protein changes included growth hormone (GH1) and STC1, which demonstrated biphasic responses to EE and RE, respectively^83^, and several other hormone and hormone-like proteins including thyroid-stimulating hormone beta (TSHB), oxytocin (OXT), ghrelin (GHRL), secretoglobin family 3A member 2 (SCGB3A2), adrenomedullin (ADM), corticotropin-releasing factor-binding protein (CRHBP), and serpin A12 (SERPINA12; vaspin) (**Figure 6C**).

### Plasma protein tissue sources may differ in the exercised and rested states

The plasma secretome is comprised of proteins released through multiple processes (e.g., “classically” secreted proteins with signal peptide sequences, leakage proteins, etc.) and originating from diverse tissue sources.^84^ Thus, we sought to characterize the putative tissue sources of exercise-responsive plasma proteins using the largest human plasma secretome atlas^85^ as well as MoTrPAC human adipose and skeletal muscle data.^37–39^ We assigned all of the EE and RE responsive plasma proteins (N=223) from our dataset to their putative tissue source, as well as the certainty of a given tissue assignment (GLS score: 1-4 or common) based on data from 4 tissue- and 2 cell-type atlases across the GTEx project, EMBL-EBI Expression Atlas, and the HATLAS resource as previously described.^85–87^ We found tissue assignments for 21 tissues and all 8 cell types, as well as proteins with similar abundance in multiple tissues/cells (common), with adipose, brain, spleen, artery and liver representing the largest putative tissue sources (**Figures 7A-B**; **Table S7**). This included well-described protein-tissue source relationships (e.g., GHRL-stomach, leptin [LEP]-adipose; and neurocan [NCAN]-brain).

**Figure 7.**
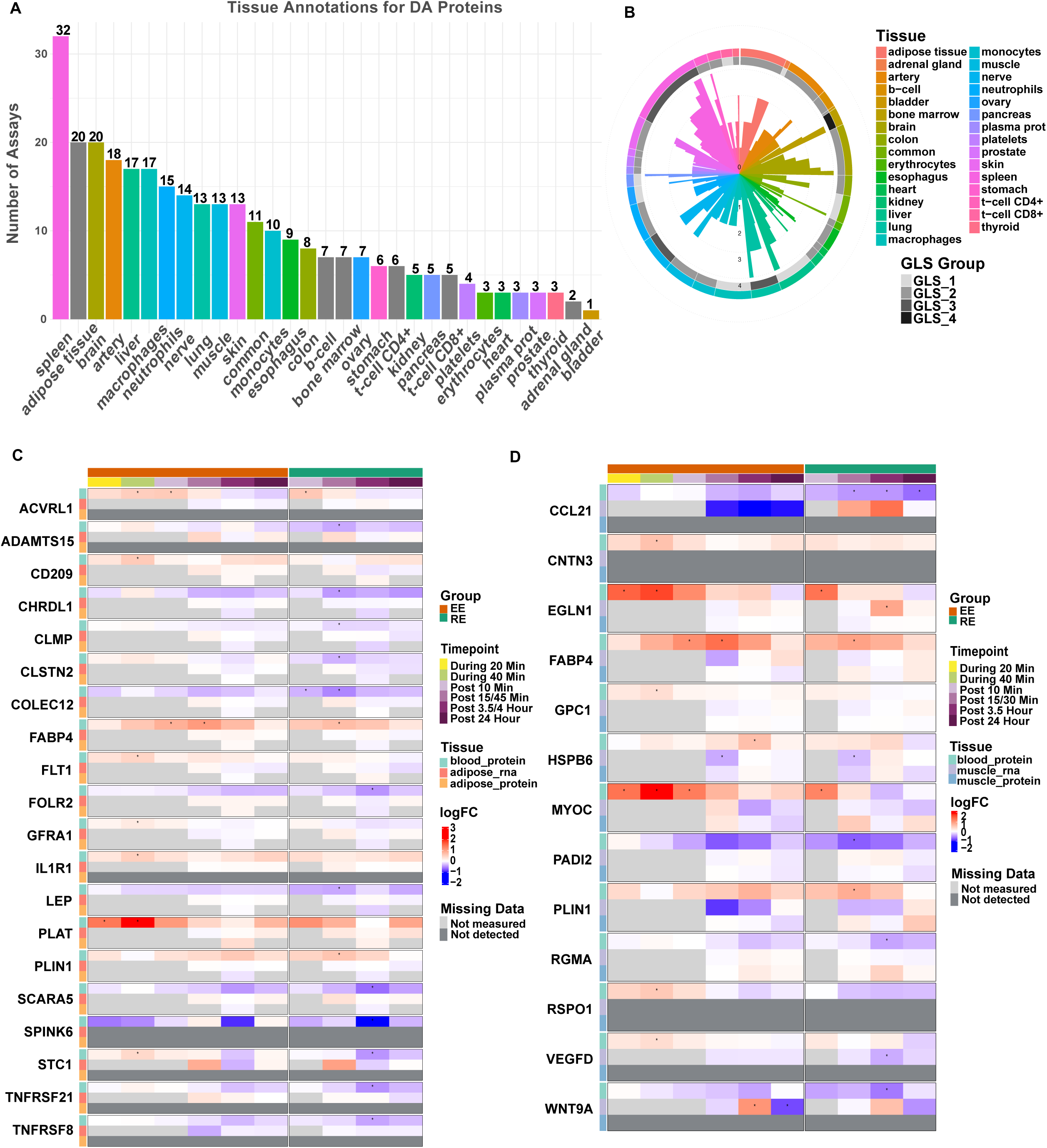
Putative tissue sources of exercise-responsive plasma proteins. (A) Tissue and cell type sources for exercise-responsive (DA) plasma proteins in MoTrPAC (N=221) according to published human plasma protein atlases.^85^ Two DA proteins did not have GLS assignments (SIGLEC5; TNFRSF6B). (B) Distribution of global longitudinal scores (GLS) for exercise-responsive protein tissue annotations. GLS is a measure of confidence for a putative protein-tissue relationship created in the HATLAS resource^85^; GLS is scored 1-4, with higher scores reflecting higher confidence for a given assignment. (C-D) Acute exercise-responsive plasma proteins annotated to (C) adipose (N=20) and/or (D) skeletal muscle (N=13) in HATLAS^85^ with corresponding heatmaps demonstrating DA changes in the plasma protein, mRNA expression, and global proteomic levels at each timepoint in MoTrPAC. *: p-adj <0.05. Light grey boxes reflect timepoints not sampled. Dark grey boxes reflect features not detected.

To investigate whether a protein’s assigned tissue source(s) at rest predicted its potential tissue source(s) during exercise, we focused on a subset of 31 proteins mapped to either adipose and/or skeletal muscle by the composite tissue label and compared their plasma responses with their transcriptional and proteomic responses to exercise in these same tissues in MoTrPAC. None of the plasma proteins demonstrated directionally concordant changes in the respective adipose or skeletal muscle transcripts and/or proteins at the same or earlier timepoint (**Figures 7C-D**). Given that exercise regulation of protein translation occurs on the order of hours,^88^ we examined all DA plasma proteins at either post 3.5 h and/or 24 h for transcriptional and global proteomic changes at or prior to that timepoint (N=55; **Figures S6A-B**). We found concordant decreases in the circulating and muscle transcriptional levels in three proteins, RAGE (AGER), BST2 (tetherin), and TNFRSF8 (CD30), a protein annotated to adipose tissue.^85^ (**Figures S6A-B)**. Collectively, these analyses suggest that a plasma protein’s tissue source in response to exercise may differ from its annotated source at rest and/or its transcriptional regulation following acute exercise.

### Resistance exercise leads to greater innate immune cellular responses

Given that whole blood is a highly dynamic tissue with rapidly changing cellular compositions - particularly among immune cell populations - we leveraged whole blood transcriptomic profiling to characterize the acute responses to EE and RE. CAMERA-PR analyses showed that the immediate response to both exercise modes was dominated by changes in immune-related pathways (**Figure 8A**). Natural killer (NK) and CD8+ T cell-related pathways increased and remained elevated post 10 and 30 min in EE and RE, respectively (**Figure 8A**). Both exercise modes also increased neutrophil-related pathways, including cytokines such as IL-4, IL-6, IL-8 and IL-17, and Toll-like receptor and CXCR4 signaling in the early post-exercise period, with more sustained changes after RE (up to 24 h). Platelet activation occurred in a similar temporal pattern, with a prolonged response following RE, as compared to EE (**Figure 8A**).

**Figure 8.**
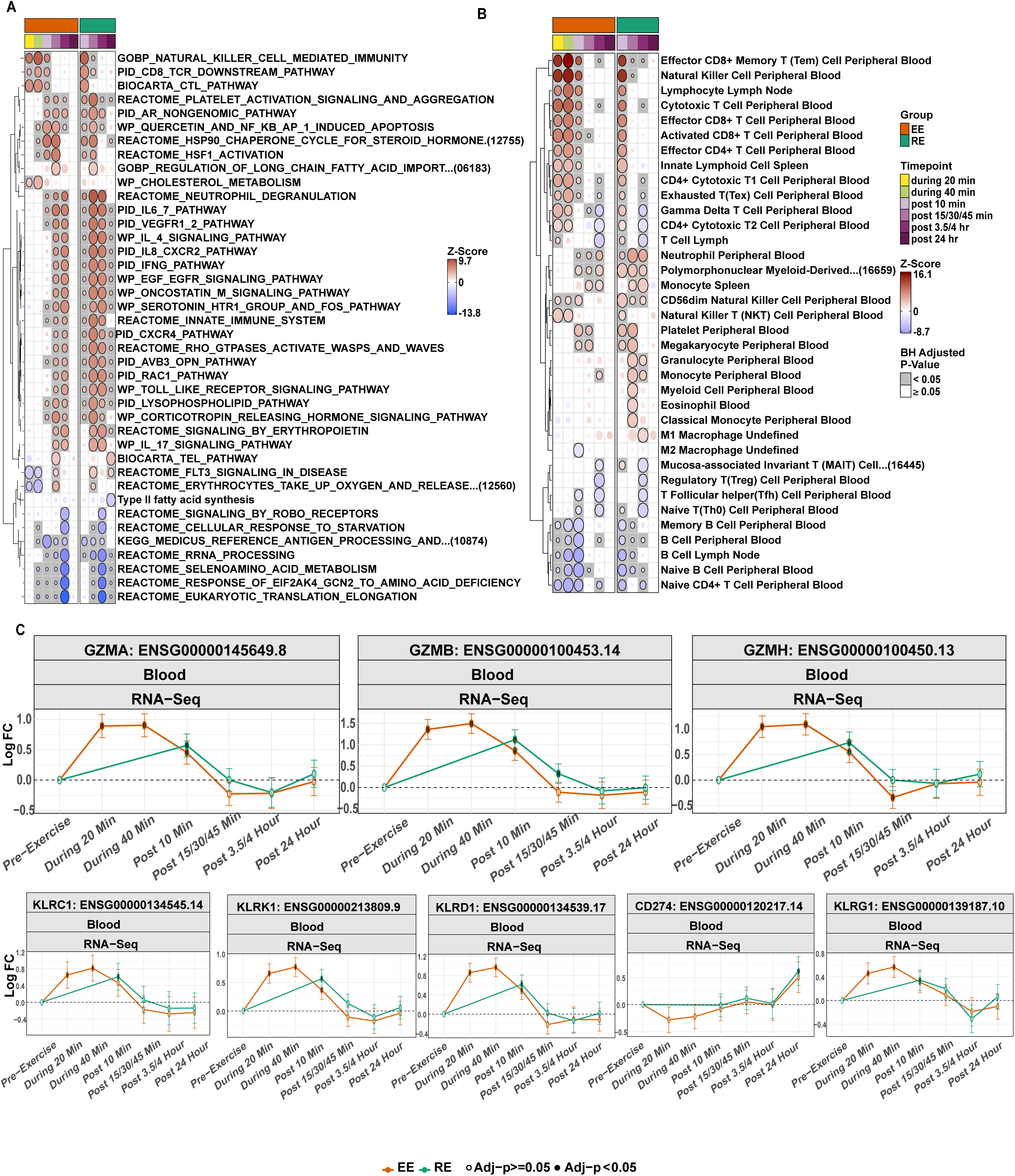
Shifts in immune cell mobilization and activation following acute EE and RE. (A) Pathway enrichment analysis using CAMERA-PR applied to transcriptomic features at each mode-timepoint pair. Features were curated to top non-redundant and biologically diverse pathway enrichments per contrast. The plotted bubbles indicate the z-score (color/intensity) and significance (grey/white background) of the enrichment term; a grey background denotes p-adj <0.05. (B) Gene set enrichment analysis of whole blood transcriptomics using CAMERA-PR referenced to the CellMarker 2.0 database for human immune cell type curations. Terms were curated for reference cell populations from the blood, peripheral blood, spleen, lymph node and undefined tissue populations. The plotted bubbles indicate the z-score (color/intensity) and significance (grey/white background) of the enrichment term. (C-D) Log2 fold-change in transcriptional abundance of cytotoxic immune cell markers from pre-exercise, relative to control are shown for EE (orange lines) and RE (green lines) over time. Points with black dots indicate significance (p-adj <0.05) by linear mixed effects model. Error bars indicate 95% confidence intervals.

To further assess shifts in either immune cellular populations and/or their level of activation, we performed GSEA using the human CellMarker 2.0 database (**Figure 8B**).^89^ Enrichment analyses demonstrated early induction and recovery of NK cells, CD8+ and CD4+ cytotoxic T cells, and (γδ) T cells, indicating a robust cytotoxic effector cellular response to both modes (**Figures 8B and S7A-C**). Plasma proteomic profiling also revealed an increase in several proteins involved in cytotoxicity, including GNLY, GZMA, and GZMB, most pronounced during EE and suggesting their early activation and cellular receptor shedding or extracellular vesicle release (**Figure S7A**).^90^ Broadly, the CD8+ T cell and NK cell signatures returned to baseline by post 3.5 h (**Figure 8B**). While samples drawn during exercise were only collected in EE, granzymes (GZMA, GZMB, GZMH) and NK cell markers (KLRC1, CD94 [KLRD1], KLRK1 [NKG2D]) revealed similar trajectories between modes at the first shared timepoint (post 10 min) (**Figure 8C**; **Figure S7B**). KLRG1, a marker of T cell differentiation, also increased in both modes while markers of other T cell subsets showed little change (**Figure S7C**).

RE produced a wider and more sustained innate immune response. CD274 (PDL1), another marker of differentiated T cells and antigen presenting cells (APCs), increased post 24 h in RE (**Figure 8C**). RE also induced unique enrichment in granulocytic populations including eosinophils and granulocytes post 30 min. Further, RE increased markers of M1 (proinflammatory) macrophages post 3.5 h, likely reflecting mobilization of non-classical monocytes given increases in CD14 and CD16 (FCGR3A and FCGR3B) without corresponding increases in CCR2 (**Figures 8B**, **S7D-E**).^91^ Exercise intensity has been positively associated with non-classical monocyte mobilization,^92^ which may facilitate tissue repair. Here, RE uniquely induced CD44 increases (**Figures S7E**), a regulator of immune cell homing, extracellular matrix repair, and myoblast differentiation.^93^ Interestingly, CD44 expression demonstrated a delayed, yet coordinated increase in the skeletal muscle in response to RE, increasing at 3.5 h, suggestive of CD44+ infiltration.^38^ Finally, RE induced greater induction of platelet activation markers, including CD63 (**Figure S7G**),^94^ consistent with the potential role for platelets in skeletal muscle regeneration.^95^

To further examine immune cell type proportions in response to EE and RE, we used CIBERSORTx and the LM22 signature matrix to deconvolute cell types from RNA-seq data (**Figure S8A-B**).^96^ Predicted monocyte abundance decreased post 3.5 h after RE with a similar, non-significant trend after EE. In parallel, gene expression associated with M0 macrophages increased post 3.5 h following both modalities (**Figure S8A-B**).

Neutrophil proportions increased up to 3.5 h following both exercise modes, consistent with pathway enrichment analyses. We further observed an acute decrease in the proportion of mast cells after both EE and RE up to 3.5 h. Resting memory CD4+ T cell populations were predicted to decrease following both EE and RE, likely reflecting proportional shifts in immune cell populations by mobilization of cytotoxic cells and monocytes. Taken together, cell deconvolution findings were broadly consistent with our enrichment analyses and highlight many shared immune responses after both EE and RE, with RE demonstrating greater innate immune cellular mobilization and/or homing.

## Discussion

Blood biochemical profiling provides an opportunity to study the systemic effects of exercise and its inter-organ communication. While several prior studies have characterized individual plasma and blood biochemical profiling technologies to acute exercise^30–32,97^, few have integrated multi-omic layers^48^, compared multiple exercise modes^34^, or included a non-exercise control group.

To our knowledge, this is the first large human study to simultaneously test the effects of acute endurance and resistance exercise using large-scale plasma proteomics, metabolomics, and whole blood transcriptomics across a 24-hour period leveraging a non-exercise control arm. We make several observations from these data. Using guideline-recommended doses of endurance and resistance exercise in untrained individuals,^98,99^ we found that RE generally induces a broader and more sustained response in plasma analytes in the post-exercise recovery period, while EE elicited more rapid and transient responses when comparing matched timepoints. We identified several mode-specific responses among specific immune cell populations, and lipid metabolism and tissue repair pathways. We observed that tissue sources of exercise-responsive plasma proteins may differ from their annotated, resting-state source(s). Further, we show how the integration of multi-tissue and multi-omic data from a single acute exercise bout may reveal new biologic relationships underlying heterogeneity in CRF. These findings and associated data are available in an easily accessible resource for the scientific community (www.motrpac-data.org).^37^

Our multi-tissue, multi-omic integrative analyses identified potentially new biology related to CRF, a robust predictor of long-term health outcomes^100^ and underscore the novelty of the MoTrPAC resource. For example, skeletal muscle expression of *PPP3CB* and *PPP3R1,* essential catalytic and regulatory subunits of calcineurin, were associated with CRF as well as CRF-dependent changes in circulating fatty acid responses to exercise. PPP3CB is itself a key regulator of mitochondrial dynamics and energy homeostasis, with both *Ppp3cb* global KO and conditional ablation of *Ppp3r1* from skeletal muscle leading to a favorable metabolic phenotype.^62^ However, while calcineurin-NFAT signaling and calcineurin’s effects on phosphodiesterases (PDE3B and PDE4D) regulate adipokine gene transcription^101^ and lipid homeostasis^102,103^, respectively, additional, mechanistic investigation is needed to test whether *PPP3CB* and *PPP3R1*, fatty acid metabolism, and CRF are directly related.

The association of short chain acylcarnitine responses to EE and VO_2_peak provides an important physiological insight. Acetylcarnitine (CAR 2:0), propionylcarnitine (CAR 3:0), and butyrylcarnitine (CAR 4:0) are produced from short-chain acyl-CoAs from both fatty acids and amino acid degradation products buffering mitochondrial acetyl-CoA/free CoA balance and supporting oxidative flux.^104^ Individuals with higher baseline CRF likely sustain a higher absolute rate of acetyl-CoA flux during a standardized bout of EE, producing more substrate for this buffering reaction resulting in increased muscle and plasma acylcarnitines. This observation can be further explored in MoTrPAC data from highly active adults undergoing the same acute exercise bout, which are currently under analysis (ref marker paper).

We note several new, acute exercise mode-specific findings among blood transcriptomics that both extend limited existing data^97^ and integrate plasma proteomic data. Both EE and RE induced the rapid expression of cytotoxic NK and CD8+ T cell markers in the early post-exercise period, and further, increased gene expression among serine proteases contained in cytotoxic granules in NK cells and CD8+ T cells, including granulysin and granzymes B, H and A during EE and post 10 min in both modes. Taken together, these data provide potential mechanistic insights behind the increased NK cell and T-cell killing of target cells in health and disease.^105–107^ In contrast, only RE strongly induced eosinophil, monocyte, and platelet activation signatures, cell types associated with skeletal muscle repair.^108–110^ Importantly, these findings were made despite MoTrPAC participants undergoing exercise familiarization sessions, a protocol performed to mitigate the undue influence of exercise, particularly RE, on a disproportionate inflammatory response through muscle damage.^36^ Cell type marker expression generally agreed with deconvolution-based estimates of cell abundance but not uniformly, likely due to exercise-induced changes in cell numbers and within cell gene expression. Thus, while deconvolution serves as a strong predictive tool for shifts in cellular proportions in various resting physiological states, the dynamic state and molecular responses to acute exercise highlight the need for future studies using flow cytometry and/or single cell profiling to further inform our understanding of the immune response to exercise.^111^

A notable feature of this study is the inclusion of a non-exercise control group which is absent from most past exercise studies.^30–32,34,48,112^ Several molecular changes we identified in the CON group mimicked the temporal pattern following EE and RE, highlighting the potential confounding effects of circadian rhythm, fasting, and tissue biopsies on the presumed dynamic changes induced by exercise.

Our in silico analyses of the exercise-responsive plasma secretome highlights the limitations of using resting tissue atlases to infer a plasma protein’s putative exercise tissue source. While there are notable examples of solid tissue transcriptional and/or proteomic changes that correlate with plasma protein levels, such as CCN1^37^, we generally did not observe this pattern. There are likely several factors for this discordance, including a given protein’s tissue specificity and that a protein’s secretory process may be distinct from its tissue transcriptional regulation (e.g., pre-synthesized protein release; likely the mechanism for CCN1^37^). Further, plasma proteins in which basal (resting) tissue gene and/or protein expression are low may be induced after an exercise stimulus. Indeed, IL-6 is a well-established example, with its dominant resting tissue sources^113^ distinct from its tissue source during exercise, skeletal muscle.^114^ Here, we find that TNFRSF8, a protein whose predominant basal expression is in adipose tissue, similarly displays a distinct tissue response after an acute bout exercise. A strength of the MoTrPAC study is the ability to relate temporal skeletal muscle, and adipose and blood tissue transcriptional and proteomics data with each other, however additional strategies, including the use of proximity-labeling technologies^115,116^ will be needed to help draw a more comprehensive assessment of the tissue-specific contributions to the exercise secretome.

We note several limitations of this work. The study was interrupted due to the SARS-CoV-2 pandemic, producing age- and sex imbalances and limiting the power for subgroup analyses, however we anticipate future studies in MoTrPAC’s post-COVID suspension cohort will be able to determine exercise response variation. In addition, few participants completed the 12-week training intervention due to the suspension, limiting our ability to relate pre-training, acute exercise findings to exercise training adaptations. As mentioned earlier, we are unable to distinguish whether whole blood transcriptional changes are driven by both shifts in cell proportions and/or changes in transcript abundance, although we employed alternative strategies to support immune analyses including our use of plasma proteomics and GSEA enriched for flow cytometry markers of immune cell phenotype.^89,117^ The Olink proteomics platform version used here has limited coverage of curated pathways, precluding meaningful enrichment.^118^ Finally, while RE appears to elicit larger magnitude effects than EE across numerous circulating analytes in this study, our findings are specific to the relative intensities of the unaccustomed resistance and aerobic bouts in MoTrPAC and may not be more generalizable to other exercise interventions.

In summary, acute EE and RE produced distinct temporal and molecular responses in blood and plasma. By integrating plasma proteomics, metabolomics, whole-blood transcriptomics, tissue data, and a non-exercise control arm, this study identifies exercise-responsive analytes, pathways, and candidate mechanisms related to exercise mode and baseline fitness. These data provide a resource for the broader scientific community to further investigate the biology of exercise.

## Methods

### Experimental model and study participant details

#### IRB

The Molecular Transducers of Physical Activity Consortium (MoTrPAC) (NCT03960827) is a multicenter study designed to isolate the effects of structured exercise training on the molecular mechanisms underlying the health benefits of exercise and physical activity. Methods are described here in sufficient detail to interpret results, with references to supplemental material and prior publications for detailed descriptions. The present work includes data from prior to the suspension of the study in March 2020 due to the COVID-19 pandemic. The study was conducted in accordance with the Declaration of Helsinki and approved by the Institutional Review Board of Johns Hopkins University School of Medicine (IRB protocol # JHMIRB5; approval date: 05/06/2019). All MoTrPAC participants provided written informed consent for the MoTrPAC study indicating whether they agreed to open data sharing and at what level of sharing they wanted. Participants could choose to openly share all de-identified data, with the knowledge that they could be reidentified, and they could also choose to openly share limited de-identified individual level data, which is lower risk of re-identification. All analyses and resulting data and results are shared in compliance with the NIH Genomic Data Sharing (GDS) policy and DSMB requirements for the randomized study.

#### Participant Characteristics

Volunteers were screened to: (a) ensure they met all eligibility criteria; (b) acquire phenotypic assessments of the study population; and (c) verify they were healthy and medically cleared to be formally randomized into the study.^35,36^ Participants were then randomized into intervention groups stratified by clinical site and completed a pre-intervention baseline acute test. A total of 176 participants completed the baseline acute test (endurance exercise [EE]=66, resistance exercise [RE]=73, and control [CON]=37), and among those, 175 (99%) had at least one biospecimen sample collected. Participants then began 12 weeks of exercise training or control conditions. Upon completion of the respective interventions, participants repeated phenotypic testing and the acute test including biospecimen sampling. Due to the COVID-19 suspension, only 45 participants (26%) completed the post-intervention follow-up acute exercise test (some with less than 12 weeks of training), with 44 completing the post-intervention biospecimen collections. See MoTrPAC Clinical Landscape (in submission to *Nature Medicine*)^35^ for further detail.

#### Participant Assessments

As described, prior to randomization, participant screening was completed via questionnaires, and measurements of anthropometrics, resting heart rate and blood pressure, fasted blood panel, cardiorespiratory fitness, muscular strength, and free-living activity and sedentary behavior.^35,36^ Cardiorespiratory fitness was assessed using a cardiopulmonary exercise test (CPET) on a cycle ergometer (Lode Excalibur Sport Lode BV, Groningen, The Netherlands) with indirect calorimetry. Quadriceps strength was determined by isometric knee extension of the right leg at a 60° knee angle using a suitable strength device.^36^ Grip strength of the dominant hand was obtained using the Jamar Handheld Hydraulic Dynamometer (JLW Instruments, Chicago, IL). See^35^ for participant assessment results.

#### Acute Exercise Intervention

The pre-intervention baseline EE acute bouts (**Figure 1**) were composed of three parts: (1) a 5 minute (min) warm up at 50% of the estimated power output to elicit 65% VO_2_peak, (2) 40 min cycling at ∼65% VO_2_peak, and (3) a 1 min recovery at ∼25 W. The pre-intervention baseline RE acute bouts were composed of a 5 min warm-up at 50-60% heart rate reserve (HRR) followed by completion of three sets each of five upper (chest press, overhead press, seated row, triceps extension, biceps curl) and three lower body (leg press, leg curl, leg extension) exercises to volitional fatigue (∼10RM) with 90 sec rest between each set. Participants randomized to CON did not complete exercise during their acute tests. Participants rested supine for 40 min to mirror the EE and RE acute test schedule. See^35^ for acute bout results.

#### Biospecimen Collection

To standardize conditions prior to the acute test, participants were instructed to comply with a variety of controls related to COX-inhibitors, biotin, caffeine, alcohol, exercise, and final nutrition consumption.^36^ Blood, muscle (SKM), and adipose (AT) samples were collected for the acute test at specific timepoints before, during, and after exercise.^37^

Participants arrived fasted in the morning, and rested supine for at least 30 min prior to pre-exercise biospecimen collections. All participants had pre-exercise sampling from all three tissues. After the pre-exercise biospecimen collections, the subsequent collection timepoints varied for each training group and their randomized temporal profile (early, middle, late, all). Temporal profiles were used to reduce the burden of repeat sampling while still maintaining biochemical coverage of the full post-exercise period.^37^ During and after exercise, biospecimen collection varied by randomized group, tissue, and temporal group. At 20 and 40 min of exercise, blood was collected from only EE and CON groups due to limitations in safely obtaining blood during RE. Immediately post exercise, at 10 minutes, a blood sample was collected in all three groups. Finally, samples from all three tissues were collected at early (SKM at 15, blood at 30, and AT at 45 min post-exercise), intermediate (blood and SKM at 3.5 h, and AT at 4 h post-exercise), and late (24 h post-exercise in all tissues) timepoints. The post-exercise timepoint began at the completion of the 40 minutes of cycling, third set of leg extension, or 40 minutes of rest, depending on the randomized group. Except for the EE blood collection timepoints during exercise, participants rested supine or seated for biospecimen collections. The collection and processing protocols for each tissue were previously described in the human clinical protocol design paper.^36^ See^35^ for biospecimen collection success results.

The number of biospecimen samples that were profiled on any given assay varied by tissue, timepoint, and randomized intervention group for feasibility reasons. Multi-omic coverage is summarized here.^37^

#### Post training intervention data

As mentioned, 44 of the 175 individuals who began the trial completed at least a portion of the training intervention period and the post-training acute bout session. These biospecimens were collected, and molecular data were generated for them. While these data are to be shared with the scientific community as detailed in the MoTrPAC Data Sharing Plan, the data are not analyzed here due to the low sample size and potential confounding between participant characteristics and groups of interest. Rather, the focus is on the baseline acute bout data. The full cohort dataset will allow exploration of multiple hypotheses related to training effects and is forthcoming.

#### HERITAGE Family Study

To externally validate the CRF metabolite signature identified in MoTrPAC, we leveraged data from the HERITAGE Family Study, a multicenter 20-week supervised endurance exercise training study of previously sedentary, healthy Black and White adults (17–65 years).^59^ Participants underwent standardized cardiopulmonary exercise testing and metabolomics profiling was performed on fasting plasma collection at rest, before the 20-week training period. Of the ten metabolites with the highest positive and negative loadings on CV1 from the MoTrPAC canonical correlation analysis, six were measured in HERITAGE (creatinine, tryptophan, 3-dehydroxycarnitine, isoleucine, creatine, and SM 32:2;O2). A CRF metabolomic score was calculated by applying the corresponding MoTrPAC CV1 canonical weights to these six metabolites, and its associations with VO max was evaluated using multivariable linear regression adjusted for age, sex, and BMI.

## Molecular Quantification Methods

### Sample identification quality control

To detect sample mislabeling and sex discordance across multi-omic datasets, we employed Somalier (v0.3.1)^119^ on BAM files from WGS, RNA-seq, and ATAC-seq samples using the RNA-seq sites VCF (sites.hg38.rna.vcf.gz). Somalier computes pairwise relatedness (ranging from 0 to 1) and identity-by-state (IBS) metrics to verify sample identity across tissues, time points, and assay types. Specifically, IBS0 represents the number of sites where one sample is homozygous-reference and another is homozygous-alternative, while IBS2 represents sites where samples share identical genotypes. Samples originating from the same individual were expected to exhibit high relatedness (>0.9), low IBS0, and high IBS2 values.

Sex concordance was verified using complementary approaches tailored to each data type. For WGS and ATAC-seq, genetic sex was inferred directly using Somalier. For RNA-seq data, sex was determined using a custom expression-based script leveraging well-established sex-linked markers (e.g., high XIST expression with low or absent DDX3Y, RPS4Y1, KDM5D, and UTY expression in females).^120,121^

We developed an integrated quality control pipeline combining Somalier metrics with sex concordance checks to systematically identify mislabeled samples. Samples were flagged as potentially mislabeled if they demonstrated very low relatedness (<0.2) and elevated IBS0 values relative to all other samples from the same participant across time points, tissues, and assays. No mislabeled samples or sex discordances were detected in the RNA-seq datasets.

### Transcriptomics

RNA Sequencing (RNA-Seq) was performed at Stanford University and the Icahn School of Medicine at Mount Sinai. Processing randomization for blood, muscle was done according to https://github.com/MoTrPAC/clinical-sample-batching. See below for adipose randomization considerations.

#### Extraction of total RNA

Tissues (∼10 mg for muscle, ∼50 mgs for adipose) were disrupted in Agencourt RNAdvance tissue lysis buffer (Beckman Coulter, Brea, CA) using a tissue ruptor (Omni International, Kennesaw, GA, #19-040E). Total RNA was extracted in a BiomekFX automation workstation according to the manufacturer’s instructions for tissue-specific extraction. Total RNA from 400 μL of blood collected in PAXgene tubes (BD Biosciences, Franklin Lakes, NJ, # 762165) was extracted using the Agencourt RNAdvance blood specific kit (Beckman Coulter). Two tissue-specific consortium reference standards were included to monitor the sample processing QC. The RNA was quantified by NanoDrop (ThermoFisher Scientific, # ND-ONE-W) and Qubit assay (ThermoFisher Scientific), and the quality was determined by either Bioanalyzer or Fragment Analyzer analysis.

#### mRNA Sequencing Library Preparation

Universal Plus mRNA-Seq kit from NuGEN/Tecan (#9133) were used for generation of RNA-Seq libraries derived from poly(A)-selected RNA according to the manufacturer’s instructions. Universal Plus mRNA-Seq libraries contain dual (i7 and i5) 8 bp barcodes and an 8 bp unique molecular identifier (UMI), which enable deep multiplexing of NGS sequencing samples and accurate quantification of PCR duplication levels. Approximately 500ng of total RNA was used to generate the libraries for muscle, 300ng for adipose, and 250ng of total RNA was used for blood. The Universal Plus mRNA-Seq workflow consists of poly(A) RNA selection, RNA fragmentation and double-stranded cDNA generation using a mixture of random and oligo(dT) priming, end repair to generate blunt ends, adaptor ligation, strand selection, AnyDeplete workflow to remove unwanted ribosomal and globin transcripts, and PCR amplification to enrich final library species. All library preparations were performed using a Biomek i7 laboratory automation system (Beckman Coulter). Tissue-specific reference standards provided by the consortium were included with all RNA isolations to QC the RNA.

#### RNA Sequencing, quantification, and normalization

RNA sequencing, quantification, and normalization Pooled libraries were sequenced on an Illumina NovaSeq 6000 platform (Illumina, San Diego, CA, USA) to a target depth of 40 million read pairs (80 million paired-end reads) per sample using a paired-end 100 base pair run configuration. In order to capture the 8-base UMIs, libraries were sequenced using 16 cycles for the i7 index read and 8 cycles for the i5 index read. Reads were demultiplexed with bcl2fastq2 (v2.20.0) using options --use-bases-mask Y*,I8Y*,I*,Y* --mask-short-adapter-reads 0 -- minimum-trimmed-read-length 0 (Illumina, San Diego, CA, USA), and UMIs in the index FASTQ files were attached to the read FASTQ files. Adapters were trimmed with cutadapt (v1.18),^122^ and trimmed reads shorter than 20 base pairs were removed. FastQC (v0.11.8) was used to generate pre-alignment QC metrics. STAR (v2.7.0d)^123^ was used to index and align reads to release 38 of the Ensembl Homo sapiens (hg38) genome and Gencode (Version 29). Default parameters were used for STAR’s genomeGenerate run mode; in STAR’s alignReads run mode, SAM attributes were specified as NH HI AS NM MD nM, and reads were removed if they did not contain high-confidence collapsed splice junctions (--outFilterType BySJout). RSEM (v1.3.1)^124^ was used to quantify transcriptome-coordinate-sorted alignments using a forward probability of 0.5 to indicate a non-strand-specific protocol. Bowtie 2 (v2.3.4.3)^125^ was used to index and align reads to globin, rRNA, and phix sequences in order to quantify the percent of reads that mapped to these contaminants and spike-ins. UCSC’s gtfToGenePred was used to convert the hg38 gene annotation (GTF) to a refFlat file in order to run Picard CollectRnaSeqMetrics (v2.18.16) with options MINIMUM_LENGTH=50 and RRNA_FRAGMENT_PERCENTAGE=0.3. UMIs were used to accurately quantify PCR duplicates with NuGEN’s “nodup.py” script (https://github.com/tecangenomics/nudup). QC metrics from every stage of the quantification pipeline were compiled, in part with multiQC (v1.6). The openWDL-based implementation of the RNA-Seq pipeline on Google Cloud Platform is available on Github (https://github.com/MoTrPAC/motrpac-rna-seq-pipeline). Filtering of lowly expressed genes and normalization were performed separately in each tissue. RSEM gene counts were used to remove lowly expressed genes, defined as having 0.5 or fewer counts per million in at least 10% of samples. These filtered raw counts were used as input for differential analysis with the variancePartition::dream,^126^ as described in the statistical analysis methods section. To generate normalized sample-level data for downstream visualization, filtered gene counts were TMM-normalized using edgeR::calcNormFactors, followed by conversion to log counts per million with edgeR::cpm.

Principal Component Analysis and calculation of the variance explained by variables of interest were used to identify and quantify potential batch effects. Based on this analysis, processing batch (muscle and blood, see below for adipose), Clinical Site (all tissues), percentage of UMI duplication, and RNA Integrity Number (RIN) technical effects were regressed out of the TMM-normalized counts via linear regression using limma::RemoveBatchEffect function in R.^127^ A design matrix including age, sex, and a combination of group and timepoint was used during batch effect removal to avoid removing variance attributable to biological effects.

#### Adipose tissue processing considerations

In the adipose tissue, RNA extraction was performed in separate batches according to exercise mode, so the extraction batch and exercise mode were perfectly collinear. This collinearity was identified at the RNA extraction step and samples were randomized prior to construction of cDNA libraries. Ultimately, under this processing implementation, differences in gene expression attributable to exercise group can be impossible to disentangle from RNA extraction batch effects, so the batch variable was not regressed out in the technical effect stage or included as a covariate in the differential analysis and left as an experimental limitation.

#### RNA Quality Inclusion Criteria

In the blood transcriptomic data, 79/1032 processed adult-sedentary samples had RIN values under 5. Based on established guidelines and internal quality control assessments, any samples with a RIN score below 5 were excluded from further analysis due to concerns about potential degradation artifacts.

### Proteomics

#### Plasma proteomics

Plasma proteomics profiling was performed using the Olink (Uppsala, Sweden) assay. The Proximity Extension Assay (PEA) technology has been described.^128^ Briefly, a ∼20μL plasma sample is first incubated with two distinct antibodies that bind proximal epitopes on the protein target. The two antibodies are conjugated with complementary DNA oligonucleotide sequences, which come in close proximity upon target binding and subsequently hybridize (immunoreaction). The sequence is then extended by DNA polymerase, creating a unique amplicon (i.e., barcode) for each individual protein, which is then amplified by polymerase chain reaction. Every sample is spiked with three internal quality controls: (1) “Immuno controls” are non-human antigens measured by Olink that assess technical variation in all three steps of the reaction; (2) “extension controls” are composed of an antibody coupled to a unique pair of DNA-tags that are always in proximity, to produce a constant signal independent of the immunoreaction. This control is used to adjust the signal from each sample with respect to extension and amplification; (3) “amplification/detection” control is a complete double-stranded DNA amplicon that does not require either proximity binding or extension to generate a signal, specifically monitoring only the amplification/detection step. Additionally, eight external controls are added to separate wells on each sample plate. Two pooled plasma or “external sample” controls, generated by pooling several μL of plasma from samples from healthy volunteers, are used to assess potential variation between runs and plates (i.e. to calculate inter-assay and intra-assay CVs). Three negative controls (buffer only) per plate are used to monitor background noise generated when DNA-tags come in close proximity without prior binding to the appropriate protein. These negative controls set the background level for each protein assay in order to calculate the limit of detection (LOD). Finally, each plate includes a Plate Control (PC) of a separate pooled EDTA plasma sample plated in triplicate; the median of the PC triplicates is used to normalize each assay and to compensate for potential variation between runs and plates. Data values for measurements below LOD are reported for all samples.

#### Adipose and skeletal muscle LC-MS proteomics

LC-MS/MS analysis of 379 muscle samples encompassing baseline and 3 post-intervention timepoints from all three groups (EE, RE, CON) was performed at the Broad Institute of MIT and Harvard (BI) and Pacific Northwest National Laboratories (PNNL). Samples were split evenly across the two sites, and a total of 14 samples were processed and analyzed at both sites to serve as cross-site replicates for evaluation of reproducibility. Additionally, 46 adipose tissue samples representing baseline and 4-hours post-intervention from all three groups were analyzed at PNNL.

#### Generation of common reference

For both tissue types, a tissue-specific common reference material was generated from bulk human samples. The common reference sample for muscle consisted of bulk tissue digest from 5 individuals at 2/3 ratio of female/male. Samples were split equally between the Broad Institute and PNNL, digested at each site following sample processing protocol described below, then mixed all digests from both sites and centrally aliquoted at PNNL. Common reference for adipose tissue was generated at PNNL using bulk tissue from 6 individuals representing a 4:2 ratio of female:male. 250 μg aliquots of both tissue specific common reference samples were made to be included in each multiplex (described below) and additional aliquots are stored for inclusion in future MoTrPAC phases to facilitate data integration.

#### Sample processing

Proteomics analyses were performed using clinical proteomics protocols described previously.^129,130^ Muscle and adipose samples were lysed in ice-cold, freshly-prepared lysis buffer (8 M urea (Sigma-Aldrich, St. Louis, Missouri), 50 mM Tris pH 8.0, 75 mM sodium chloride, 1 mM EDTA, 2 μg/ml Aprotinin (Sigma-Aldrich, St. Louis, Missouri), 10 μg/ml Leupeptin (Roche CustomBiotech, Indianapolis, Indiana), 1 mM PMSF in EtOH, 10 mM sodium fluoride, 1% phosphatase inhibitor cocktail 2 and 3 (Sigma-Aldrich, St. Louis, Missouri), 10 mM Sodium Butyrate, 2 μM SAHA, and 10 mM nicotinamide and protein concentration was determined by BCA assay. Protein lysate concentrations were normalized within samples of the same tissue type, and protein was reduced with 5 mM dithiothreitol (DTT, Sigma-Aldrich) for 1 hour at 37°C with shaking at 1000 rpm on a thermomixer, alkylated with iodoacetamide (IAA, Sigma-Aldrich) in the dark for 45 minutes at 25°C with shaking at 1000 rpm, followed by dilution of 1:4 with Tris-HCl, pH 8.0 prior to adding digestion enzymes. Proteins were first digested with LysC endopeptidase (Wako Chemicals) at a 1:50 enzyme:substrate ratio (2 hours, 25 °C, 850 rpm), followed by digestion with trypsin (Promega) at a 1:50 enzyme:substrate ratio (or 1:10 ratio for adipose tissue; 14 hours, 25 °C, 850 rpm). The next day formic acid was added to a final concentration of 1% to quench the reaction. Digested peptides were desalted using Sep-Pac C18 columns (Waters), concentrated in a vacuum centrifuge, and a BCA assay was used to determine final peptide concentrations. 250μg aliquots of each sample were prepared, dried down by vacuum centrifugation and stored at -80°C.

Tandem mass tag (TMT) 16-plex isobaric labeling reagent (ThermoFisher Scientific) was used for this study. Samples were randomized across the first 15 channels of TMT 16-plexes, and the last channel (134N) of each multiplex was used for a common reference that was prepared prior to starting the study (see above). Randomization of samples across the plexes within each site was done using https://github.com/MoTrPAC/clinical-sample-batching, with the goal to have all timepoints per participant in the same plex, and uniform distribution of groups (EE, RE, CON), sex and sample collection clinical site across the plexes.

Peptide aliquots (250 μg per sample) were resuspended to a final concentration of 5 μg/μL in 200 mM HEPES, pH 8.5 for isobaric labeling. TMT reagent was added to each sample at a 1:2 peptide: TMT ratio, and labeling proceeded for 1 hour at 25°C with shaking at 400 rpm. The labeling reaction was diluted to a peptide concentration of 2 µg/µL using 62.5 μL of 200 mM HEPES and 20% ACN. 3 μL was removed from each sample to quantify labeling efficiency and mixing ratio. After labeling QC analysis, reactions were quenched with 5% hydroxylamine and samples within each multiplex were combined and desalted with Sep-Pac C18 columns (Waters).

Combined TMT multiplexed samples were then fractionated using high pH reversed phase chromatography on a 4.6mm ID x 250mm length Zorbax 300 Extend-C18 column (Agilent) with 5% ammonium formate/2% Acetonitrile as solvent A and 5% ammonium formate/90% acetonitrile as solvent B. Samples were fractionated with 96min separation gradient at flow rate of 1mL/min and fractions were collected at each minute onto a 96-well plate. Fractions are then concatenated into 24 fractions with the following scheme: fraction 1 = A1+C1+E1+G1, fraction 2 = A2+C2+E2+G2, fraction 3 = A3+C3+E3+G3, all the way to fraction 24 = B12+D12+F12+H12 following the same scheme. 5% of each fraction was removed for global proteome analysis using immobilized metal affinity chromatography (IMAC).

#### Data acquisition

Broad Institute: Proteomic samples were analyzed on 75um ID Picofrit columns packed in-house with ReproSil-Pur 120 Å, C18-AQ, 1.9 µm beads to the length of 20-24cm. Online separation was performed on Easy nLC 1200 systems (ThermoFisher Scientific) with solvent A of 0.1% formic acid/3% acetonitrile and solvent B of 0.1% formic acid/90% acetonitrile, flow rate of 200nL/min and the following gradient: 2-6% B in 1min, 6-20% B in 52min, 20-35% B in 32min, 35-60% B in 9min, 60-90% B in 1min, followed by a 5min hold at 90%, and 9min hold at 50%.

Proteome fractions were analyzed on a Q Exactive Plus mass spectrometer (ThermoFisher Scientific) with MS1 scan across the 300-1800 m/z range at 70,000 resolution, AGC target of 3x10^6^ and maximum injection time of 5ms. MS2 scans of most abundant 12 ions were performed at 35,000 resolution with AGC target of 1x10^5^ and maximum injection time of 120ms, isolation window of 0.7m/z and normalized collision energy of 27.

PNNL: For mass spectrometry analysis of the global proteome of muscle samples, online separation was performed using a nanoAcquity M-Class UHPLC system (Waters) and a 25 cm x 75 μm i.d. picofrit column packed in-house with C18 silica (1.7 μm UPLC BEH particles, Waters Acquity) with solvent A of 0.1% formic acid/3% acetonitrile and solvent B of 0.1% formic acid/90% acetonitrile, flow rate of 200nL/min and the following gradient: 1% B for 8min, 8-20% B in 90min, 20-35% B in 13min, 35-75% B in 5min, 75-95% B in 3min, followed by a 6min hold at 90%, and 9min hold at 50%. Proteome fractions were analyzed on a Q Exactive Plus mass spectrometer (ThermoFisher Scientific) with MS1 scan across the 300-1800 m/z range at 60,000 resolution, AGC target of 3x10^6^ and maximum injection time of 20ms. MS2 scans of most abundant 12 ions were performed at 30,000 resolution with AGC target of 1x10^5^ and maximum injection time of 100ms, isolation window of 0.7m/z and normalized collision energy of 30.

For MS analysis of the global proteome of adipose samples, online separation was performed using a Dionex Ultimate 3000 UHPLC system (ThermoFisher) and a 25 cm x 75 μm i.d. picofrit column packed in-house with C18 silica (1.7 μm UPLC BEH particles, Waters Acquity) with solvent A of 0.1% formic acid/3% acetonitrile and solvent B of 0.1% formic acid/90% acetonitrile, flow rate of 200nL/min and the following gradient: 1-8% B in 10min, 8-25% B in 90min, 25-35% B in 10min, 35-75% B in 5min, and 75-5% in 3min. Proteome fractions were analyzed on a Q Exactive HF-X Plus mass spectrometer (ThermoFisher Scientific) with MS1 scan across the 300-1800 m/z range at 60,000 resolution, AGC target of 3x10^6^ and maximum injection time of 20ms. MS2 scans of most abundant 12 ions were performed at 30,000 resolution with AGC target of 1x10^5^ and maximum injection time of 100ms, isolation window of 0.7m/z and normalized collision energy of 30.

#### Data searching

Proteome data from both the Broad Institute and PNNL were searched against a composite protein database at the Bioinformatics Center (BIC) using the MSGF+ cloud-based pipeline previously described.^131^ This database comprised UniProt canonical sequences (downloaded 2022-09-13; 20383 sequences), UniProt human protein isoforms (downloaded 2022-09-13; 21982 sequences), and common contaminants (261 sequences), resulting in 42,626 sequences.

#### QC/Filtering/Normalization

*Plasma proteomics*: In total, 378 MoTrPAC sedentary adult samples were assayed across 9 plates in the same batch. No samples failed Olink standard quality control and median intra-assay and inter-assay CVs were 14.5% and 30.2%, respectively. Principal component analysis was performed and demonstrated minimal effects by plate and clinical site. Sample outliers were identified as those >3x the interquartile range for at least one of the first three principal components. Four samples (1.1%) from 3 different individuals were flagged as potential outliers and excluded from analyses. Final data are presented as NPX (normalized protein expression) values, Olink’s relative protein quantification unit on a log2 scale.

*LC-MS proteomics:* The log2 Reporter ion intensity (RII) ratios to the common reference were used as quantitative values for all proteomics features. Datasets were filtered to remove features identified from contaminant proteins and decoy sequences. Datasets were visually evaluated for sample outliers by looking at top principal components, examining median feature abundance and distributions of RII ratio values across samples, and by quantifying the number of feature identifications within each sample. No outliers were detected in the muscle tissue dataset. In the adipose datasets, four samples were flagged as outliers based on inspection of the top principal components and evaluation of the raw quantitative (median feature abundance <-1.0). These outlier samples were removed from the dataset, and the list of outlier samples can be found here.^37^ The Log_2_ RII ratio values were normalized within each sample by median centering to zero. Principal Component Analysis and calculation of the variance explained by variables of interest were used to identify and quantify potential batch effects. Based on this analysis, TMT plex (muscle and adipose), Clinical Site (muscle and adipose), and Chemical Analysis Site (muscle only, where samples were analyzed at both the Broad Institute and PNNL) batch effects were removed using Linear Models Implemented in the *limma::RemoveBatchEffect()* function in R.^127^ A design matrix including age, sex, and group_timepoint was used during batch effect removal in order to preserve the effect of all variables included in later statistical analysis. Correlations between technical replicates analyzed within and across CAS (where applicable) were calculated to evaluate intra- and inter-site reproducibility; the data from technical replicates were then averaged for downstream analysis. Finally, features with quantification in less of 30% of all samples were removed. For specific details of the process, see available code.

### Metabolomics and Lipidomics

The metabolomic analysis was performed by investigators across multiple *Chemical Analysis Sites* that employed different technical platforms for data acquisition. At the highest level, these platforms were divided into two classes: *Targeted* and *Untargeted*. The data generated by the untargeted platforms were further divided into Named (confidently identified chemical entities) and Unnamed (confidently detected, but no chemical name annotation) compounds.

Targeted metabolomics data were generated at 3 analysis Sites: Duke University, the Mayo Clinic, and Emory University. Duke quantified metabolites belonging to the metabolite classes Acetyl-CoA, Keto Acids, and Nucleic Acids ("acoa", "ka", "nuc"), Mayo quantified amines and TCA intermediates (“amines” and “tca”), while Emory quantified Oxylipins (“oxylipneg”).

The untargeted metabolomics data were generated at 3 analysis Sites: the Broad Institute, the University of Michigan, and the Georgia Institute of Technology (Georgia Tech). The Broad Institute applied Hydrophilic interaction liquid chromatography (HILIC) in the positive ion mode (hilicpos), Michigan applied reverse phase liquid chromatography in both positive and negative ion modes (“rppos” and “rpneg”) and ion-pairing chromatography in the negative mode (“ionpneg”), and Georgia Tech performed lipidomics assays using reverse phase chromatography in both positive and negative ion modes (“lrppos” and “lrpneg”).

#### LC-MS/MS analysis of branched-chain keto acids

Duke University conducted targeted profiling of branched-chain keto acid metabolites. Plasma samples containing isotopically labeled ketoleucine (KIC)-d3, ketoisovalerate (KIV)-^13^C_5_ (from Cambridge Isotope Laboratories), and 3-methyl-2-oxovalerate (KMV)-d8 (from Toronto Research Chemicals, Canada) internal standards, were subjected to deproteinization with 3M perchloric acid. Tissue homogenates were prepared at 100 mg/ml in 3M perchloric acid and 200 μL was centrifuged.

Next, 200 μL of 25 M o-phenylenediamine (OPD) in 3M HCl were added to both the plasma and tissue supernatants. The samples were incubated at 80°C for 20 minutes. Keto acids were then extracted using ethyl acetate, following a previously described protocol.^132,133^ The extracts were dried under nitrogen, reconstituted in 200 mM ammonium acetate, and subjected to analysis on a Xevo TQ-S triple quadrupole mass spectrometer coupled to an Acquity UPLC (Waters), controlled by the MassLynx 4.1 operating system.

The analytical column used was a Waters Acquity UPLC BEH C18 Column (1.7 μm, 2.1 × 50 mm), maintained at 30°C. 10 μL of the sample were injected onto the column and eluted at a flow rate of 0.4 ml/min. The gradient consisted of 45% mobile phase A (5 mM ammonium acetate in water) and 55% mobile phase B (methanol) for 2 minutes. This was succeeded by a linear gradient to 95% B from 2 to 2.5 minutes, holding at 95% B for 0.7 minutes, returning to 45% A, and finally re-equilibrating the column at initial conditions for 1 minute. The total run time was 4.7 minutes.

In positive ion mode, mass transitions of m/z 203 → 161 (KIC), 206 → 161 (KIC-d3), 189 → 174 (KIV), 194 → 178 (KIV-^5^C_13_), 203 → 174 (KMV), and 211 → 177 (KMV-d8) were monitored.

Endogenous keto acids were quantified using calibrators prepared by spiking dialyzed fetal bovine serum with authentic keto acids (Sigma-Aldrich).

#### Flow injection MS/MS analysis of acyl CoAs

Duke University conducted targeted profiling of acyl CoAs. Here, 500 μL of tissue homogenate, prepared at a concentration of 50 mg/ml in isopropanol/0.1 M KH2PO4 (1:1), underwent extraction with an equal volume of acetonitrile. The resulting mixture was then centrifuged at 14,000 x g for 10 minutes, following a previously established procedure.^134,135^

The supernatants were acidified with 0.25 ml of glacial acetic acid, and the acyl CoAs were subjected to additional purification through solid-phase extraction (SPE) using 2-(2-pyridyl) ethyl functionalized silica gel (Sigma-Aldrich), as outlined.^136^ Prior to use, the SPE columns were conditioned with 1 ml of acetonitrile/isopropanol/water/glacial acetic acid (9/3/4/4 : v/v/v/v). After application and flow-through of the supernatant, the SPE columns underwent a washing step with 2 ml of acetonitrile/isopropanol/water/glacial acetic acid (9/3/4/4 : v/v/v/v). Acyl CoAs elution was achieved with 2 ml of methanol/250 mM ammonium formate (4/1 : v/v), followed by analysis using flow injection MS/MS in positive ion mode on a Xevo TQ-S triple quadrupole mass spectrometer (Waters). The mobile phase used was methanol/water (80/20, v/v) with 30 mM ammonium hydroxide. Spectra were acquired in the multichannel acquisition mode, monitoring the neutral loss of 507 amu (phosphoadenosine diphosphate) and scanning from m/z 750 to 1100. As an internal standard, heptadecanoyl CoA was employed.

Quantification of endogenous acyl CoAs was carried out using calibrators created by spiking tissue homogenates with authentic Acyl CoAs (Sigma-Aldrich) that covered saturated acyl chain lengths from C0 to C18. Empirical corrections for heavy isotope effects, particularly ^13^C, to the adjacent m+2 spectral peaks in a specific chain length cluster were made by referring to the observed spectra for the analytical standards.

#### LC-MS/MS analysis of nucleotides

Duke University conducted targeted profiling of nucleotide metabolites. 300 μL of tissue homogenates, prepared at a concentration of 50 mg/ml in 70% methanol, underwent spiking with nine internal standards: ^13^C^10^,^15^N^5^-adenosine monophosphate, ^13^C^10^,^15^N^5^-guanosine monophosphate, ^13^C^10^,^15^N^2^-uridine monophosphate, ^13^C^9^,^15^N^3^-cytidine monophosphate, ^13^C^10^-guanosine triphosphate, ^13^C^10^-uridine triphosphate, ^13^C^9^-cytidine triphosphate, ^13^C^10^-adenosine triphosphate, and nicotinamide-1, N^6^-ethenoadenine dinucleotide (eNAD) (all from Sigma-Aldrich).

Nucleotides were extracted using an equal volume of hexane, following the procedures previously outlined.^136,137^ After vortexing and centrifugation at 14,000 x g for 5 minutes, the bottom layer was subjected to another centrifugation. Chromatographic separation and MS analysis of the supernatants were performed using an Acquity UPLC system (Waters) coupled to a Xevo TQ-XS quadrupole mass spectrometer (Waters).

The analytical column employed was a Chromolith FastGradient RP-18e 50-2mm column (EMD Millipore, Billerica, MA, USA), that was maintained at 40°C. The injection volume was 2 μL. Nucleotides were separated using a mobile phase A consisting of 95% water, 5% methanol, and 5 mM dimethylhexylamine adjusted to pH 7.5 with acetic acid. Mobile phase B comprised 20% water, 80% methanol, and 10 mM dimethylhexylamine. The flow rate was set to 0.3 ml/min. The 22-minute gradient (t=0, %B=0; t=1.2, %B=0; t=22, %B=40) was followed by a 3-minute wash and 7-minute equilibration. Nucleotides were detected in the negative ion multiple reaction monitoring (MRM) mode based on characteristic fragmentation reactions. Endogenous nucleotides were quantified using calibrators prepared by spiked-in authentic nucleotides obtained from Sigma-Aldrich.

#### LC-MS/MS analysis of amino metabolites

The Mayo Clinic conducted LC-MS-based targeted profiling of amino acids and amino metabolites, as previously outlined.^138,139^ In summary, either 20 ml of plasma samples or 5 mg of tissue homogenates were supplemented with an internal standard solution comprising isotopically labeled amino acids (U-^13^C^4^ L-aspartic acid, U-^13^C_3_ alanine, U-^13^C_4_ L-threonine, U-^13^C L-proline, U-^13^C_6_ tyrosine, U-^13^C_5_ valine, U-^13^C_6_ leucine, U-^13^C_6_ phenylalanine, U-^13^C_3_ serine, U-^13^C_5_ glutamine, U-^13^C_2_ glycine, U-^13^C_5_ glutamate, U-^13^C_6_,^15^N_2_ lysine, U-^13^C_5_,^15^N methionine, 1,1U-^13^C_2_ homocysteine, U-^13^C_6_ arginine, U-^13^C_5_ ornithine, ^13^C_4_ asparagine, ^13^C_2_ ethanolamine, D_3_ sarcosine, D_6_ 4-aminobutyric acid).

The supernatant was promptly derivatized using 6-aminoquinolyl-N-hydroxysuccinimidyl carbamate with a MassTrak kit (Waters). A 10-point calibration standard curve underwent a similar derivatization procedure after the addition of internal standards. Both the derivatized standards and samples were subjected to analysis using a Quantum Ultra triple quadrupole mass spectrometer (ThermoFischer) coupled with an Acquity liquid chromatography system (Waters). Data acquisition utilized selected ion monitoring (SRM) in positive ion mode. The concentrations of 42 analytes in each unknown sample were calculated against their respective calibration curves.

#### GC-MS analysis of TCA metabolites

The Mayo Clinic conducted GC-MS-targeted profiling of tricarboxylic acid (TCA) metabolites, as previously detailed^140,141^, with some modifications. In summary, 5 mg of tissue were homogenized in 1X PBS using an Omni bead ruptor (Omni International, Kennesaw, GA), followed by the addition of 20 μL of an internal solution containing U-^13^C labeled analytes (^13^C_3_ sodium lactate, ^13^C_4_ succinic acid, ^13^C_4_ fumaric acid, ^13^C_4_ alpha-ketoglutaric acid, ^13^C_4_ malic acid, ^13^C_4_ aspartic acid, ^13^C_5_ 2-hydroxyglutaric acid, ^13^C_5_ glutamic acid, ^13^C_6_ citric acid, ^13^C_2_,^15^N glycine, ^13^C_2_ sodium pyruvate). For plasma, 50 μL were used.

The proteins were precipitated out using 300 μL of a mixture of chilled methanol and acetonitrile solution. After drying the supernatant in the speedvac, the sample was derivatized first with ethoxime and then with MtBSTFA + 1% tBDMCS (N-Methyl-N-(t-Butyldimethylsilyl)-Trifluoroacetamide + 1% t-Butyldimethylchlorosilane) and then analyzed on an Agilent 5977B GC/MS (Santa Clara, California) under single ion monitoring conditions using electron ionization. Concentrations of lactic acid (m/z 261.2), fumaric acid (m/z 287.1), succinic acid (m/z 289.1), ketoglutaric acid (m/z 360.2), malic acid (m/z 419.3), aspartic acid (m/z 418.2), 2-hydroxyglutaratic acid (m/z 433.2), cis-aconitic acid (m/z 459.3), citric acid (m/z 591.4), isocitric acid (m/z 591.4), and glutamic acid (m/z 432.4) were measured against 7-point calibration curves that underwent the same derivatization procedure.

#### Targeted lipidomics of low-level lipids

Lipid targeted profiling was conducted at Emory University following established methodologies as previously described.^142,143^ In summary, 20 mg of powdered tissue samples were homogenized in 100 μL PBS using Bead Ruptor (Omni International, Kennesaw, GA).

Homogenized tissue samples (or plasma samples) were diluted with 300 μL 20% methanol and spiked with a 1% BHT solution to a final BHT concentration of 0.1% and pH of 3.0 by acetic acid addition. After centrifugation (10 minutes, 14000 rpm), the supernatants were transferred to 96-well plates for further extraction.

The supernatants were loaded onto Isolute C18 solid-phase extraction (SPE) columns that had been conditioned with 1000 μL ethyl acetate and 1000 μL 5% methanol. The SPE columns were washed with 800 μL water and 800 μL hexane, and the oxylipins were then eluted with 400 μL methyl formate. The SPE process was automated using a Biotage Extrahera (Uppsala, Sweden). The eluate was dried with nitrogen and reconstituted with 200 μL acetonitrile before LC-MS analysis. Sample blanks, pooled extract samples used as quality controls (QC), and consortium reference samples were prepared for analysis using the same methods. All external standards were purchased from Cayman Chemical (Ann Arbor, Michigan) at a final concentration in the range 0.01-20 μg/ml and consisted of: prostaglandin E2 ethanolamide (Catalog No. 100007212), oleoyl ethanolamide (Catalog No. 90265), palmitoyl ethanolamide (Catalog No. 10965), arachidonoyl ethanolamide (Catalog No. 1007270), docosahexaenoyl ethanolamide (Catalog No. 10007534), linoleoyl ethanolamide (Catalog No. 90155), stearoyl ethanolamide (Catalog No. 90245), oxy-arachidonoyl ethanolamide (Catalog No. 10008642), 2-arachidonyl glycerol (Catalog No. 62160), docosatetraenoyl ethanolamide (Catalog No. 90215), α-linolenoyl ethanolamide (Catalog No. 902150), oleamide (Catalog No. 90375), dihomo-γ-linolenoyl ethanolamide (Catalog No. 09235), decosanoyl ethanolamide (Catalog No. 10005823), 9,10 DiHOME (Catalog No. 53400), prostaglandin E2-1-glyceryl ester (Catalog No. 14010), 20-HETE (Catalog No. 10007269), 9-HETE (Catalog No. 34400), 14,15 DiHET (Catalog No. 10007267), 5(S)-HETE (Catalog No. 34210), 12(R)-HETE (Catalog No. 10007247), 11(12)-DiHET (Catalog No. 10007266), 5,6-DiHET (Catalog No. 10007264), thromboxane B2 (Catalog No. 10007237), 12(13)-EpOME (Catalog No. 52450), 13 HODE (Catalog No. 38600), prostaglandin F2α (Catalog No. 10007221), 14(15)-EET (Catalog No. 10007263), 8(9)-EET (Catalog No. 10007261), 11(12)-EET (Catalog No. 10007262), leukotriene B4 (Catalog No. 20110), 8(9)-DiHET (Catalog No. 10007265), 13-OxoODE (Catalog No. 38620), 13(S)-HpODE (Catalog No. 48610), 9(S)-HpODE (Catalog No. 48410), 9(S)-HODE (Catalog No. 38410), resolvin D3 (Catalog No. 13834), resolvin E1 (Catalog No. 10007848), resolvin D1 (Catalog No. 10012554), resolvin D2 (Catalog No. 10007279), 9(S)HOTrE (Catalog No. 39420), 13(S)HOTrE (Catalog No. 39620), 8-iso Progstaglandin F2α (Catalog No. 25903), maresin 1 (Catalog No. 10878), maresin 2 (Catalog No. 16369).

LC-MS data were acquired using an Agilent 1290 Infinity II chromatograph from Agilent (Santa Clara, CA), equipped with a ThermoFisher Scientific AccucoreTM C18 column (100 mm × 4.6, 2.6 µm particle size), and coupled to Agilent 6495 mass spectrometer for polarity switch multiple reaction monitoring (MRM) scan. The mobile phases consisted of water with 10 mM ammonium acetate (mobile phase A) and acetonitrile with 10 mM ammonium acetate (mobile phase B). The chromatographic gradient program was: 0.5 minutes with 95% A; 1 minute to 2 minutes with 65% A; 2.1 minutes to 5.0 minutes with 45% A; 7 minutes to 20 minutes with 25% A; and 21.1 minutes until 25 minutes with 95% A. The flow rate was set at 0.40 ml/min. The column temperature was maintained at 35°C, and the injection volume was 6 μL. For MS analysis, the MRM analysis is operated at gas temperature of 290 °C, gas flow of 14 L/min, Nebulizer of 20 psi, sheath gas temperature of 300 °C, sheath gas flow of 11 L /min, capillary of 3000 V for both positive and negative ion mode, nozzle voltage of 1500 V for both positive and negative ion mode, high pressure RF of iFunnel parameters of 150V for both positive and negative ion mode, and low pressure of RF of iFunnel parameters of 60 V for both positive and negative ion mode. Skyline (version 25.1.0.142)^144^ was utilized to process raw LC-MS data. Standard curves were constructed for each oxylipin/ethanolamide and scrutinized to ensure that all concentration points fell within the linear portion of the curve with an R-squared value not less than 0.9.

Additionally, features exhibiting a high coefficient of variation (CV) among the quality control (QC) samples were eliminated from the dataset. Pearson correlation among the QCs for each tissue type was computed using the Hmisc R library, and the figures documented in the QC report were generated and visualized with the corrplot R library.^145,146^

#### Hydrophilic interaction LC-MS metabolomics

The untargeted analysis of polar metabolites in the positive ionization mode was conducted at the Broad Institute of MIT and Harvard. The LC-MS system consisted of a Shimadzu Nexera X2 UHPLC (Shimadzu Corp., Kyoto, Japan) coupled to a Q-Exactive hybrid quadrupole Orbitrap mass spectrometer (Thermo Fisher Scientific). Plasma samples (10 μL) were extracted using 90 μL of 74.9/24.9/0.2 v/v/v acetonitrile/methanol/formic acid containing valine-d8 and phenylalanine-d8 internal standards. Following centrifugation, the supernatants were directly injected onto a 150 x 2 mm, 3 µm Atlantis HILIC column (Waters). Tissue (10 mg) homogenization was performed at 4°C using a TissueLyser II (QAIGEN) bead mill set to two 2 min intervals at 20 Hz in 300 μL of 10/67.4/22.4/0.018 v/v/v/v water/acetonitrile/methanol/formic acid valine-d8 and phenylalanine-d8 internal standards. The column was eluted isocratically at a flow rate of 250 μL/min with 5% mobile phase A (10 mM ammonium formate and 0.1% formic acid in water) for 0.5 minute, followed by a linear gradient to 40% mobile phase B (acetonitrile with 0.1% formic acid) over 10 minutes, then held at 40% B for 4.5 minutes. MS analyses utilized electrospray ionization in the positive ion mode, employing full scan analysis over 70-800 m/z at 70,000 resolution and 3 Hz data acquisition rate. Various MS settings, including sheath gas, auxiliary gas, spray voltage, and others, were specified for optimal performance.

Data quality assurance was performed by confirming LC-MS system performance with a mixture of >140 well-characterized synthetic reference compounds, daily evaluation of internal standard signals, and the analysis of four pairs of pooled extract samples per sample type inserted in the analysis queue at regular intervals. One sample from each pair was used to correct for instrument drift using “nearest neighbor” scaling while the second reference sample served as a passive QC for determination of the analytical coefficient of variation of every identified metabolite and unknown. Raw data processing involved the use of TraceFinder software (v3.3, Thermo Fisher Scientific) for targeted peak integration and manual review, as well as Progenesis QI (v3.0, Nonlinear Dynamics, Waters) for peak detection and integration of both identified and unknown metabolites. Metabolite identities were confirmed using authentic reference standards.

#### Reversed Phase-High Performance & Ion-pairing LC metabolomics

Reversed-phase and ion pairing LC-MS profiling of polar metabolites was conducted at the University of Michigan. LC-MS grade solvents and mobile phase additives were procured from Sigma-Aldrich, while chemical standards were obtained from either Sigma-Aldrich or Cambridge Isotope Labs. Plasma (50 μL aliquot) was extracted by addition of 200 μL of 1:1:1 v:v methanol:acetonitrile:acetone containing the following internal standards diluted from stock solutions to yield the specified concentrations: D_4_-succinic acid, 12.5 µM; :D_3_-malic acid,12.5 µM; D_5_ -glutamic acid,12.5 µM; D_10_ -leucine,12.5 µM; D_5_-tryptophan,12.5 µM;D_5_- phenylalanine,12.5 µM; D_3_-caffeine,12.5 µM; D_8_-lysine,12.5 µM; D_4_-chenodeoxycholic acid,12.5 µM; phosphatidylcholine(17:0/17:0),2.5 µM; phosphatidylcholine(19:0/19:0),2.5 µM; D_31_-palmitic acid,12.5 µM; D_35_-stearic acid,12.5 µM; gibberellic acid, 2.5 µM; epibrassinolide, 2.5 µM. The extraction solvent also contained a 1:400 dilution of Cambridge Isotope carnitine/acylcarnitine mix NSK-B, resulting in the following concentrations of carnitine/acylcarnitine internal standards: D_9_-L-carnitine, 380 nM; D_3_-L-acetylcarnitine, 95 nM; D_3_-L-propionylcarnitine, 19 nM; D_3_-L- butyrylcarnitine, 19 nM; D_9_-L-isovalerylcarnitine, 19 nM; D_3_-L-octanoylcarnitine, 19 nM; D_9_-L- myristoylcarnitine, 19 nM; D_3_-L-palmitoylcarnitine, 38 nM. After addition of solvent, samples were vortexed, incubated on ice for 10 minutes, and then centrifuged at 15,000 x g for 5 minutes. 150 μL supernatant were retrieved from the pellet, transferred to a glass autosampler vial with a 250 uL footed insert, and dried under a constant stream of nitrogen gas at ambient temperature. A QC sample was created by pooling residual supernatant from multiple samples. This QC sample underwent the same drying and reconstitution process as described for individual samples. Dried supernatants were stored at -80 C until ready for instrumental analysis. On the day of analysis, dried samples were reconstituted in 37.5 μL of 8:2 v:v water:methanol, vortexed thoroughly, and submitted for LC-MS.

Frozen tissue samples (skeletal muscle and adipose) were rapidly weighed into pre-tared, pre- chilled Eppendorf tubes and extracted in 1:1:1:1 v:v methanol:acetonitrile:acetone:water containing the following internal standards diluted from stock solutions to yield the specified concentrations: ^13^C_3_-lactic acid, 12.5 µM; ^13^C_5_-oxoglutaric acid, 125 nM;^13^C_5_-citric acid,1.25 µM; ^13^C_4_-succinic acid,125 nM; ^13^C_4_-malic acid, 125 nM; U-^13^C amino acid mix (Cambridge Isotope CLM-1548-1), 2.5 µg/mL; ^13^C_5_-glutamine, 6.25 µM;^15^N_2_-asparagine, 1.25 µM;^15^N_2_-tryptophan, 1.25 µM; ^13^C_6_-glucose, 62.5 µM; D_4_-thymine, 1 µM; ^15^N-anthranillic acid, 1 µM; gibberellic acid,1 µM; epibrassinolide, 1 µM. The extraction solvent also contained a 1:400 dilution of Cambridge Isotope carnitine/acylcarnitine mix NSK-B, resulting in the following concentrations of carnitine/acylcarnitine internal standards: D_9_-L-carnitine, 380 nM; D_3_-L-acetylcarnitine, 95 nM; D_3_-L-propionylcarnitine, 19 nM; D_3_-L-butyrylcarnitine, 19 nM; D_9_-L-isovalerylcarnitine, 19 nM; D_3_- L-octanoylcarnitine, 19 nM; D_9_-L-myristoylcarnitine, 19 nM; D_3_-L-palmitoylcarnitine, 38 nM. Sample extraction was performed by adding chilled extraction solvent to tissue sample at a ratio of 1 ml solvent to 50 mg wet tissue mass. Immediately after solvent addition, the sample was homogenized using a Branson 450 probe sonicator. Subsequently, the tubes were mixed several times by inversion and then incubated on ice for 10 minutes. Following incubation, the samples were centrifuged and 300 μL of the supernatant was carefully transferred to two autosampler vials with flat-bottom inserts and dried under a constant stream of nitrogen gas. A QC sample was created by pooling residual supernatants from multiple samples. This QC sample underwent the same drying and reconstitution process as described for individual samples. Dried supernatants were stored at -80 C until ready for instrumental analysis. On the day of analysis, samples were reconstituted in 60 μL of 8:2 v:v water:methanol and then submitted to LC-MS.

Reversed phase LC-MS samples were analyzed on an Agilent 1290 Infinity II / 6545 qTOF MS system with a JetStream electrospray ionization (ESI) source (Agilent Technologies, Santa Clara, California) using a Waters Acquity HSS T3 column, 1.8 µm 2.1 x 100 mm equipped with a matched Vanguard precolumn (Waters Corporation). Mobile phase A was 100% water with 0.1% formic acid and mobile phase B was 100% methanol with 0.025% formic acid. The gradient was as follows: Linear ramp from 0% to 100% B from 0-10 minutes, hold 100% B until 17 minutes, linear return to 0% B from 17 to 17.1 minutes, hold 0% B until 20 minutes. The flow rate was 0.45 ml/min, the column temperature was 55°C, and the injection volume was 5 μL. Each sample was analyzed twice, once in positive and once in negative ion mode MS, scan rate 2 spectra/sec, mass range 50-1200 m/z. Source parameters were: drying gas temperature 350°C, drying gas flow rate 10 L/min, nebulizer pressure 30 psig, sheath gas temperature 350°C and flow 11 L/minute, capillary voltage 3500 V, internal reference mass correction enabled. A QC sample run was performed at minimum every tenth injection.

Ion-pair LC-MS samples were analyzed on an identically configured LC-MS system using an Agilent Zorbax Extend C18 1.8 µm RRHD column, 2.1 x 150 mm ID, equipped with a matched guard column. Mobile phase A was 97% water, 3% methanol. Mobile phase B was 100% methanol. Both mobile phases contained 15 mM tributylamine and 10 mM acetic acid. Mobile phase C was 100% acetonitrile. Elution was carried out using a linear gradient followed by a multi-step column wash including automated (valve-controlled) backflushing, detailed as follows: 0-2min, 0%B; 2-11 min, linear ramp from 0-99%B; 12-16 min, 99%B, 16-17.5min, 99-0%B. At 17.55 minutes, the 10-port valve was switched to reverse flow (back-flush) through the column. From 17.55-20.45 min the solvent was ramped from 99%B to 99%C. From 20.45-20.95 min the flow rate was ramped up to 0.8 mL/min, which was held until 22.45 min, then ramped down to 0.6mL/min by 22.65 min. From 22.65-23.45 min the solvent was ramped from 99% to 0% C while flow was simultaneously ramped down from 0.6-0.4mL/min. From 23.45 to 29.35 min the flow was ramped from 0.4 to 0.25mL/min; the 10-port valve was returned to restore forward flow through the column at 25 min. . Column temperature was 35°C and the injection volume was 5 μL. MS acquisition was performed in negative ion mode, scan rate 2 spectra/sec, mass range 50-1200 m/z. Source parameters were: drying gas temperature 250°C, drying gas flow rate 13 L/min, nebulizer pressure 35 psig, sheath gas temp 325°C and flow 12 L/min, capillary voltage 3500V, internal reference mass correction enabled. A QC sample run was performed at minimum every tenth injection.

Iterative MS/MS data was acquired for both reverse phase and ion pairing methods using the pooled sample material to enable compound identification. Eight repeated LC-MS/MS runs of the QC sample were performed at three different collision energies (10, 20, and 40) with iterative MS/MS acquisition enabled. The software excluded precursor ions from MS/MS acquisition within 0.5 minute of their MS/MS acquisition time in prior runs, resulting in deeper MS/MS coverage of lower-abundance precursor ions.

Feature detection and alignment was performed utilizing a hybrid targeted/untargeted approach. Targeted compound detection and relative quantitation was performed by automatic integration followed by manual inspection and correction using Profinder v8.0 (Agilent Technologies, Santa Clara, CA.) Non-targeted feature detection was performed using custom scripts that automate operation of the “find by molecular feature” workflow of the Agilent Masshunter Qualitative

Analysis (v7) software package. Feature alignment and recursive feature detection were performed using Agilent Mass Profiler Pro (v8.0) and Masshunter Qualitative Analysis (“find by formula” workflow), yielding an aligned table including m/z, RT, and peak areas for all features.

Data Cleaning and Degeneracy Removal: The untargeted features and named metabolites were merged to generate a combined feature table. Features missing from over 50% of all samples in a batch or over 30% of QC samples were then removed prior to downstream normalization procedure. Next, the software package Binner was utilized to remove redundancy and degeneracy in the data.^142^ Briefly, Binner first performs RT-based binning, followed by clustering of features by Pearson’s correlation coefficient, and then assigns annotations for isotopes, adducts or in-source fragments by searching for known mass differences between highly correlated features.

Normalization and Quality Control: Data were then normalized using a Systematic Error Removal Using Random Forest (SERRF) approach,^147^ which helps correct for drift in peak intensity over the batch using data from the QC sample runs. When necessary to correct for residual drift, peak area normalization to closest-matching internal standard was also applied to selected compounds. Both SERRF correction and internal standard normalization were implemented in R. Parameters were set to minimize batch effects and other observable drifts, as visualized using principal component analysis score plots of the full dataset. Normalization performance was also validated by examining relative standard deviation values for additional QC samples not included in the drift correction calculations.

Compound identification: Metabolites from the targeted analysis workflow were identified with high confidence (MSI level 1)^148^ by matching retention time (+/- 0.1 minute), mass (+/- 10 ppm) and isotope profile (peak height and spacing) to authentic standards. MS/MS data corresponding to unidentified features of interest from the untargeted analysis were searched against a spectral library (NIST 2020 MS/MS spectral database or other public spectral databases) to generate putative identifications (MSI level 2) or compound-class level annotations (MSI level 3) as described previously.^145^

#### LC-MS/MS untargeted lipidomics

Sample preparation: Non-targeted lipid analysis was conducted at the Georgia Institute of Technology. Powdered tissue samples (10 mg) were extracted in 400 μL isopropanol containing stable isotope-labeled internal standards (IS) with bead homogenization with 2mm zirconium oxide beads (Next Advance) using a TissueLyser II (10min, 30 Hz). Samples were then centrifuged (5 min, 21,100xg), and supernatants were transferred to autosampler vials. Plasma samples (25 μL) were extracted by mixing with 75 μL isopropanol containing the IS mix followed by centrifugation. Sample blanks, pooled extract samples used as quality controls (QC), and consortium reference samples, were prepared for analysis using the same methods. The IS mix consisted of PC (15:0-18:1(d7)), Catalog No. 791637; PE (15:0-18:1(d7)), Catalog No. 791638; PS (15:0-18:1(d7)), Catalog No. 791639; PG(15:0-18:1(d7)), Catalog No. 791640; PI(15:0- 18:1(d7)), Catalog No. 791641; LPC(18:1(d7)), Catalog No. 791643; LPE(18:1(d7)); Catalog No. 791644; Chol Ester (18:1(d7)), Catalog No. 700185; DG(15:0-18:1(d7)), Catalog No. 791647; TG(15:0-18:1(d7)-15:0), Catalog No. 791648; SM(18:1(d9)), Catalog No. 791649; Cholesterol (d7), Catalog No. 700041. All internal standards were purchased from Avanti Polar Lipids (Alabaster, Alabama) and added to the extraction solvent at a final concentration in the 0.1-8 μg/ml range.

Data collection: Lipid LC-MS data were acquired using a Vanquish (ThermoFisher Scientific) chromatograph fitted with a ThermoFisher Scientific AccucoreTM C30 column (2.1 × 150 mm, 2.6 µm particle size), coupled to a high-resolution accurate mass Q-Exactive HF Orbitrap mass spectrometer (ThermoFisher Scientific) for both positive and negative ionization modes. For positive mode analysis, the mobile phases were 40:60 water:acetonitrile with 10 mM ammonium formate and 0.1% formic acid (mobile phase A), and 10:90 acetonitrile:isopropyl alcohol, with 10 mM ammonium formate and 0.1% formic acid (mobile phase B). For negative mode analysis, the mobile phases were 40:60 water:acetonitrile with 10 mM ammonium acetate (mobile phase A), and 10:90 acetonitrile:isopropyl alcohol, with 10 mM ammonium acetate (mobile phase B).

The chromatographic method used for both ionization modes was the following gradient program: 0 minutes 80% A; 1 minute 40% A; 5 minutes 30% A; 5.5 minutes 15% A; 8 minutes 10% A; held 8.2 minutes to 10.5 minutes 0% A; 10.7 minutes 80% A; and held until 12 minutes. The flow rate was set at 0.40 ml/min. The column temperature was set to 50°C, and the injection volume was 2 μL.

For analysis of the organic phase the electrospray ionization source was operated at a vaporizer temperature of 425°C, a spray voltage of 3.0 kV for positive ionization mode and 2.8 kV for negative ionization mode, sheath, auxiliary, and sweep gas flows of 60, 18, and 4 (arbitrary units), respectively, and capillary temperature of 275°C. The instrument acquired full MS data with 240,000 resolution over the 150-2000 m/z range. LC-MS/MS experiments were acquired using a DDA strategy. MS2 spectra were collected with a resolution of 120,000 and the dd-MS2 were collected at a resolution of 30,000 and an isolation window of 0.4 m/z with a loop count of top 7. Stepped normalized collision energies of 10%, 30%, and 50% fragmented selected precursors in the collision cell. Dynamic exclusion was set at 7 seconds and ions with charges greater than 2 were omitted.

Data processing: Data processing steps included peak detection, spectral alignment, grouping of isotopic peaks and adduct ions, drift correction, and gap filling. Compound Discoverer V3.3 (ThermoFisher Scientific) was used to process the raw LC-MS data. Drift correction was performed on each individual feature, where a Systematic Error Removal using Random Forest (SERRF) method builds a model using the pooled QC sample peak areas across the batch and was then used to correct the peak area for that specific feature in the samples. Detected features were filtered with background and QC filters. Features with abundance lower than 5x the background signal in the sample blanks and that were not present in at least 50% of the QC pooled injections with a coefficient of variance (CV) lower than 80% (not drift corrected) and 50% (drift corrected) were removed from the dataset. Lipid annotations were accomplished based on accurate mass and relative isotopic abundances (to assign elemental formula), retention time (to assign lipid class), and MS2 fragmentation pattern matching to local spectral databases built from curated experimental data. Lipid nomenclature followed that described previously.^149^

Quality control procedures: System suitability was assessed prior to the analysis of each batch. A performance baseline for a clean instrument was established before any experiments were conducted. The mass spectrometers were mass calibrated, mass accuracy and mass resolution were checked to be within manufacturer specifications, and signal-to-noise ratios for the suite of IS checked to be at least 75% of the clean baseline values. For LC-MS assays, an IS mix consisting of 12 standards was injected to establish baseline separation parameters for each new column. The performance of the LC gradient was assessed by inspection of the column back pressure trace, which had to be stable within an acceptable range (less than 30% change). Each IS mix component was visually evaluated for chromatographic peak shape, retention time (lower than 0.2 minute drift from baseline values) and FWHM lower than 125% of the baseline measurements. The CV of the average signal intensity and CV of the IS (<=15%) in pooled samples were also checked. These pooled QC samples were used to correct for instrument sensitivity drift over the various batches using a procedure similar to that described by the Human Serum Metabolome (HUSERMET) Consortium. To evaluate the quality of the data for the samples themselves, the IS signals across the batch were monitored, PCA modeling for all samples and features before and after drift correction was conducted, and Pearson correlations calculated between each sample and the median of the QC samples.

#### Metabolomics/Lipidomics Data filtering and normalization

The untargeted metabolomics datasets were categorized as either “named”, for chemical compounds confidently identified, or “unnamed”, for compounds with specific chemical properties but without a standard chemical name. While the preprocessing steps were performed on the named and unnamed portions together, only the named portions were utilized for differential analysis. For each dataset i.e. each assay for each tissue type, the following steps are performed:

- Average rows that have the same metabolite ID.
- Merge the “named” and “unnamed” subparts of the untargeted datasets.
- Convert negative and zero values to NAs.
- Remove features with > 20% missing values.
- Features with < 20% missing values are imputed either using K-Nearest Neighbor imputation (for datasets with > 12 features) or half-minimum imputation (for datasets with < 12 features).
- All data are log2-transformed, and the untargeted data are median-MAD normalized if neither sample medians nor upper quartiles were significantly associated with sex or sex-stratified training group (Kruskal-Wallis p-value < 0.01). Note that for all log2 calculations, 1 is added to each value before log-normalization. This allows metabolite values that fall between 0 and 1 to have a positive log2 value.

Outlier detection was performed by examining the boxplot of each Principal Component and extending its whiskers to the predefined multiplier above and below the interquartile range (5x the IQR). Samples outside this range are flagged. All outliers were reviewed by Metabolomics CAS, and only confirmed technical outliers were removed.

#### Redundant Metabolite/Lipid Management

To address metabolites measured on multiple platforms (e.g. a metabolite measured on the HILIC positive and RP positive platforms), and metabolites with the same corresponding RefMet ID (e.g. alpha-Aminoadipic-acid, Aminoadipic acid both correspond to RefMet name ‘Aminoadipic acid’), we utilize the set of internal standards described above to make a decision on which platform’s measurement of a given feature to include in further analysis. For each tissue, based on whichever platform has the lowest coefficient of variation across all included reference standards for a given refmet id, that metabolite was chosen. The other platforms or metabolites, for this tissue, corresponding to this refmet id were removed from further downstream analysis. Data for all metabolites removed, including information about the coefficient of variation in the standards, normalized data, or differential analysis results, is available in the R Package (see below), but is not loaded by default.

### Statistical analysis

#### Differential analysis

To model the effects of both exercise mode and time, relative to non-exercising control, each measured molecular feature was treated as an outcome in a linear mixed effects model accounting for fixed effects of exercise group (RE, EE, or CON), timepoint, as well as demographic and technical covariates (see covariate selection below for more info). Participant identification was treated as a random effect. For each molecular feature, a cell-means model is fit to estimate the mean of each exercise group-timepoint combination, and all hypothesis tests are done comparing the means of the fixed effect group-timepoint combinations. In order to model the effects of exercise against non-exercising control, a difference-in-changes model was used which compared the change from pre-exercise to a during or post-exercise timepoint in one of the two exercise groups to the same change in control. This effect is sometimes referred to as a “delta-delta” or “difference-in-differences” model.

#### Model selection via simulation

In order to determine the optimal statistical package/model for these data – given MoTrPAC’s unique sampling design (see ‘acute exercise intervention’ above) – a simulation study was conducted to determine the impact of various approaches to account for correlation in repeated measurements. The simulation evaluated type 1 error, power, and bias for multiple plausible modeling approaches, to identify those that would obtain higher power while maintaining nominal type 1 error rates. For a subset of molecular features in the datasets, mean, covariance, skew, and kurtosis were summarized over the relevant timepoints in the acute exercise bout within subgroups defined by sex and exercise type. Then, these summary values were used in combination with the “covsim” R package to generate non-normally distributed simulated data.^150^ Additionally, sample size and missingness patterns were aligned in the simulated data to match those of the observed data. Using a total of 5 million simulated instances where data were generated under the null hypothesis (i.e., no change in the mean values of the outcome over the exercise timepoints) or alternative hypothesis (i.e., the mean value of the outcome differs between at least two timepoints), type 1 error rate and power, respectively, were evaluated for six analytic strategies: (1) pairwise t-tests between timepoints of interest, (2) ordinary least squares regression, (3) differential expression for repeated measures (i.e., dream) with random intercepts,^126^ (4) linear mixed models with random intercepts, (5) mixed models for repeated measures using an unstructured covariance matrix, and (6) generalized linear models using generalized estimating equations (GEEs) and a first degree autoregressive correlation structure. These analyses were replicated for adipose, blood, and muscle samples. The dream function obtained type 1 error rates 5.2, 5.4, and 5.2% for adipose, blood, and muscle analyses, respectively, with higher power than all other methods apart from generalized linear models with GEEs. Although generalized linear models with GEEs had higher power than dream, they also had higher type 1 error with 7.7%, 6.0%, and 6.5% type 1 error rates for adipose, blood, and muscle analyses, respectively. Therefore, due to stability of type 1 error and relatively high power compared to other approaches, the *dream* function from the variancePartition R package was selected for the primary analyses for every omic platform.^126^

#### Covariate selection

Covariates were selected through a combination of a priori knowledge about factors influencing molecular levels as well through empiric screening. Factors were considered for model inclusion by correlating the principal components of each tissue-ome feature set to demographic and technical factors using variancePartition::canCorPairs.^126^ Visualizations and computational analysis that describe this process for the decisions for covariate selection can be found at: https://github.com/MoTrPAC/motrpac-human-presuspension-acute/tree/main/QC. Ultimately, the following fixed effect covariates were included in the models for every omic platform: group and timepoint in a cell means model, age, sex, and BMI, with clinical site additionally included for all platforms except proteomics. ‘Participant id’ was included as a random effect in every model. Each -ome then had ome-specific covariates selected as described in prior sections.

A targeted analysis was conducted to assess whether and how race, ethnicity, and genetic ancestry could be incorporated, given prior evidence that these factors can influence omic data.^151,152^ We evaluated multiple models that included patient-reported race and/or ethnicity, as well as principal components derived from whole-genome sequencing (WGS), to determine their impact on model fit using the BIC. However, no combinations of individual or joint inclusion of race, ethnicity, or WGS-derived principal components led to an improvement in average model BIC across all features for a platform.

The final set of covariates included in each model for each tissue and -ome can be found in the R Package *MotrpacHumanPreSuspensionAnalysis*, and in the MoTrPAC Multi-omic Landscape paper (in submission to *Nature Medicine*).^37^

#### Cell type sensitivity analysis

To evaluate whether shifts in cellular composition confounded the differential expression results, a sensitivity analysis was performed using deconvolution-derived cell-type proportions in blood (see MoTrPAC Multi-omics Landscape paper, in submission to *Nature Medicine*).^37^ Cellular composition was first estimated from whole-blood transcriptomes by deconvolving the bulk RNA-seq profiles with CIBERSORTx (cibersortx.stanford.edu) using the LM22 leukocyte signature matrix with ’Batch Correct’ mode disabled. Because LM22 resolves 22 closely related leukocyte subsets whose marker profiles are highly correlated, the subsets were collapsed by summing their fractions into broader populations: T cells, B cells, NK cells, monocytes, macrophages, dendritic cells, neutrophils, and mast cells. The reliability of these estimates was then assessed by correlating CIBERSORTx proportions at pre-exercise with matched complete blood count values for the same participants, where available. Finally, the blood RNA-seq differential analysis was refit with the coarse cell-type fractions (T cells, B cells, NK cells, monocytes, and neutrophils) added as numeric covariates. Proportions were z-scored before inclusion, and the model otherwise matched the original differential analysis pipeline.

Comparing the refit results against the original model indicated whether accounting for cellular composition altered the estimated exercise effects.

#### Missingness

Data missingness varied for multiple reasons including experimental design (i.e. temporal randomization which randomized some participants to have samples obtained at a subset of timepoints, see assay limitations (metabolomics and proteomics can have missingness as described in their individual sections). For some omic sets, imputation to alleviate missingness could be performed (metabolomics) but for others was not, including proteomics. Analysis demonstrated proteomics missingness was approximately at random (data not shown).

Given the above issues affecting the ultimate sample size for effect estimation, for a feature to be included in the analysis, a minimum number of 3 participants were required to have a paired pre-exercise sample for all groups and all during/post-exercise timepoints. Thus, all group comparisons (e.g. RE vs CON at 3.5 h post-exercise) required 3 participants with matched pre- and post-exercise samples in each group. This requirement was satisfied in all omes and tissues except a subset of MS-acquired proteomics features in SKM and AT. Features not meeting this criterion in any group-timepoint set were excluded from differential analysis entirely, though raw and QC-normalized values are available.

#### Specific omic-level statistical considerations

Models for all omes were fit using ‘variancePartition::dream’.^126^ For the transcriptomics, which are measured as numbers of counts, the mean-variance relationship was measured using ‘variancePartition::voomWithDreamWeights’ as previously described.^126^ For all other omes the normalized values were used directly as input to the statistical model.

The full implementation of the statistical models for each dataset can be found in the R Package *MotrpacHumanPreSuspensionAnalysis*.

#### Contrast types

The specific contrasts made in the statistical analysis include three categories of comparison: difference-in-changes relative to control, group-specific, and resistance vs endurance. All comparisons were structured using ‘variancePartition::makeContrastsDream’.^126^

The difference-in-changes model as described at the opening of this section represents the primary DA analysis, and any individual feature mentioned as statistically significantly changing due to exercise will be referring to the difference-in-changes results unless otherwise specifically stated.

The next category of contrasts is a group-specific comparison, which compares a given post- intervention timepoint to pre-exercise within a given group, without comparing to the control group. This contrast was implemented to more easily quantify effect sizes within each group independently but are not used as a general endpoint.

Finally, the RE vs EE is very similar to the difference-in-differences model, but sets the RE group as a matched control to the EE group. In this comparison, the change in response to exercise at a given timepoint in one exercise group is directly compared to the corresponding change in the other exercise group. This contrast can directly test the hypothesis that the EE effect is different from the RE effect. There are no samples obtained during RE bouts, so those timepoints in blood were excluded from this comparison.

#### Significance thresholds

For each of the above contrasts, p-values were adjusted for multiple comparisons for each unique contrast-group-tissue-ome-timepoint combination separately using the Benjamini- Hochberg method to control False Discovery Rate (FDR).^153^ Features were considered significant at an adjusted p-value < 0.05 unless otherwise stated.

#### Human feature to gene mapping

The feature-to-gene map links each feature tested in differential analysis to a gene, using Ensembl version 105 (mapped to GENCODE 39) as the gene identifier source.^154^ Proteomics feature IDs (UniProt IDs) were mapped to gene symbols and Entrez IDs using UniProt’s mapping files.^155^ Gene symbols, Entrez IDs, and Ensembl IDs were assigned to features using biomaRt version 2.58.2 (Bioconductor 3.18).^156,157^

Metabolite features were mapped to KEGG IDs using KEGGREST, RefMet REST API, or web scraping from the Metabolomics Workbench on August 6, 2026.^158,159^

### Enrichment analysis

#### Preparation of differential analysis results

The differential analysis results tables for each combination of tissue and ome were converted to matrices of z-scores with either gene symbols or RefMet metabolite/lipid IDs as rows and contrasts as columns. These matrices serve as input for the enrichment analyses. For the transcriptomics, transcripts were first mapped to gene symbols. To resolve cases where multiple features mapped to a single gene, only the most extreme z-score for each combination of tissue, contrast, and gene was retained. Metabolite and lipid identifiers were standardized using the Metabolomic Workbench Reference List of Metabolite Names (RefMet) database.

#### Gene set selection

Gene sets were obtained from the MitoCarta3.0 database, CellMarker 2.0 database,^89^ and the C2-CP (excluding KEGG_LEGACY) and C5 collections of the human Molecular Signatures Database (MSigDB; v2023.2.Hs).^89,160,161^ Metabolites and lipids were grouped according to chemical subclasses from the RefMet database.^162^ This includes subclasses such as “Acyl carnitines” and “Saturated fatty acids”.

For each combination of tissue and ome, molecular signatures were filtered to only those genes and metabolites/lipids that appeared in the differential analysis results. After filtering, all molecular signatures were required to contain at least 5 features, with no restriction on the maximum size of sets. Additionally, gene sets were only kept if they retained at least 70% of their original genes to increase the likelihood that the genes that remain in a given set are accurately described by the set label. Gene set enrichment was thus not performed among plasma proteomics given limited membership among the targeted platform.

#### Analysis of Molecular Signatures

Analysis of molecular signatures was carried out with the pre-ranked Correlation Adjusted MEan RAnk (CAMERA-PR) gene set test using the z-score matrices described in the “Preparation of differential analysis results” section to summarize the differential analysis results for each contrast at the level of molecular signatures.^163^ Z-scores were selected as the input statistics primarily to satisfy the normality assumption of CAMERA-PR, and the analysis was carried out with the *cameraPR.matrix* function from the TMSig R/Bioconductor package.^164,165^

#### Over-representation analysis (ORA)

Over-representation analysis (ORA) was conducted using the run_ORA function from the *MotrpacHumanPreSuspensionAnalysis* R package which uses the hypergeometric test for statistical significance. The input consisted of differentially expressed genes (adjusted p-value < 0.05) identified from one or more omic layers in a particular tissue. The background gene set included all genes detected in each of the ome(s) and tissue(s) under consideration.

#### Statistical significance thresholds

For both CAMERA-PR and ORA results, p-values were adjusted within each combination of tissue, ome, contrast, and broad molecular signature collection (MitoCarta3.0, CellMarker 2.0, C2, C5, and RefMet) using the Benjamini-Hochberg method. Gene sets and RefMet subclasses were declared significant if their adjusted p-values were less than 0.05.

#### Visualization methods

Bubble heatmaps were generated from the enrichment analysis results using the *enrichmap* function from the TMSig R/Bioconductor package.^164,165^

#### In silico plasma secretome analyses

Published atlases of human plasma protein tissue and/or cellular sources were used to map the putative location of exercise-responsive plasma proteins. Briefly, these resources used MS proteomic and/or RNA data across 29 tissues and 8 cell types from three separate human studies, EMBL-EBI, GTEx, and HATLAS to infer the tissue source for a given plasma protein; detailed descriptions of these datasets are found here.^85–87^ Protein-tissue relationships were categorized according to: single tissue- or cell-enriched; multiple tissues or cell-types; or common; and subsequently assigned a confidence score (global label score, GLS) based on tissue labels across the three atlases.^85^ Proteins were then assigned to the tissue or cell labels with the highest GLS on a 1-4 scale (or common), with increasing score reflecting increasing confidence in a given tissue assignment. Using these methods, we subsequently mapped 221 of the 223 unique DA plasma proteins in EE and/or RE in MoTrPAC to their annotated tissue source and by GLS. MoTrPAC participants’ adipose and skeletal muscle transcriptional and global proteomic responses to exercise for a given DA plasma protein were then compared to evaluate the plasma protein’s putative tissue source during exercise.

#### Cell deconvolution

We used the program CIBERSORTx^96^ to conduct cell type deconvolution operations. The normalized blood RNAseq data across all control and exercise samples were used as input and the LM22 leukocyte cell type signature matrix was used as the reference.^117^ CIBERSORTx generated as output a predicted cell type composition for each sample. Mean cell type proportion was displayed for each cell type across all pre-exercise samples to present a baseline estimate for cell type composition. Statistical significance of differences in cell type composition for each cell type at each timepoint within each exercise mode and control were calculated using the linear model R function *lm* with the formula: Cell.Type ∼ group_timepoint + Sex + calculatedAge + BMI + codedsiteid for each cell type. P-values were adjusted using the R function *p.adjust* with the Benjamini Hochberg (BH) method.

#### Canonical Correlation Analysis

To identify metabolite signatures associated with variation across cardiorespiratory fitness, muscle strength, and metabolic phenotypes, we applied sparse CCA to baseline plasma metabolomic profiles and exercise-related clinical traits. This multivariate approach identifies shared axes of variation between two high-dimensional datasets by maximizing the correlation between linear combinations of features from each domain.^58^ Unlike principal component analysis or traditional regression, CCA jointly optimizes associations between domains without predefining one as explanatory and the other as outcome.

We used pre-exercise plasma metabolomics data and a curated set of exercise phenotypes spanning aerobic fitness (e.g. VO_₂_peak, O_₂_ pulse, ventilatory threshold), hemodynamic responses (e.g. exercise systolic/diastolic blood pressure), and muscular strength (e.g. handgrip and isokinetic torque). Sparse penalized CCA was implemented using the *PMA* R package, with optimal regularization parameters for each domain selected via 1,000 permutations. The final model retained the top three canonical variates for interpretation. Canonical weights were visualized using clustered barplots to highlight metabolite-trait associations for each variate.

Each canonical variate was visualized based on the strongest exercise trait loadings and the associated top metabolite contributors.

#### Association analysis between molecular features and clinical phenotypes

For each molecular feature (plasma metabolite or skeletal muscle transcript), a linear regression model was fit (using the lm() function in the stats package in R)^165^ to test the association between pre-exercise molecular abundance and the clinical variable of interest. Models required a minimum of 5 residual degrees of freedom for stable estimation (i.e., n > p + 5, where n is the number of samples and p is the number of predictors). The outcome variable was feature abundance, and the primary independent variable was the clinical trait, adjusted for age, sex, and BMI. Subsequently, the estimated regression coefficient, standard error, t-statistic, nominal p-value, and number of observations for each feature were extracted for each clinical trait.

Multiple testing correction was performed within each tissue-ome-trait combination using the Benjamini-Hochberg false discovery rate method.

Pre-exercise VO_2_peak (ml O_2_*kg body mass^-1^min^-1^) results were integrated with DA plasma metabolites with acute exercise by first restricting to features with an adjusted p-value < 0.05, and then classifying exercise-responsive features as described in ‘Contrast types’ above. All analyses were performed in R version 4.4.1.

### QUANTIFICATION AND STATISTICAL ANALYSIS

#### Statistical parameter reporting

All relevant statistical parameters for the analyses performed in this study such as sample size (*N*), center and spread (e.g. mean/median, standard deviation/error), statistical methodology (e.g. mixed-effects linear model) and significance cutoffs (e.g. adjusted *p*-value < 0.05) are reported in the main text, figure legends, methods, and/or supplementary information. Where appropriate, methods used to determine the validity of certain statistical assumptions are discussed.

#### Statistical limitations

This report presents analyses and results for 175 participants randomized before the suspension of the MoTrPAC study due to the COVID-19 pandemic. The main post-suspension MoTrPAC study will include over 1,500 participants randomized under a slightly modified protocol.^36^ The current paper focuses on evaluating feasibility and generating hypotheses for the main study. Results should be interpreted with caution for several reasons: 1) small sample sizes reduce the power to detect even moderate effects;^166^ 2) simple randomization of small groups can create imbalances in both known and unknown confounders;^167,168^ and 3) NIH initiatives have long emphasized caution in interpreting small studies to enhance reproducibility.^169^

Although blocked randomization with site-based stratification was employed, discrepancies in regulatory approvals, start-up times, and pandemic-related interruptions resulted in imbalanced and small sample sizes. Specific limitations in the baseline (pre-intervention) data include: 1) subgroup sample sizes based on intervention group, sex, and timepoint as small as 4 participants; 2) limited generalizability, as 50% of controls with biosamples were randomized at two of the ten sites, and 74% of participants at four sites; and 3) "sex differences" may reflect "body composition differences" due to insufficient data to disentangle sex from body composition, and variations in DXA machine operators and brands across sites.

Testing for heterogeneity of response in small subgroups was largely avoided. As Brookes et al. show,^170^ interaction effects must be at least double the main effect to achieve 80% power. While the EE and RE groups each had around 70 participants with biosamples, providing 80% power to detect a 0.5 effect size (difference in means/SD) using a two-sample t-test (alpha = 0.05, two-sided), for tests of interaction effects to have 80% power an interaction effect size > 1 would be required, which is considered large.^171^ Consequently, heterogeneity of response was explored only descriptively within the larger randomized groups (EE/RE), with some inferential statistics (e.g., confidence intervals, p-values) emphasizing interval estimation. Our aim was to present the results from a hypothesis-generating perspective, following Ioannidis’s cautionary guidance,^166^ and to lay a solid foundation for future analyses in the main MoTrPAC study.

### Additional resources

#### Package assembly and distribution

The *MotrpacHumanPreSuspensionAnalysis* R package (https://github.com/MoTrPAC/MotrpacHumanPreSuspensionAnalysis) contains functions to generate visualizations, as well as data objects for differential abundance analysis, summary statistics for normalized expression data, feature to gene mapping, and molecular signature datasets as described in the methods.

The *MotrpacHumanPreSuspensionData* R package (https://github.com/MoTrPAC/MotrpacHumanPreSuspensionData) is a package containing individual level data, which cannot be made openly available and can be accessed with approval from a data access committee. See motrpac-data.org for details.

Data used in the preparation of this article were obtained from the Molecular Transducers of Physical Activity Consortium (MoTrPAC) database, which is available for public access at motrpac-data.org. The specific collection versions are: quant-id/human-precovid/c1.0, analysis/human-precovid-sed-adu/c2.0, phenotype/humanprecovid-sed-adu/c3.0.

Lastly, the MotrpacPreSuspensionAcute GitHub repository (https://github.com/MoTrPAC/motrpac-human-presuspension-acute) contains code and individual parameters for each figure panel, utilizing the MotrpacHumanPreSuspensionAnalysis and MotrpacHumanPreSuspensionData packages.

## Acknowledgements

The MoTrPAC Study is supported by NIH grants U24OD026629 (Bioinformatics Center), U24DK112349, U24DK112342, U24DK112340, U24DK112341, U24DK112326, U24DK112331, U24DK112348 (Chemical Analysis Sites), U01AR071133, U01AR071130, U01AR071124, U01AR071128, U01AR071150, U01AR071160, U01AR071158 (Clinical Centers), U24AR071113 (Consortium Coordinating Center), U01AG055133, U01AG055137, U01AG055135, U01AG070959, U01AG070960, and U01AG070928 (Pre-Clinical Animal Sites).

JMR is supported by NIH K23 HL150327; R03OD038387. MEL is supported by Wu Tsai Human Performance Alliance.

## Author information

### MoTrPAC Study Group

**Bioinformatics Center:** David Amar, Trevor Hastie, David Jimenez-Morales, Daniel H. Katz, Malene E. Lindholm, Samuel Montalvo, Robert Tibshirani, Jay Yu, Jimmy Zhen, Euan A. Ashley, Matthew T. Wheeler

**Biospecimens Repository:** Sandra T. May, Jessica L. Rooney, Russell Tracy

**Data Management, Analysis, and Quality Control Center:** Catherine Gervais, Fang-Chi Hsu, Byron C. Jaeger, David Popoli, Joseph Rigdon, Courtney G. Simmons, Cynthia L. Stowe, Michael E. Miller

**Exercise Intervention Core:** W. Jack Rejeski

**NIH:** Ashley Y. Xia

**Preclinical Animal Study Sites:** Sue C. Bodine, R. Scott Rector

**Chemical Analysis Sites:** Hiba Abou Assi, Mary Anne S. Amper, Brian J. Andonian, Isaac K. Attah, Jacob L. Barber, Kevin Bonanno, Clarisa Chavez Martinez, Natalie M. Clark, Johanna Y. Fleischman, David A. Gaul, Yongchao Ge, Marina A. Gritsenko, Joshua R. Hansen, Patrick Hart, Zhenxin Hou, Chelsea M. Hutchinson-Bunch, Olga Ilkayeva, Gayatri Iyer, Pierre M. Jean-Beltran, Christopher A. Jin, Maureen T. Kachman, Hasmik Keshishian, Damon T. Leach, Minghui Lu, D. R. Mani, Gina M. Many, Nada Marjanovic, Nikhil Milind, Matthew E. Monroe, Ronald J. Moore, Venugopalan D. Nair, German Nudelman, Nora-Lovette Okwara, Vladislav A.

Petyuk, Paul D. Piehowski, Hanna Pincas, Wei-Jun Qian, Prashant Rao, Abraham Raskind, Alexander Raskind, Stas Rirak, Jeremy M. Robbins, Margaret Robinson, Tyler J. Sagendorf, James A. Sanford, Gregory R. Smith, Kevin S. Smith, Yifei Sun, Gaurav Tiwari, Mital Vasoya, Nikolai G. Vetr, Alexandria Vornholt, Yilin Xie, Xuechen Yu, Elena Zaslavsky, Zidong Zhang, Bingqing Zhao, Joshua N. Adkins, Charles F. Burant, Steven A. Carr, Clary B. Clish, Facundo M. Fernandez, Robert E. Gerszten, Stephen B. Montgomery, Christopher B. Newgard, Eric A. Ortlund, Stuart C. Sealfon, Michael P. Snyder, Martin J. Walsh

**Clinical Sites:** Cheehoon Ahn, Alicia Belangee, Bryan C. Bergman, Daniel H. Bessesen, Gerard A. Boyd, Anna R. Brandt, Nicholas T. Broskey, Toby L. Chambers, Clarisa Chavez Martinez, Maria Chikina, Alex Claiborne, Zachary S. Clayton, Paul M. Coen, Katherine A. Collins-Bennett, Tiffany M. Cortes, Brian J. Coyne, Gary R. Cutter, Matthew Douglass, Daniel E. Forman, Will A. Fountain, Aaron H. Gouw, Kevin J. Gries, Fadia Haddad, Joseph A. Houmard, Kim M. Huffman, Ryan P. Hughes, John M. Jakicic, Catherine M. Jankowski, Neil M. Johannsen, Johanna L. Johnson, Erin E. Kershaw, Dillon J. Kuszmaul, Bridget Lester, Colleen E. Lynch, Edward L. Melanson, Cristhian Montenegro, Kerrie L. Moreau, Masatoshi Naruse, Bradley C Nindl, Tuomo Rankinen, Ulrika Raue, Ethan Robbins, Kaitlyn R. Rogers, Renee J. Rogers, Irene E. Schauer, Robert S. Schwartz, Chad M. Skiles, Lauren M. Sparks, Guillaume Spielmann, Maja Stefanovic-Racic, Andrew M. Stroh, Kristen J. Sutton, Anna Thalacker-Mercer, Todd A. Trappe, Caroline S. Vincenty, Elena Volpi, Katie L. Whytock, Gilhyeon Yoon, Thomas W. Buford, Dan M. Cooper, Sara E. Espinoza, Bret H. Goodpaster, Wendy M. Kohrt, William E. Kraus, Nicolas Musi, Shlomit Radom-Aizik, Blake B. Rasmussen, Eric Ravussin, Scott Trappe

### MoTrPAC Acknowledgements

Nicole Adams, Abdalla Ahmed, Andrea Anderson, Carter Asef, Arianne Aslamy, Marcas M. Bamman, Jerry Barnes, Susan Barr, Kelsey Belski, Will Bennett, Kevin Bonanno, Amanda Boyce, Brandon Bukas, Emily Carifi, Chih-Yu Chen, Haiying Chen, Shyh-Huei Chen, Maria Chikina, Samuel Cohen, Audrey Collins, Gavin Connolly, Elaine Cornell, Julia Dauberger, Carola Ekelund, Shannon S. Emilson, Karyn A. Esser, Jerome Fleg, Nicole Gagne, Mary- Catherine George, Ellie Gibbons, Jillian Gillespie, Laurie Goodyear, Aditi Goyal, Bruce Graham, Xueyun Gulbin, Jere Hamilton, Leora Henkin, Andrew Hepler, Andrea Hevener, Olga Ilkayeva, Lidija Ivic, Ronald Jackson, Andrew Jones, Lyndon Joseph, Leslie Kelly, Ian Lanza, Gary Lee, Jun Li, Adrian Loubriel, Kristal M. Maner-Smith, Ryan Martin, Padma Maruvada, Alyssa Mathews, Curtis McGinity, Lucas Medsker, Kiril Minchev, Samuel G. Moore, Michael Muehlbauer, K Sreekumaran Nair, Anne Newman, John Nichols, Concepcion R. Nierras, George Papanicolaou, Lorrie Penry, Vladislav A. Petyuk, June Pierce, David Popoli, Megan Reaves, Eric W. Reynolds, Jeremy Rogers, Scott Rushing, Santiago Saldana, Rohan Shah, Samiya M. Shimly, Cris Slentz, Deanna Spaw, Debbie Steinberg, Suchitra Sudarshan, Alyssa Sudnick, Jennifer W. Talton, Christy Tebsherani, Nevyana Todorova, Mark Viggars, Jennifer Walker, Michael P. Walkup, Anthony Weakland, Gary Weaver, Christopher Webb, Sawyer Welden, John P. Williams, Marilyn Williams, Leslie Willis, Yi Zhang

### Disclosures

Euan A. Ashley is Founder: Personalis, Deepcell, Svexa, Saturnus Bio, Swift Bio. Founder Advisor: Candela, Parameter Health. Advisor: Pacific Biosciences. Non-executive director: AstraZeneca, Dexcom. Publicly traded stock: Personalis, Pacific Biosciences, AstraZeneca. Collaborative support in kind: Illumina, Pacific Biosciences, Oxford Nanopore, Cache, Cellsonics. Stephen A. Carr is on the scientific advisory boards of PrognomIQ, MOBILion Systems, Kymera, and Stand Up2 Cancer. Pierre Jean Beltran is currently an employee at Pfizer, Inc. Bret H. Goodpaster has served as a member of scientific advisory boards. Byron Jaeger receives consulting fees from Perisphere Real World Evidence, LLC, unrelated to this project. Stephen B. Montgomery is a member of the scientific advisory board for PhiTech, MyOme and Valinor Therapeutics. Gary R. Cutter is a part of: Data and Safety Monitoring Boards: Applied Therapeutics, AI therapeutics, Amgen-NMO peds, AMO Pharma, Argenx, Astra-Zeneca, Bristol Meyers Squibb, CSL Behring, DiamedicaTherapeutics, Horizon Pharmaceuticals, Immunic, Inhrbx-sanfofi, Karuna Therapeutics, Kezar Life Sciences, Medtronic, Merck, Meiji Seika Pharma, Mitsubishi Tanabe Pharma Holdings, Prothena Biosciences, Novartis, Pipeline Therapeutics (Contineum), Regeneron, Sanofi-Aventis, Teva Pharmaceuticals, United BioSource LLC, University of Texas Southwestern, Zenas Biopharmaceuticals. Consulting or Advisory Boards: Alexion, Antisense Therapeutics/Percheron, Avotres, Biogen, Clene Nanomedicine, Clinical Trial Solutions LLC, Endra Life Sciences, Genzyme, Genentech, Immunic, Klein-Buendel Incorporated, Kyverna Therapeutics, Inc., Linical, Merck/Serono, Noema, Neurogenesis, Perception Neurosciences, Protalix Biotherapeutics, Regeneron, Revelstone Consulting, Roche, Sapience Therapeutics, Tenmile. Dr. Cutter is employed by the University of Alabama at Birmingham and President of Pythagoras, Inc. a private consulting company located in Birmingham AL. John M. Jakicic is on the Scientific Advisory Board for Wondr Health, Inc. Erin E. Kershaw is a consultant for NodThera and Sparrow Pharmaceuticals and a site PI for clinical trials for Arrowhead Pharmaceuticals. Bradly. C. Nindl is member of the Science Advisory Council, Institute of Human and Machine Cognition, Pensacola, FL. Jeremy M. Robbins is a consultant for Edwards Lifesciences; Abbott Laboratories; Janssen Pharmaceuticals. Renee Rogers is a scientific advisor to AstraZeneca, Neurocrine Biosciences, and the American Council on Exercise, and a consultant to Wonder Health, Inc. and Seca. Stuart C. Sealfon is a founder of GNOMX Corp, leads its scientific advisory board and serves as its temporary Chief Scientific Officer. Michael P. Snyder is a cofounder and shareholder of January AI. Alexandria Vornholt is a consultant for GNOMX Corp.

Disclaimer: The content of this manuscript is solely the responsibility of the authors and does not necessarily represent the views of the National Institutes of Health, or the United States Department of Health and Human Services.

**Table S3. Plasma metabolites that change in both exercise and control groups**

**Table S4. Loadings for top metabolites and phenotypes contributing to canonical variates 1-3**

**Table S5. Differential plasma metabolite changes according to VO_2_peak**

**Table S6. Relationship between plasma metabolite changes and skeletal muscle RNA associated with VO_2_peak**

**Table S7. Differentially abundant plasma protein tissue sources according to HATLAS resource**

---

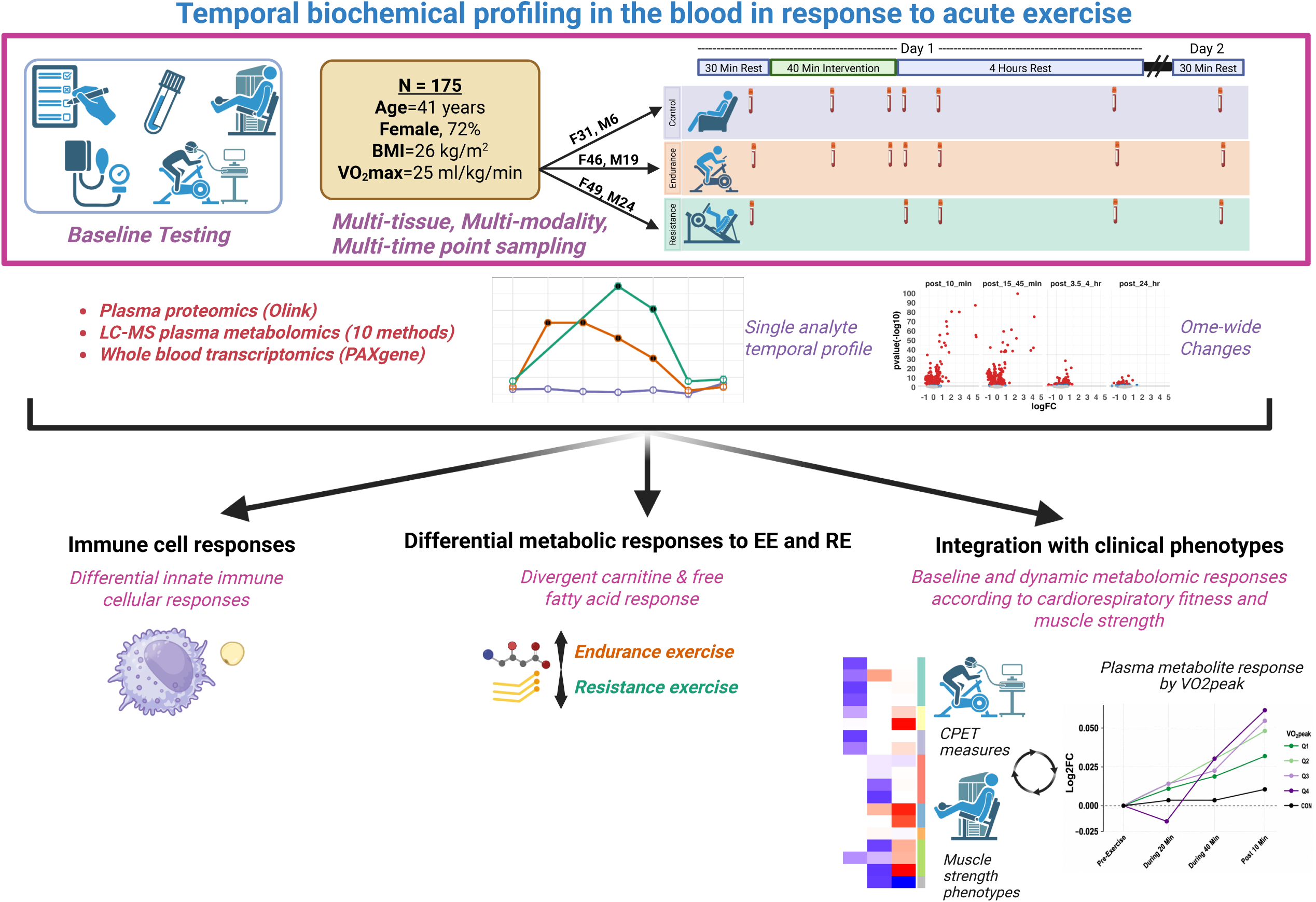

**Figure S1.**
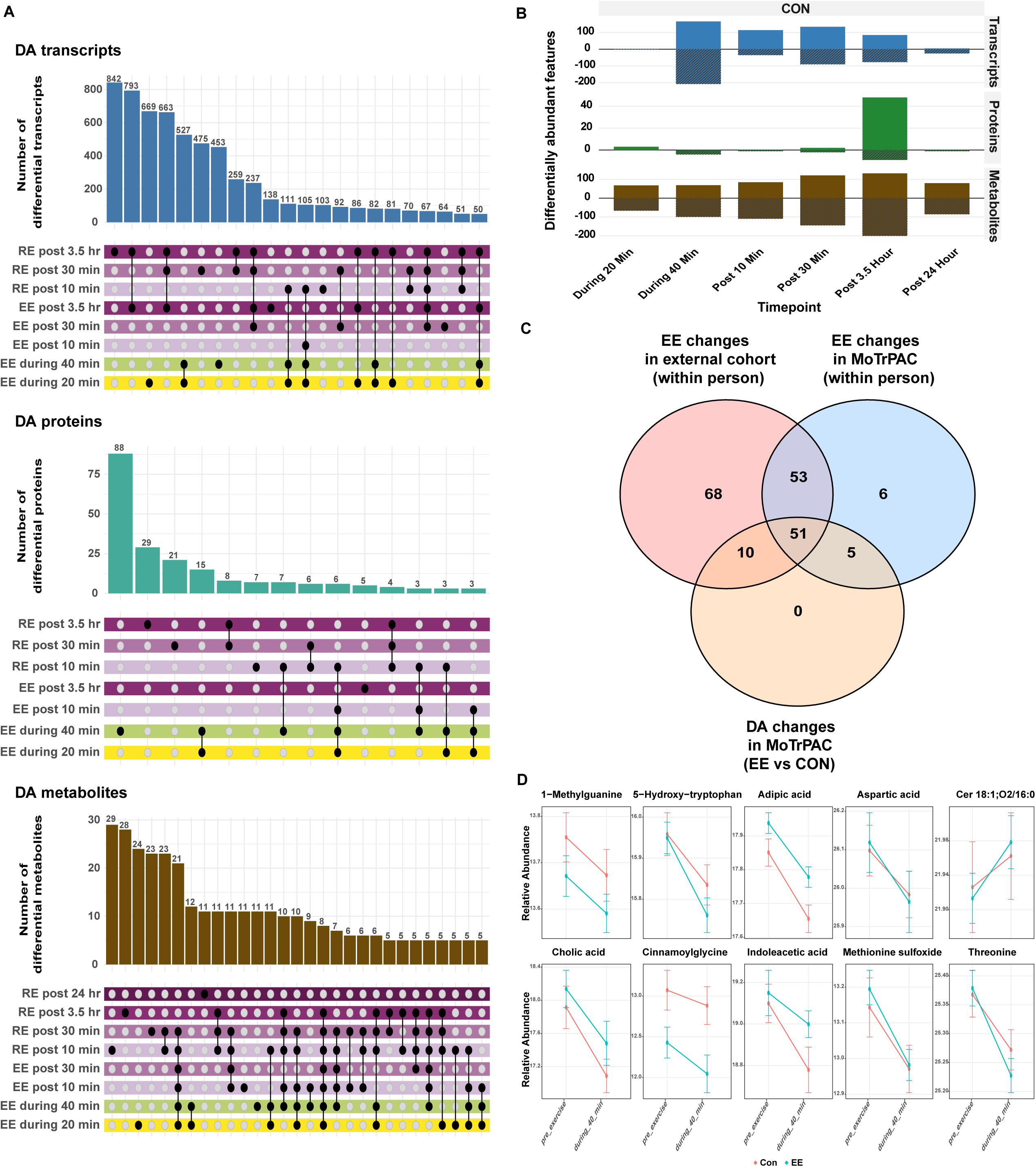
Blood biochemical changes in response to acute endurance and resistance exercise according to blood biochemical profiling platform and in non-exercise controls. (A) UpSet plots showing differentially abundant (DA) features for each blood or plasma biochemical profiling method across all EE and RE timepoints. Black dots indicate the plotted group. A connected line between black dots indicates that the plotted bar represents features shared between those groups. The UpSet plot is ordered by the greatest number of features. (B) Histogram showing molecular changes in the non-exercise controls at each timepoint in each ome (p-adj <0.05). (C) Venn diagram comparing metabolites identified as ‘responsive’ to EE during 40 mins i) in the primary DA analysis, ii) in the within-participant analysis (without accounting for the control arm), and iii) metabolites reported as exercise-responsive in an independent study of 411 individuals (without a control arm^31^). Fifty-two metabolites were identified as exercise responsive by both the within-participant MoTrPAC analysis and the independent study but were no longer significant after accounting for temporal changes observed in the control group, indicating that their apparent responses are not specific to exercise. (D) Representative metabolite trajectories for select metabolites reclassified following inclusion of the non-exercise control arm. Although metabolites appear to change in response to EE in a within-participant analysis in MoTrPAC and an external cohort^31^, similar changes occurred in MoTrPAC CON participants, resulting in no significant exercise-specific effect. Lines represent mean relative abundance in the exercise (red) and control (blue) groups; error bars indicate SEM.

**Figure S2.**
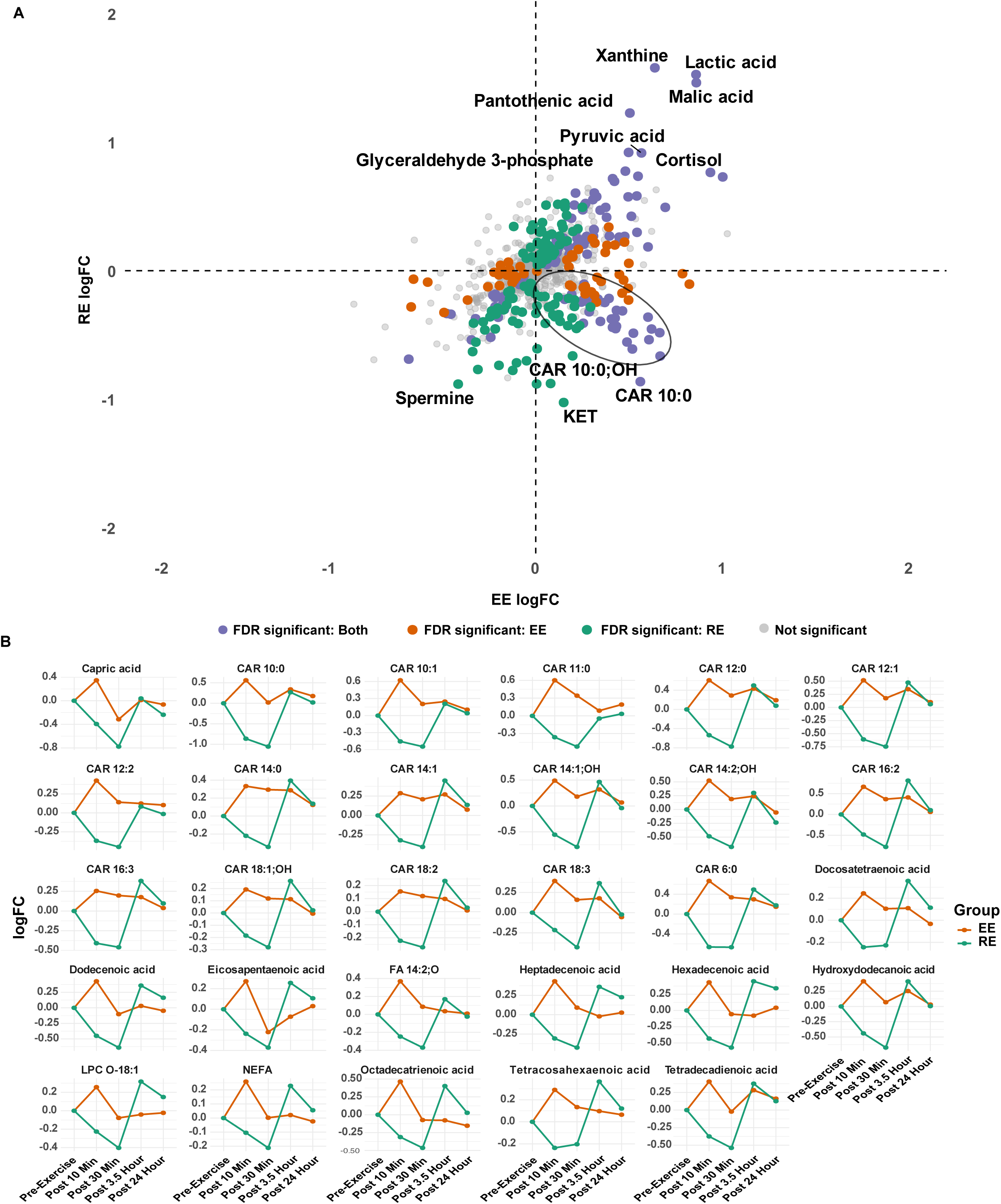
Differential plasma metabolomic responses to EE and RE. (A) Metabolite changes at post 10 min between EE vs CON (log fold-change on X-axis) and RE vs CON (log fold-change on y-axis). (B) Exercise mode-divergent metabolites (circled metabolites in (A) that are FDR significant for EE and RE) highlighting plasma medium- and long-chain acylcarnitine and fatty acid responses (orange lines reflect EE response, green lines reflect RE responses).

**Figure S3.**
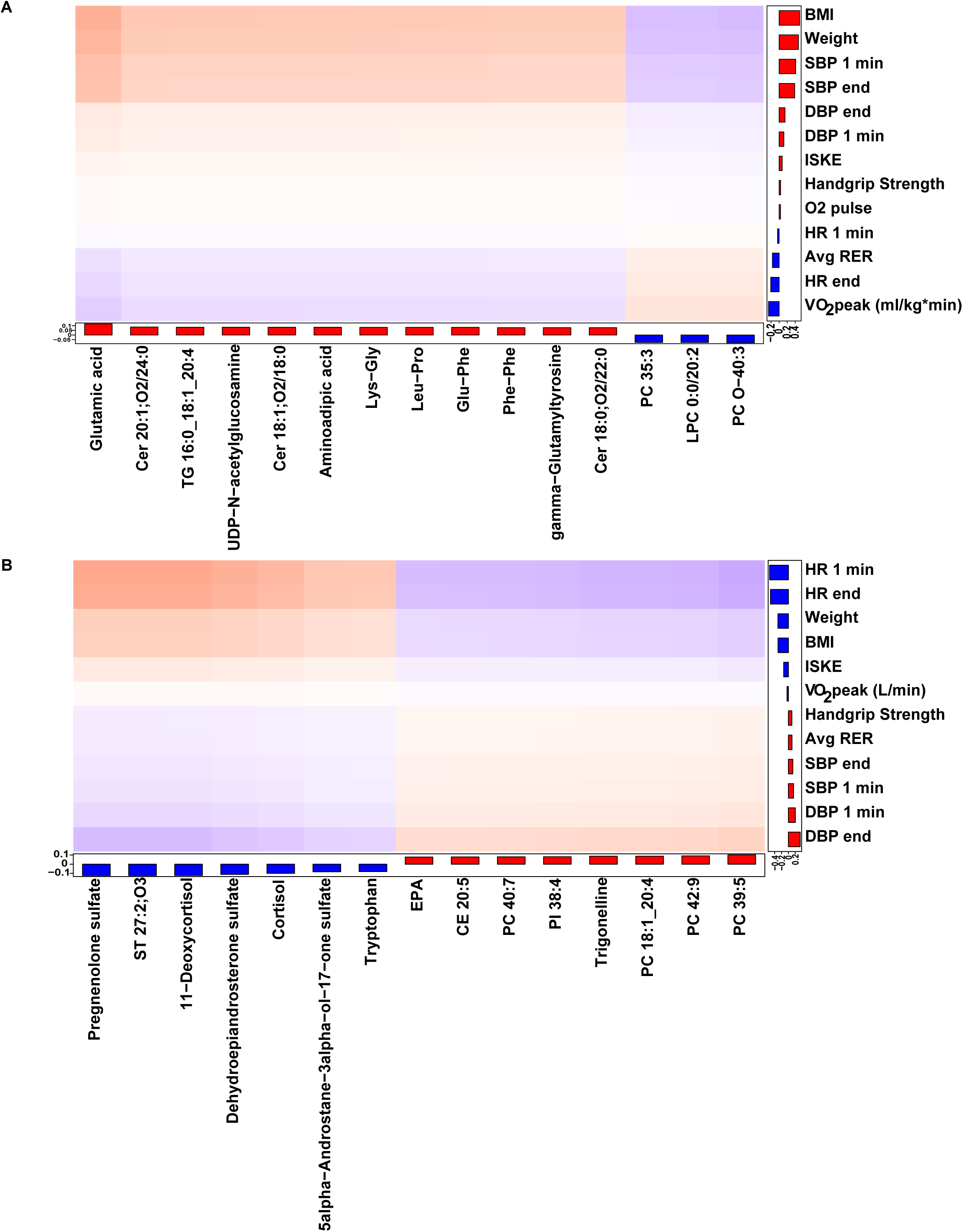
Canonical Variates 2 and 3 from CCA of Plasma Metabolites and Clinical Traits. (A-B) Loadings for metabolites and phenotypes contributing to canonical variate (CV2) (A) and CV3 (B). Bar height reflects the magnitude of contribution to the respective variate. Red bars denote positive loadings; blue bars denote negative loadings. CV1 = metabolite canonical variate 1; CV2 = metabolite canonical variate 2; CV3 = metabolite canonical variate 3; end Watts = end Watts at termination; handgrip strength = average handgrip strength; peak torque = average peak torque from isokinetic test; avg VCO2 = average VCO2 during CPET; avg VE = average ventilation during CPET; DBP end = end diastolic blood pressure; SBP end = end systolic blood pressure; DBP 1 min = diastolic blood pressure at 1 minute recovery; SBP 1 min = systolic blood pressure at 1 minute recovery; HR end = end heart rate; HR 1 min = heart rate at 1 minute recovery; avg RER = average respiratory exchange ratio during CPET; O2 pulse= oxygen pulse (VO_2_/heart rate)

**Figure S4.**
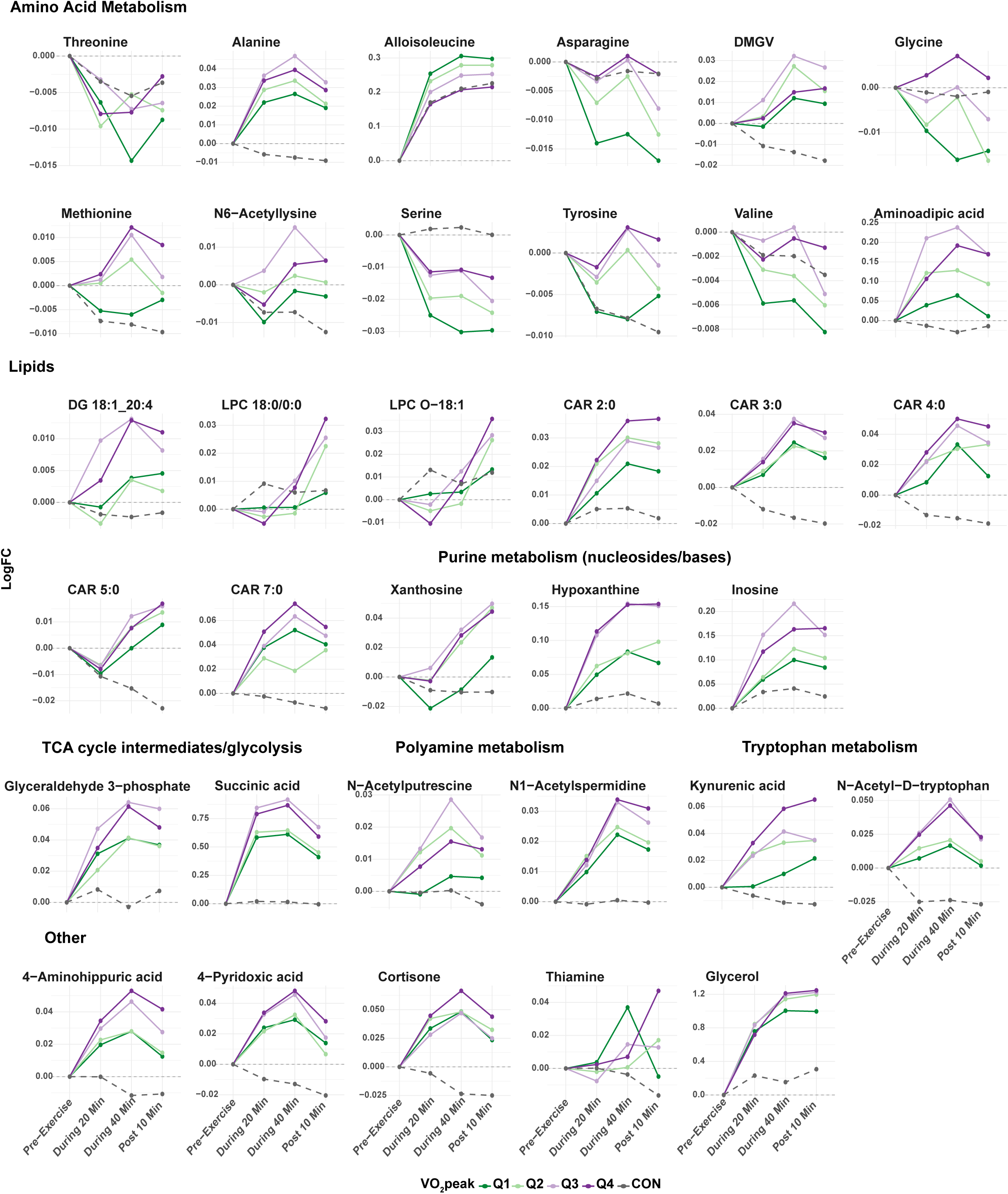
VO_2_peak-associated plasma metabolite changes and skeletal muscle transcriptional expression. Metabolites exhibiting a significant interaction between baseline VO_2_peak and EE change at the post-10 min timepoint. Representative trajectories are shown after stratifying participants into quartiles of baseline VO_2_peak (Q1–Q4). The dashed grey line represents CON. Values are presented as log_2_ fold change relative to baseline. Metabolites shown include representatives from pathways involved in lipid metabolism, tricarboxylic acid cycle intermediates, glycolysis, amino acid and nucleotide metabolism.

**Figure S5.**
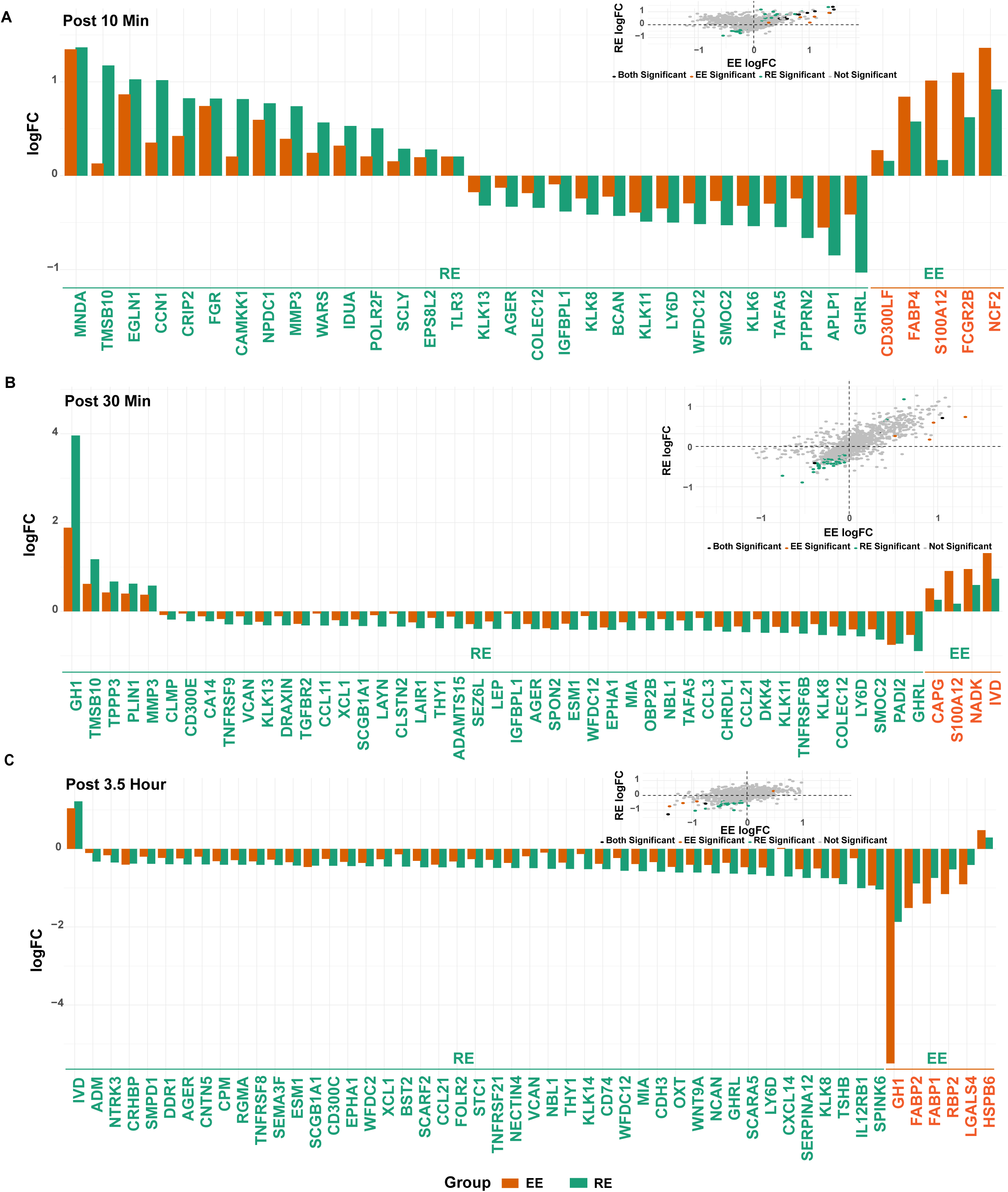
Comparison of resistance versus endurance exercise on plasma proteins. (A-C) Four-quadrant plots displaying log fold-change of DA proteins in EE (x-axis) and RE (y-axis) at (A) post 10 min, (B) post 30 min, and (C) post 3.5 h timepoints. Corresponding bar charts with DA features in each group (RE, green arrows; EE, orange arrows) that compare the magnitude of exercise effect between modes. Three DA proteins at post 24 h not shown.

**Figure S6.**
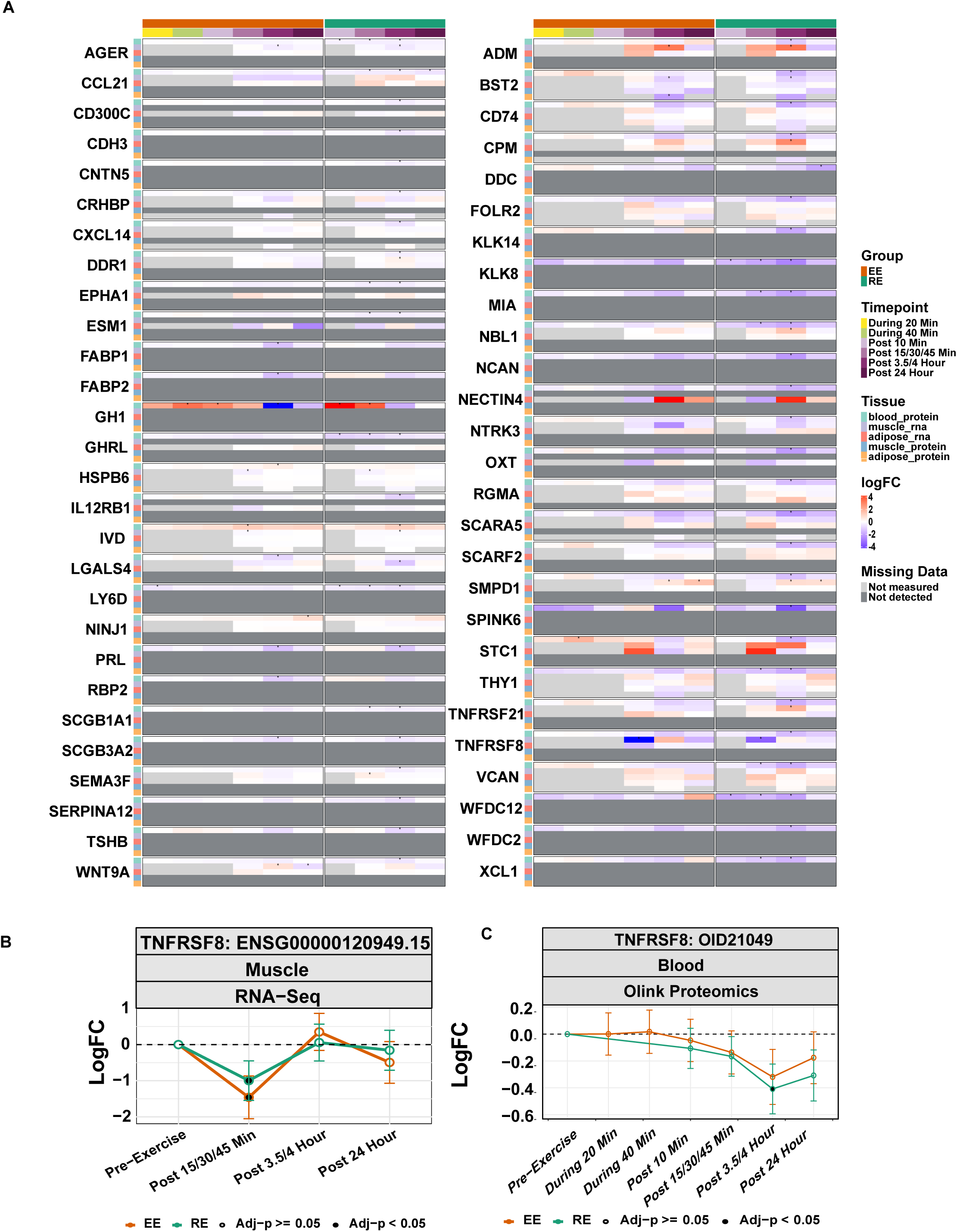
Tissue sources of plasma proteins that change in late exercise recovery. (A-B) Heatmap of all DA plasma proteins at the 3.5 h and/or 24 h timepoints show plasma protein changes, and skeletal muscle and adipose mRNA and global proteomic changes in EE and RE in MoTrPAC: *p-adj <0.05 (A). Light grey boxes reflect timepoints not sampled. Dark grey boxes reflect features not detected. Red arrow highlighting that TNFRSF8 (CD30), a protein annotated to adipose tissue, shows no change in adipose mRNA expression during exercise but (B) decreased skeletal muscle mRNA expression after RE and EE with a subsequent decrease and trend towards decrease in plasma protein levels after RE and EE, respectively. Points with black dots indicate significance (p-adj <0.05) by linear mixed effects model. Error bars indicate 95% confidence intervals.

**Figure S7.**
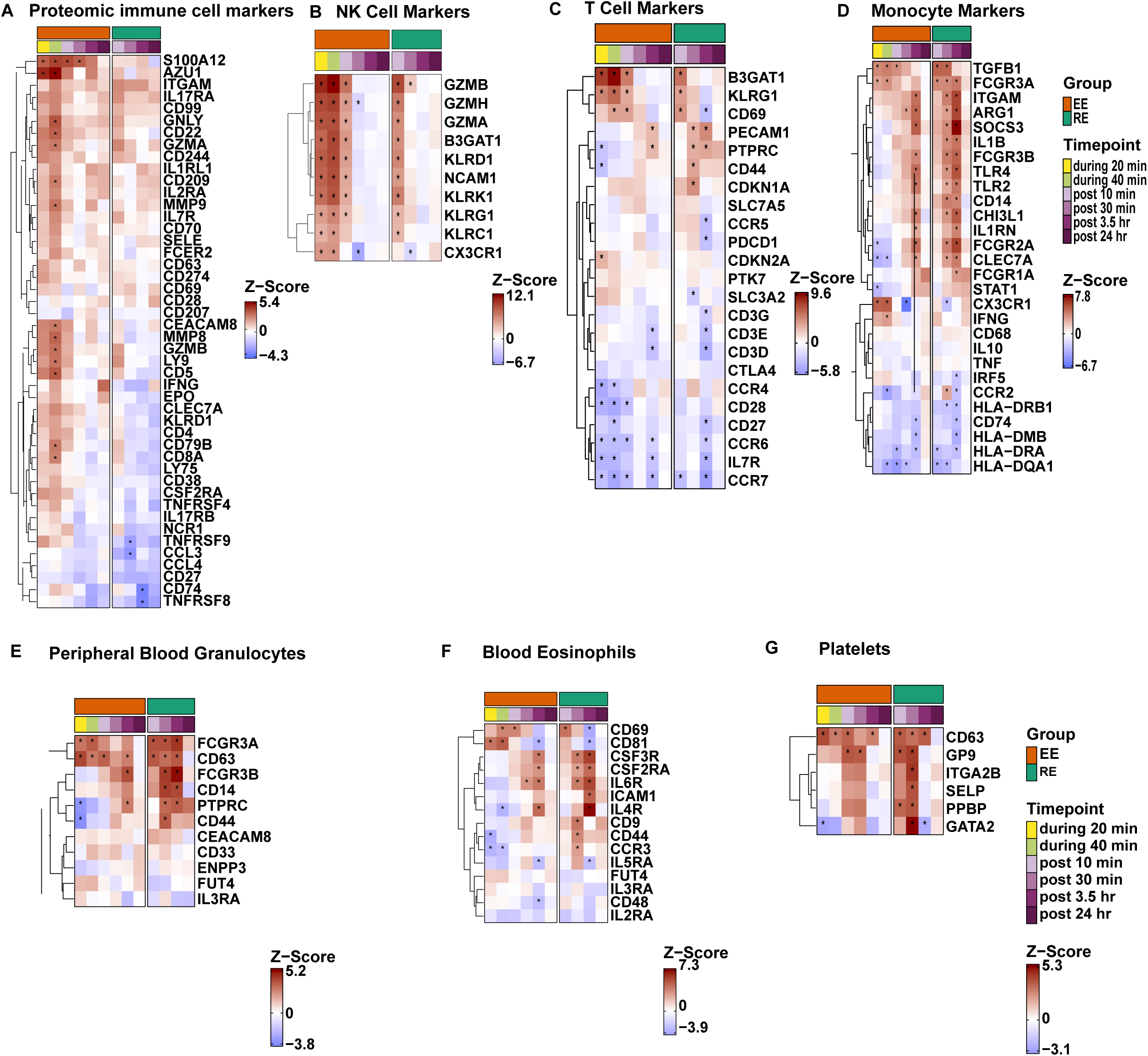
Feature level immune cell enrichments displaying mode-specific regulation. (A) Heatmap displaying Z-scores of select plasma proteomic lymphocyte markers reflects concordance in cytotoxic cell markers at the whole-blood transcriptomic and plasma proteomic levels; an * denotes significance at p-adj <0.05 by linear mixed effects model. (B-G) Heatmaps depicting z-scores of differential abundance of manually curated transcriptomic markers of: (B) NK cells, (C) T cells and (D) monocytes and in CellMarker 2.0 for (E) granulocyte, (F) eosinophil, and (G) platelet human blood transcriptomic pathway enrichments according to exercise mode and timepoint for blood. In (E) FCGR3A and FCGR3B represent CD16A and CD16B, respectively. Select MHC Class II genes by select HLA subclasses, with CD11B denoted as ITGAM. In all figures, an * denotes significance at p-adj <0.05 by linear mixed effects model.

**Figure S8.**
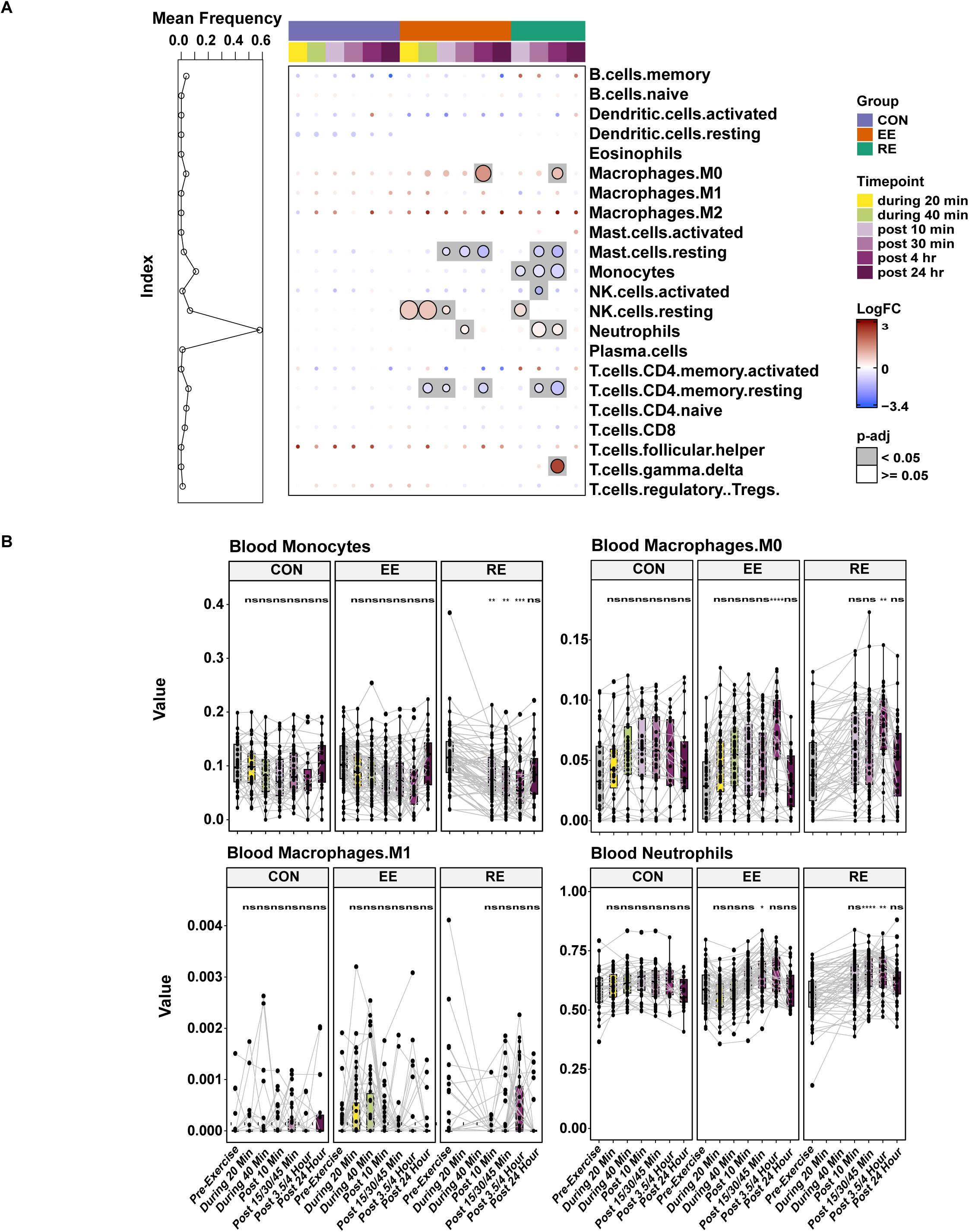
Immune cell responses to acute exercise according to cellular deconvolution. (A) Left square displays the estimated percentage (mean proportion) of each cell type in the blood across groups in the sedentary state, with neutrophils representing the largest cell type in the peripheral blood. Right side bubble heatmap displays log2 fold-change (color) of each estimated cell type in the blood in each of the post-exercise timepoints and in time-matched controls, as well as the adjusted p-value significance of that change (size of bubble). (B) Temporal changes in select cell type proportions are plotted to display individual changes in average log2 fold-change trajectories through all pre-, during and post- exercise timepoints; here a line is used to connect individual subjects. Significance in difference in cell type proportion is measured through a linear model. *p-adj <0.05; ** p-adj <0.005; *** p-adj <5e-4; ****p-adj <5e-5; ns: non-significant.

**Table S1.**
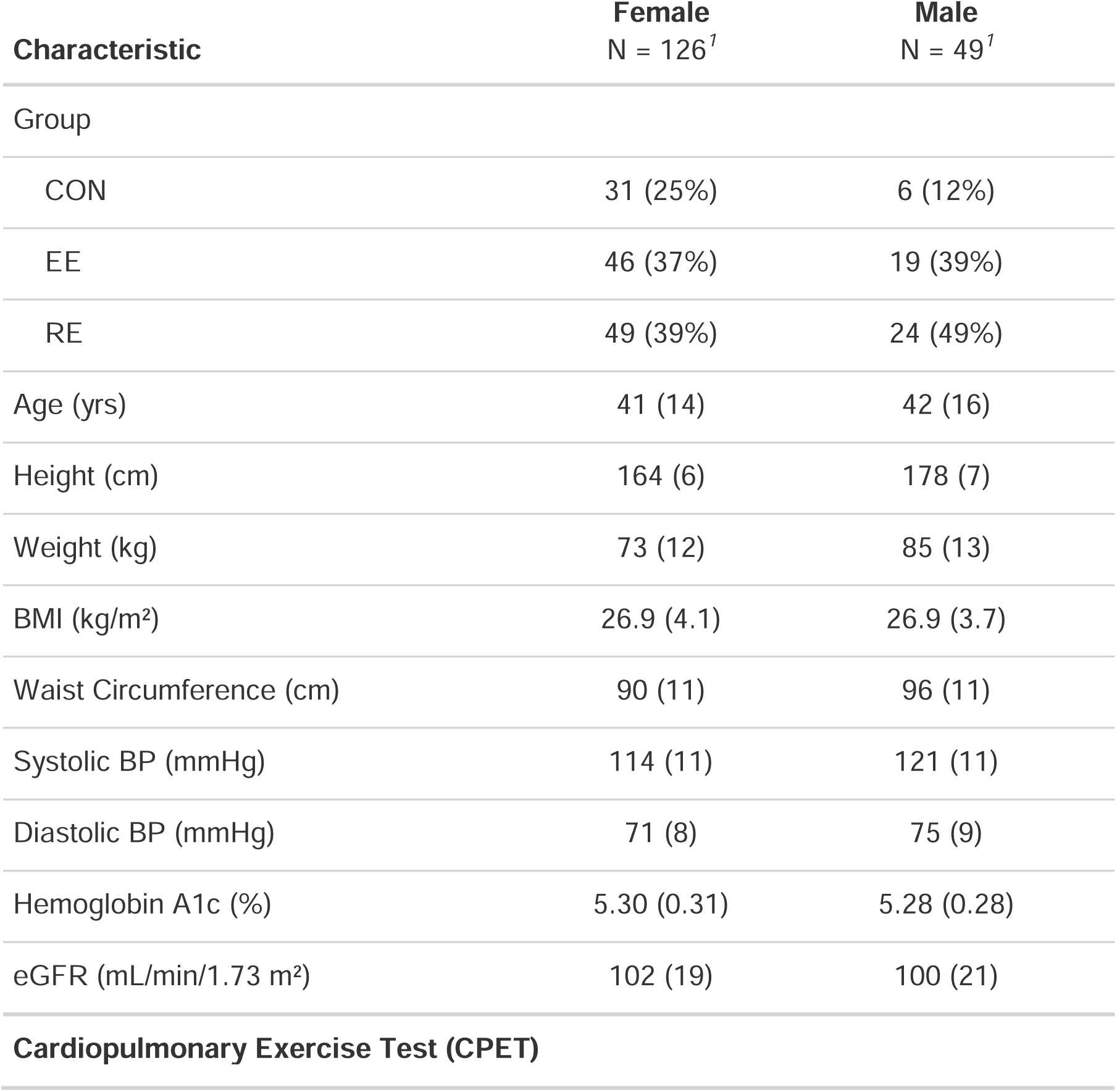
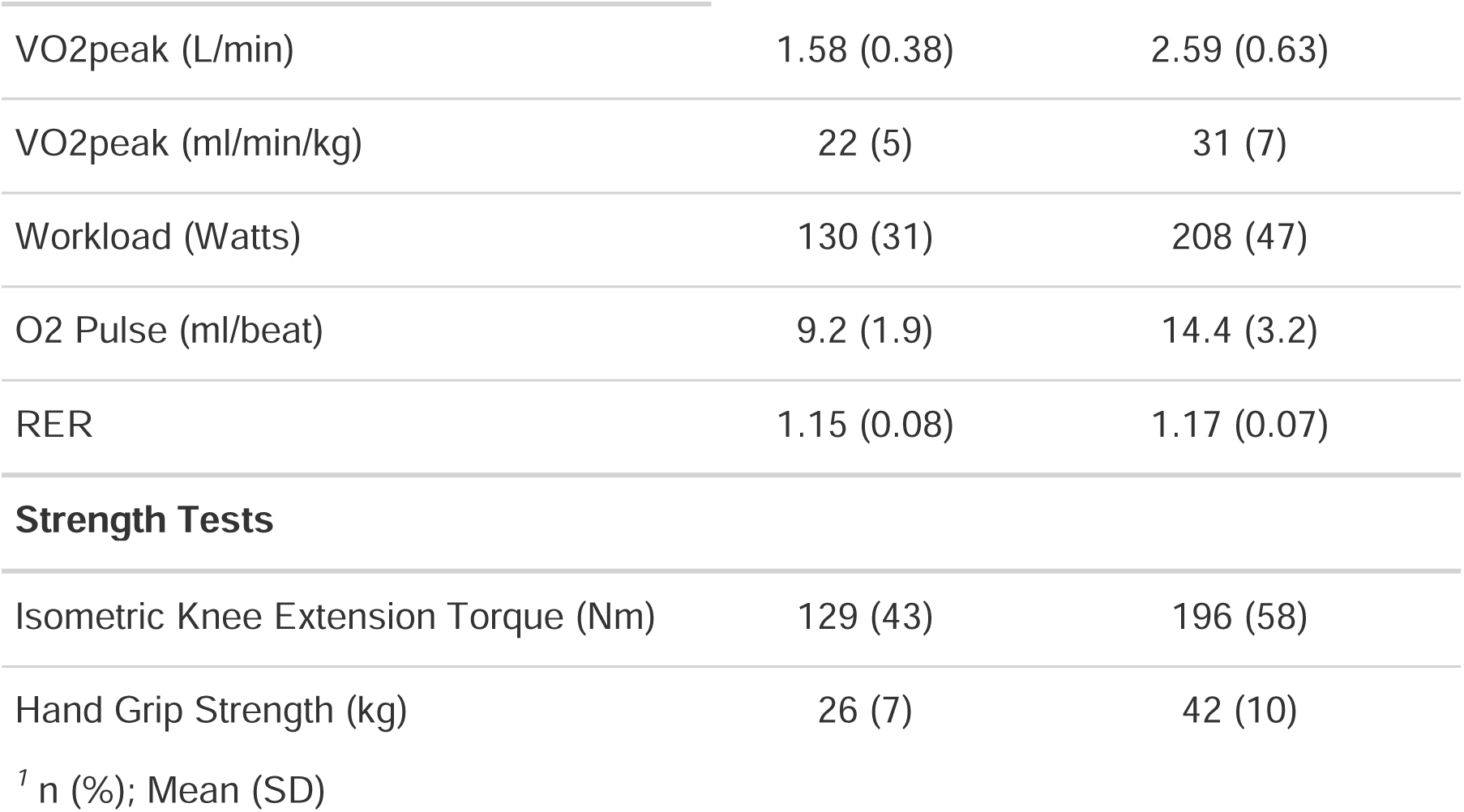
Clinical Characteristics of the MoTrPAC Pre-COVID Blood Cohort by Sex.

| Characteristic | Female<br>N = 126<sup>1</sup> | Male<br>N = 49<sup>1</sup> |
| --- | --- | --- |
| Group |  |  |
| CON | 31 (25%) | 6 (12%) |
| EE | 46 (37%) | 19 (39%) |
| RE | 49 (39%) | 24 (49%) |
| Age (yrs) | 41 (14) | 42 (16) |
| Height (cm) | 164 (6) | 178 (7) |
| Weight (kg) | 73 (12) | 85 (13) |
| BMI (kg/m<sup>2</sup>) | 26.9 (4.1) | 26.9 (3.7) |
| Waist Circumference (cm) | 90 (11) | 96 (11) |
| Systolic BP (mmHg) | 114 (11) | 121 (11) |
| Diastolic BP (mmHg) | 71 (8) | 75 (9) |
| Hemoglobin A1c (%) | 5.30 (0.31) | 5.28 (0.28) |
| eGFR (mL/min/1.73 m<sup>2</sup>) | 102 (19) | 100 (21) |
| **Cardiopulmonary Exercise Test (CPET)** |  |  |
| VO2peak (L/min) | 1.58 (0.38) | 2.59 (0.63) |
| VO2peak (ml/min/kg) | 22 (5) | 31 (7) |
| Workload (Watts) | 130 (31) | 208 (47) |
| O2 Pulse (ml/beat) | 9.2 (1.9) | 14.4 (3.2) |
| RER | 1.15 (0.08) | 1.17 (0.07) |
| Strength Tests |  |  |
| Isometric Knee Extension Torque (Nm) | 129 (43) | 196 (58) |
| Hand Grip Strength (kg) | 26 (7) | 42 (10) |

<sup>1</sup> n (%); Mean (SD)

**Table S2.** Clinical Characteristics of the MoTrPAC Pre-COVID Blood Cohort by Biochemical Profiling Platfor.

| Characteristic | Transcriptomics |  |  |  | Proteomics |  |  |  | Metabolomics |  |  |  |
| --- | --- | --- | --- | --- | --- | --- | --- | --- | --- | --- | --- | --- |
|  | Overall<br>N=173<sup>1</sup> | CON<br>N=37<sup>1</sup> | EE<br>N=64<sup>1</sup> | RE<br>N=72<sup>1</sup> | Overall<br>N=44<sup>1</sup> | CON<br>N=12<sup>1</sup> | EE<br>N=14<sup>1</sup> | RE<br>N=18<sup>1</sup> | Overall<br>N=175<sup>1</sup> | CON<br>N=37<sup>1</sup> | EE<br>N=65<sup>1</sup> | RE<br>N=73<sup>1</sup> |
| Sex |  |  |  |  |  |  |  |  |  |  |  |  |
| Female | 125<br>(72%) | 31<br>(84%) | 46<br>(72%) | 48<br>(67%) | 33<br>(75%) | 9<br>(75%) | 11<br>(79%) | 13<br>(72%) | 126<br>(72%) | 31<br>(84%) | 46<br>(71%) | 49<br>(67%) |
| Male | 48<br>(28%) | 6<br>(16%) | 18<br>(28%) | 24<br>(33%) | 11<br>(25%) | 3<br>(25%) | 3<br>(21%) | 5<br>(28%) | 49<br>(28%) | 6<br>(16%) | 19<br>(29%) | 24<br>(33%) |
| Age (yrs) | 41 (15) | 42<br>(15) | 41<br>(14) | 40<br>(15) | 43 (16) | 52<br>(13) | 40<br>(18) | 39<br>(15) | 41 (15) | 42<br>(15) | 42<br>(14) | 40<br>(15) |
| Height (cm) | 168 (9) | 167<br>(8) | 168<br>(9) | 169<br>(10) | 166<br>(10) | 166<br>(9) | 166<br>(11) | 166<br>(10) | 168 (9) | 167<br>(8) | 168<br>(9) | 168<br>(10) |
| Weight (kg) | 76 (14) | 73<br>(13) | 76<br>(13) | 77<br>(14) | 73<br>(12) | 71<br>(11) | 75<br>(15) | 73<br>(10) | 76<br>(14) | 73<br>(13) | 76<br>(13) | 77<br>(14) |
| BMI (kg/m²) | 26.9<br>(4.0) | 26.2<br>(3.9) | 27.0<br>(4.0) | 27.1<br>(4.0) | 26.6<br>(3.5) | 25.8<br>(3.2) | 27.4<br>(4.1) | 26.6<br>(3.2) | 26.9<br>(4.0) | 26.2<br>(3.9) | 27.0<br>(4.0) | 27.2<br>(4.0) |
| Waist Circumference (cm) | 92<br>(12) | 89<br>(13) | 92<br>(11) | 93<br>(12) | 91<br>(10) | 88<br>(11) | 92<br>(12) | 92<br>(8) | 92<br>(12) | 89<br>(13) | 93<br>(11) | 93<br>(12) |
| Systolic BP (mmHg) | 116<br>(12) | 114<br>(11) | 117<br>(11) | 116<br>(12) | 117<br>(11) | 119<br>(13) | 120<br>(10) | 113<br>(11) | 116<br>(12) | 114<br>(11) | 117<br>(11) | 116<br>(12) |
| Diastolic BP (mmHg) | 72 (8) | 72 (8) | 73 (9) | 72 (8) | 71 (8) | 73 (9) | 74 (7) | 69 (8) | 72 (8) | 72 (8) | 73 (9) | 72 (8) |
| Hemoglobin A1c (%) | 5.29<br>(0.30) | 5.29<br>(0.21) | 5.28<br>(0.31) | 5.31<br>(0.32) | 5.33<br>(0.29) | 5.33<br>(0.18) | 5.31<br>(0.37) | 5.35<br>(0.28) | 5.30<br>(0.30) | 5.29<br>(0.21) | 5.29<br>(0.32) | 5.31<br>(0.32) |
| eGFR (mL/min/1.73 m²) | 101<br>(20) | 99<br>(19) | 103<br>(18) | 101<br>(22) | 101<br>(18) | 94<br>(18) | 107<br>(19) | 100<br>(17) | 101<br>(20) | 99<br>(19) | 103<br>(18) | 101<br>(22) |
| VO2peak<br>(L/min) | 1.86<br>(0.65) | 1.73<br>(0.56) | 1.90<br>(0.69) | 1.90<br>(0.66) | 1.82<br>(0.73) | 1.64<br>(0.53) | 1.92<br>(0.79) | 1.86<br>(0.80) | 1.86<br>(0.65) | 1.73<br>(0.56) | 1.90<br>(0.68) | 1.89<br>(0.65) |
| VO2peak<br>(ml/min/kg) | 25 (7) | 24 (6) | 25 (7) | 25 (7) | 25 (8) | 23 (6) | 25 (9) | 25 (9) | 24 (7) | 24 (6) | 25 (7) | 25 (7) |
| Workload<br>(Watts) | 152<br>(51) | 142<br>(40) | 156<br>(54) | 154<br>(53) | 148<br>(59) | 136<br>(42) | 157<br>(63) | 149<br>(68) | 152<br>(50) | 142<br>(40) | 155<br>(53) | 153<br>(52) |
| O2 Pulse<br>(ml/beat) | 10.7<br>(3.3) | 10.0<br>(2.8) | 10.9<br>(3.5) | 10.8<br>(3.4) | 10.4<br>(3.6) | 9.9<br>(3.1) | 10.8<br>(3.8) | 10.5<br>(3.9) | 10.6<br>(3.3) | 10.0<br>(2.8) | 10.9<br>(3.5) | 10.8<br>(3.4) |
| RER | 1.16<br>(0.08) | 1.16<br>(0.08) | 1.17<br>(0.07) | 1.15<br>(0.08) | 1.15<br>(0.07) | 1.18<br>(0.09) | 1.15<br>(0.05) | 1.12<br>(0.07) | 1.16<br>(0.08) | 1.16<br>(0.08) | 1.17<br>(0.07) | 1.15<br>(0.08) |
| Isometric Knee Extension Torque (Nm) | 147<br>(57) | 137<br>(53) | 143<br>(53) | 156<br>(60) | 145<br>(44) | 148<br>(39) | 145<br>(53) | 141<br>(42) | 148<br>(57) | 137<br>(53) | 145<br>(54) | 156<br>(60) |
| Hand Grip Strength (kg) | 30<br>(11) | 29<br>(8) | 30<br>(11) | 31<br>(12) | 28<br>(12) | 29<br>(11) | 28<br>(12) | 27<br>(13) | 30<br>(11) | 29<br>(8) | 30<br>(11) | 31<br>(12) |

n (%); Mean (SD)

